# Clustering of Ca_V_1.3 L-type calcium channels by Shank3

**DOI:** 10.1101/2022.10.21.513252

**Authors:** Qian Yang, Tyler L. Perfitt, Juliana Quay, Lan Hu, Roger J. Colbran

**Author notes:** Address correspondence to: Roger J. Colbran; Rm. 702 Light Hall, Vanderbilt University School of Medicine, Nashville, TN 37232-0615 (.).

## Abstract

Clustering of neuronal L-type voltage-gated Ca^2+^ channels (LTCC) in the plasma membrane is increasingly implicated in creating highly localized Ca^2+^ signaling nanodomains. For example, LTCC activation can increase phosphorylation of the nuclear CREB transcription factor by increasing Ca^2+^ concentrations within a nanodomain close to the channel, without requiring bulk Ca^2+^ increases in the cytosol or nucleus. However, the molecular basis for LTCC clustering is poorly understood. The postsynaptic scaffolding protein Shank3 specifically associates with one of the major neuronal LTCCs, the Ca_V_1.3 calcium channel, and is required for optimal LTCC-dependent excitation-transcription coupling. Here, we co-expressed Ca_V_1.3 α1 subunits with two distinct epitope-tags with or without Shank3 in HEK cells. Co-immunoprecipitation studies using the cell lysates revealed that Shank3 can assemble multiple Ca_V_1.3 α1 subunits in a complex under basal conditions. Moreover, Ca_V_1.3 LTCC complex formation was facilitated by Ca_V_β subunits (β3 and β2a), which also interact with Shank3. Shank3 interactions with Ca_V_1.3 LTCCs and multimeric Ca_V_1.3 LTCC complex assembly were disrupted following addition of Ca^2+^ and calmodulin (Ca^2+^/CaM) to cell lysates, perhaps simulating conditions within an activated Ca_V_1.3 LTCC nanodomain. In intact HEK293T cells, co-expression of Shank3 enhanced the intensity of membrane-localized Ca_V_1.3 LTCC clusters under basal conditions, but not after Ca^2+^channel activation. Live cell imaging studies also revealed that Ca^2+^ influx through LTCCs disassociated Shank3 from Ca_V_1.3 LTCCs clusters and reduced the Ca_V_1.3 cluster intensity. Deletion of the PDZ domain from Shank3 prevented both binding to Ca_V_1.3 and the changes in multimeric Ca_V_1.3 LTCC complex assembly in vitro and in HEK293 cells. Finally, we found that shRNA knock-down of Shank3 expression in cultured rat primary hippocampal neurons reduced the intensity of surface-localized Ca_V_1.3 LTCC clusters in dendrites. Taken together, our findings reveal a novel molecular mechanism contributing to neuronal LTCC clustering under basal conditions.

## Introduction

Voltage-gated L-type calcium channels (LTCCs) are widely expressed in the central nervous system, endocrine cells, atrial myocytes, and cardiac pacemaker cells, and regulate numerous physiological processes (Catterall, 2011; Striessnig & Koschak, 2008). Clustering of the major neuronal LTCC subtypes, Ca_V_1.2 and Ca_V_1.3, amplifies Ca^2+^ influx in local Ca^2+^ nanodomains (Dixon et al., 2012; Moreno et al., 2016; Navedo & Santana, 2013) that can be sufficient to initiate some downstream pathways, without requiring Ca^2+^ increases in the bulk cytosol or nucleus (Deisseroth et al., 1996; Stern, 1992; Tadross et al., 2013). Although the importance of LTCC clustering in creating these Ca^2+^ nanodomains has been recognized, the molecular basis for cluster formation remains poorly understood.

LTCCs are comprised of a pore-forming α1 subunit (Ca_V_1.1-1.4) that co-assembles with auxiliary Ca_V_β, Ca_V_α2δ and Ca_V_γ subunits (Simms & Zamponi, 2014). The C-terminal domains of Ca_V_1.2 and Ca_V_1.3 α1 subunits play an important role in modulating LTCC cell surface expression and downstream signaling. For example, deletion of the C-terminal PDZ domain-interacting motif from Ca_V_1.2 or Ca_V_1.3 interferes with excitation-transcription (E-T) coupling (Weick et al., 2003; Zhang et al., 2005). Alternative mRNA splicing gives rise to long and short forms of the Ca_V_1.3 α1 subunit C terminal domain (Ca_V_1.3_42_ or Ca_V_1.3_L;_ Ca_V_1.3_42A_ or Ca_V_1.3_S_; Ca_V_1.3_43S_), which functionally alter voltage- and Ca^2+^-dependent gating properties (Bock et al., 2011; Hui et al., 1991; Moreno et al., 2016; Singh et al., 2008; Tan et al., 2011). Scaffolding proteins containing PDZ domains, such as Shank3, densin, and erbin interact with the C-terminal PDZ domain-interacting motif of Ca_V_1.3_L_, but not Ca_V_1.2 or Ca_V_1.3_S_, to differentially modulate the levels and pattern of cell surface Ca_V_1.3_L_ expression and Ca_V_1.3_L_ activity (Calin-Jageman et al., 2007; Jenkins et al., 2010; Stanika et al., 2016; Zhang et al., 2005). However, Ca_V_1.3_L_ and Ca_V_1.3_S_ were reported to form similar clusters in the plasma membrane that were estimated to contain an average of 8 α1 subunits in neurons (Moreno et al., 2016). Interestingly, calmodulin (CaM) binds to preIQ and IQ motifs in the C-terminal domain to facilitate cooperative channel opening of Ca_V_1.3_S_, but not Ca_V_1.3_L_, and Ca^2+^ influx (Moreno et al., 2016). Collectively these findings suggest an important role for the α1 subunit C-terminal domain in regulating LTCC activity and surface expression, as well as E-T coupling.

Of the PDZ-domain containing proteins that bind to the C-terminal domain of Ca_V_1.3_L_, Shank3 has been most intensively studied, in part because it is a multi-domain postsynaptic scaffolding protein strongly linked to multiple neuropsychiatric disorders. Previous studies found that Shank3 facilitates synaptic Ca_V_1.3_L_ surface expression (Zhang et al., 2006; Zhang et al., 2005) and is a dose-dependent regulator of calcium currents (Pym et al., 2017). In addition, Shank3 is required for normal downstream LTCC signaling to the nucleus (Perfitt et al., 2020; Pym et al., 2017; Zhang et al., 2006; Zhang et al., 2005). Although the C-terminal SAM domains of Shank3 have been shown to mediate “tail-to-tail” multimerization (Sheng & Kim, 2000), potentially facilitating the assembly of larger multi-protein complexes, the role of Shank3 in Ca_V_1.3 LTCCs clustering is poorly understood.

Here, we show that Shank3 facilitates the assembly of complexes containing multiple Ca_V_1.3_L_ α1 subunits *in vitro* and on the surface of intact HEK293 cells, and that clustering is further enhanced by Ca_V_β subunits. This robust Shank3-dependent clustering under basal conditions is disrupted by the addition of Ca^2+^/CaM in vitro, or by LTCC activation in HEK cells. Moreover, we found that knock-down of Shank3 expression disrupted basal cell surface Ca_V_1.3 clustering in the dendrites of cultured rat hippocampal neurons. Taken together, our data indicate that Shank3 assembles Ca_V_1.3 LTCCs clusters under basal conditions, which may be important for downstream Ca^2+^ signaling.

### Experimental procedures

#### DNA constructs

Original sources of previously described DNA constructs are provided in the Key Resources Table. The Shank3 construct containing a deletion of the PDZ domain (GFP-Shank3-ΔPDZ) was generated by in-frame PCR deletion of the entire 270 bp region encoding ^572^Iso-Val^661^ from the parent GFP-Shank3 construct. Sequences of all constructs were confirmed by DNA sequencing.

#### Culture and transfection of HEK cells

HEK293 and HEK293T cells were grown at 37°C and 5% CO_2_ in DMEM plus 10% (v/v) fetal bovine serum (Gibco), 1% (w/v) penicillin/streptomycin (Gibco), 1% (v/v) MEM non-essential amino acid solution (Sigma, catalog no. RNBK3078), and 1% GlutaMAX (Gibco, catalog no. 2248970). Cells were co-transfected at ∼70% confluence using Lipofectamine 2000 (Invitrogen). HA-Ca_V_1.3_L_ and α2δ with or without mCherry-Ca_V_1.3_L_ were co-expressed with the empty Flag vector or vectors encoding FLAG-β3 or -β2a together with GFP or GFP-Shank3 (WT or with PDZ deletion), as indicated (ratio of α1: α2δ: β: Shank3 was 3:1:1:1.5). For co-immunoprecipitation and GST pulldown experiments, HEK293T cells were transfected using a total of ≤10 μg of DNA per 10-cm culture dish (Corning, catalog no. 430167). The medium was completely changed 24 h after transfection and cells were harvested after 48 h for co-immunoprecipitation or GST pulldown assay. For immunostaining and live-cell imaging, HEK293 cells, which generally express lower levels of recombinant proteins, were transfected with ≤2 μg of DNA per well of a 6-well plate. Cells were re-plated at low density 24 h after transfection into a 24-well plate with 12×12 mm^2^ coverslips (Fisher Scientific, catalog no. 22293232) or into a 29 mm dish with 10 mm bottom well (Cellvis, catalog no. D29-10-1.5-N) for live-cell imaging. Cells were grown for another 24-48 h before treatment and fixation, or live-cell imaging.

#### Co-immunoprecipitation (Co-IP)

Transfected cells were lysed in lysis buffer (50 mM Tris-HCl, 150 mM NaCl, 1 mM EDTA, 1 mM EGTA, 1 mM DTT, 1% Nonidet P-40 (v/v), 1 mM Microcystin-LR, and protease inhibitor mixtures). Cell lysates were homogenized (15-25 strokes) with Branson Sonifier 450 (VWR SCIENTIFIC) and then were cleared by low-speed centrifugation (500 x g). The supernatant was then incubated at 4°C for 4 h with rabbit anti-HA (Cell Signaling; 1:500; Figures 2A, 4B and 5A) or 1 h with rabbit anti-GFP (Thermo Fisher Scientific; 1:1000; Figure 3A) or for 2-3 h with mouse anti-Flag M2 antibody (Sigma; 1:500: Figure 3C) and 10 μl of prewashed Dynabeads Protein A (Thermo Fisher Scientific, catalog no. 10002D; for rabbit antibodies) or Dynabeads Protein G (Thermo Fisher Scientific, catalog no. 10004D; for mouse antibodies). Where indicated, lysates were supplemented with 2 mM CaCl_2_, and 1 μM calmodulin (final concentrations) during the incubation. The beads were isolated magnetically and washed three times using lysis buffer before eluting proteins using 2X SDS-PAGE sample buffer.

**Figure 1.**
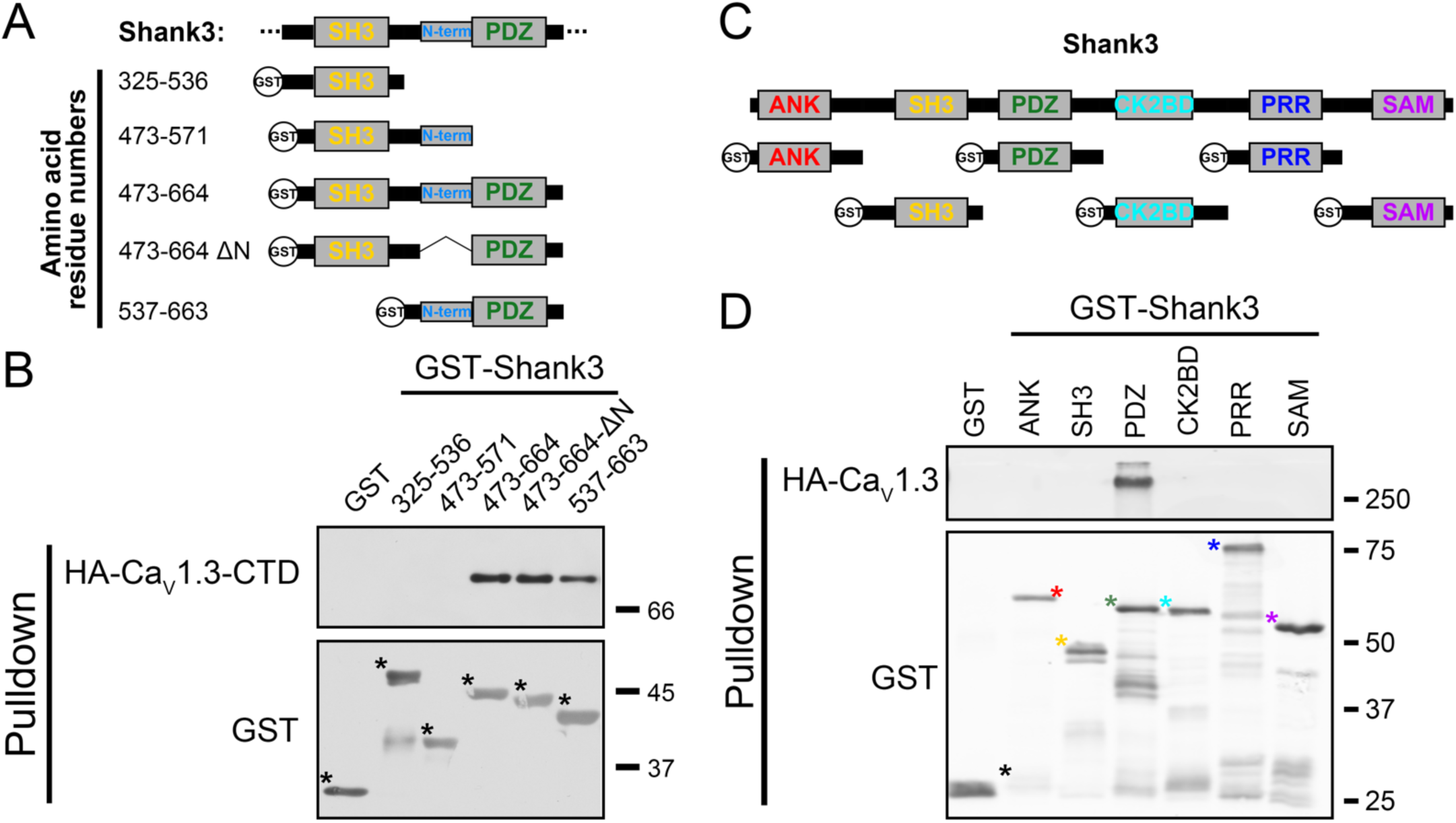
The Shank3 PDZ domain is necessary and sufficient for interaction with the CaV1.3 C-terminal domain. A) Schematic of Shank3 truncations and deletions expressed as GST fusion proteins for use in panel B, with amino acid residue numbers. The ΔN-term deletion removed residues 543-564. B) An anti-HA immunoblot (top) of glutathione agarose co-sedimentation assays revealed that HA-Ca_V_1.3-CTD binds to all GST-Shank3 proteins containing the PDZ domain but not to proteins lacking the PDZ domain. Full-length GST fusion proteins are marked with asterisks on the corresponding GST immunoblot (bottom). C) Domain structure of full-length Shank3 and six GST-Shank3 fusion proteins spanning the entire Shank3 protein used in panel D. Canonical Shank3 domains are depicted as gray boxes: ANK = ankyrin-rich repeats, aa 1-324; SH3 = Src homology 3 domain, aa 325-536; PDZ = PSD95/Dlg1/zo-1 domain, aa 537-828; CK2BD = CaMKII binding domain, aa 829-1130; PRR = proline-rich region, aa 1131-1467; SAM = Sterile alpha motif, aa 1468-1740. D) An anti-HA immunoblot (top) of a glutathione agarose co-sedimentation assay detected binding of full-length HA-Ca_V_1.3 α subunit only to the GST-Shank3-PDZ domain protein. Full-length GST fusion proteins are marked with color coded asterisks on the corresponding GST immunoblot (bottom). Panels B and D are representative of three independent biological replicates.

**Figure 2.**
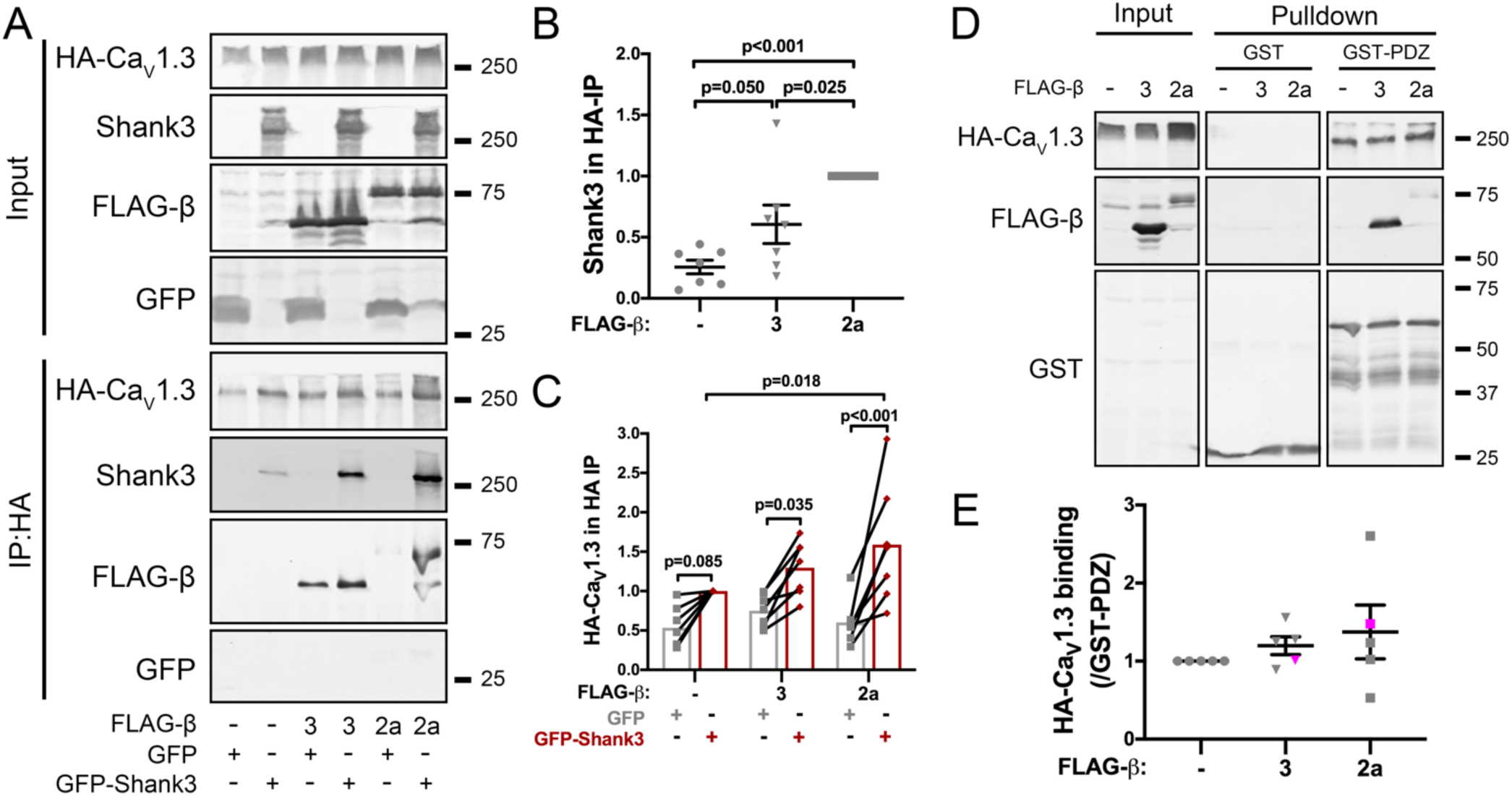
Association of GFP-Shank3 with HA-Ca_V_1.3 is facilitated by co-expression of Flag-β subunits. A) Representative immunoblots of HA, Shank3, FLAG, and GFP signals in the input (top) and anti-HA immune complexes (bottom) isolated from soluble fractions of HEK293T cells co-expressing HA-Ca_V_1.3 with GFP or GFP-Shank3, with or without FLAG-β2a or -β3 subunits, as indicated below. Quantifications of the Shank3 (B) and HA-Ca_V_1.3 (C) signals in HA-IPs: mean ± SEM, n = 7 independent transfections. B: One-way ANOVA followed by Tukey’s post hoc test. C: Two-way ANOVA followed by Sidak’s post hoc test when comparing GFP to GFP-Shank3 or by Turkey’s post hoc test when comparing between no β, β3, and β2a. D) Representative immunoblots of HA, FLAG, and GST signals in soluble fractions of HEK293T cells co-expressing HA-Ca_V_1.3, α2δ, with or without Flag-β2a or -β3 subunits (input) and in glutathione agarose co-sedimentation assays following incubation with the GST-Shank3-PDZ domain (2 µg). E) Quantification of HA-Ca_V_1.3 signals in GST complexes obtained from 5 independent transfected cell samples incubated with two different GST-Shank3-PDZ domain constructs containing either residues 537-828 (as in Fig. 1D: 1 replicate, magenta symbols) or residues 572-691 (4 replicates, gray symbols). Mean ± SEM, n = 5. No significant differences between groups by one-way ANOVA.

**Figure 3.**
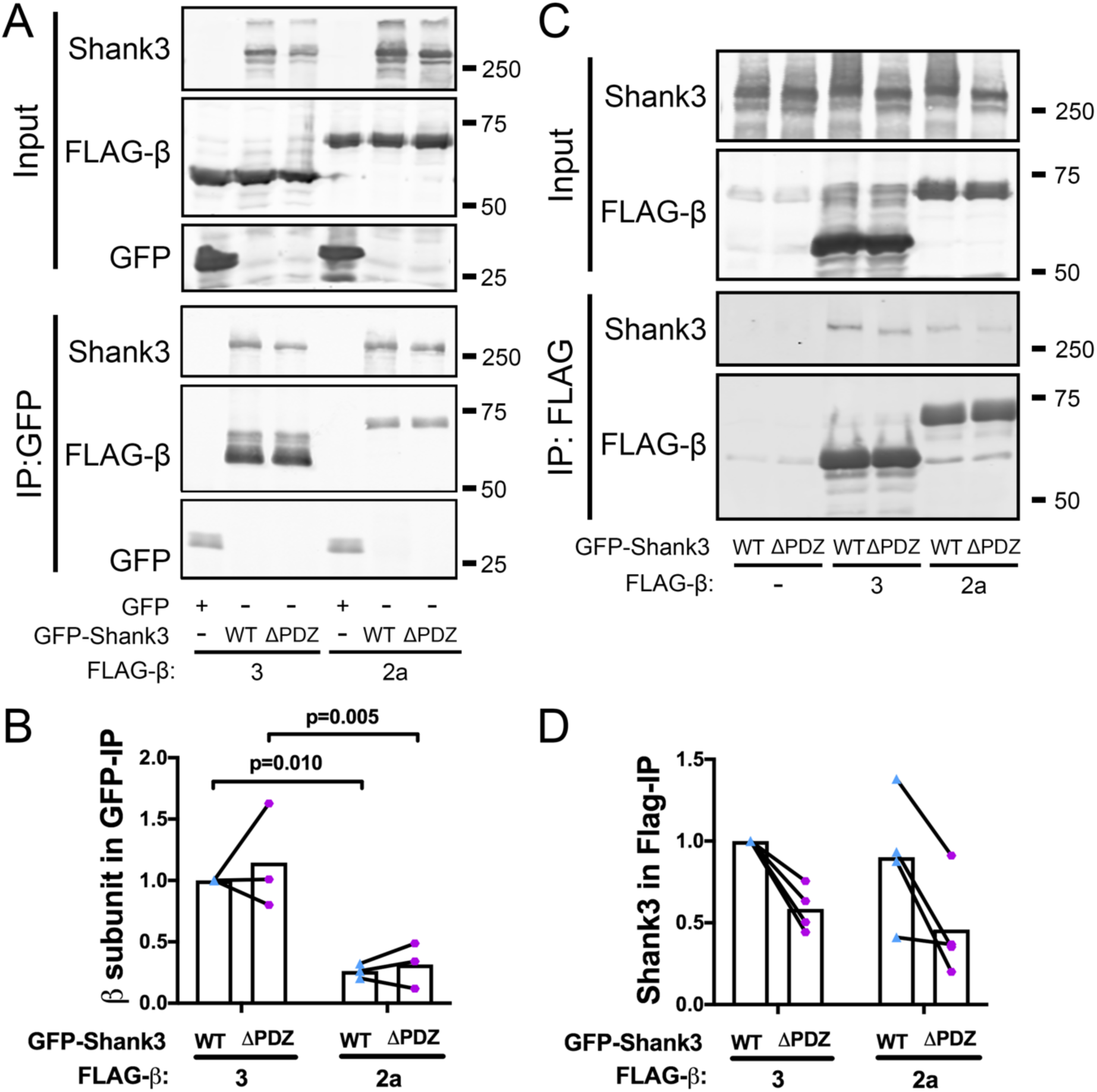
Association of Flag-β subunits with GFP-Shank3. A) Representative Shank3, Flag and GFP immunoblots of soluble fractions (Input) of HEK293T cells co-expressing GFP (control) or GFP-Shank3 (WT or ΔPDZ) with or without FLAG-β3 or -β2a subunits, and corresponding isolated anti-GFP immune complexes. B) Quantification of FLAG-β subunit signals in GFP-Shank3 immune complexes from 3 independent transfected cell replicates. Mean ± SEM: two-way ANOVA followed by Sidak’s post hoc test. C) Representative Shank3 and Flag immunoblots of inputs and anti-FLAG immune complexes isolated from HEK293T cells expressing GFP-Shank3 (WT or ΔPDZ) with or without FLAG-β3 or β2a. D) Quantification of GFP-Shank3 signals in FLAG-β immune complexes from 4 independent transfected cell replicates. Mean ± SEM: two-way ANOVA followed by Sidak’s post hoc test.

**Figure 4.**
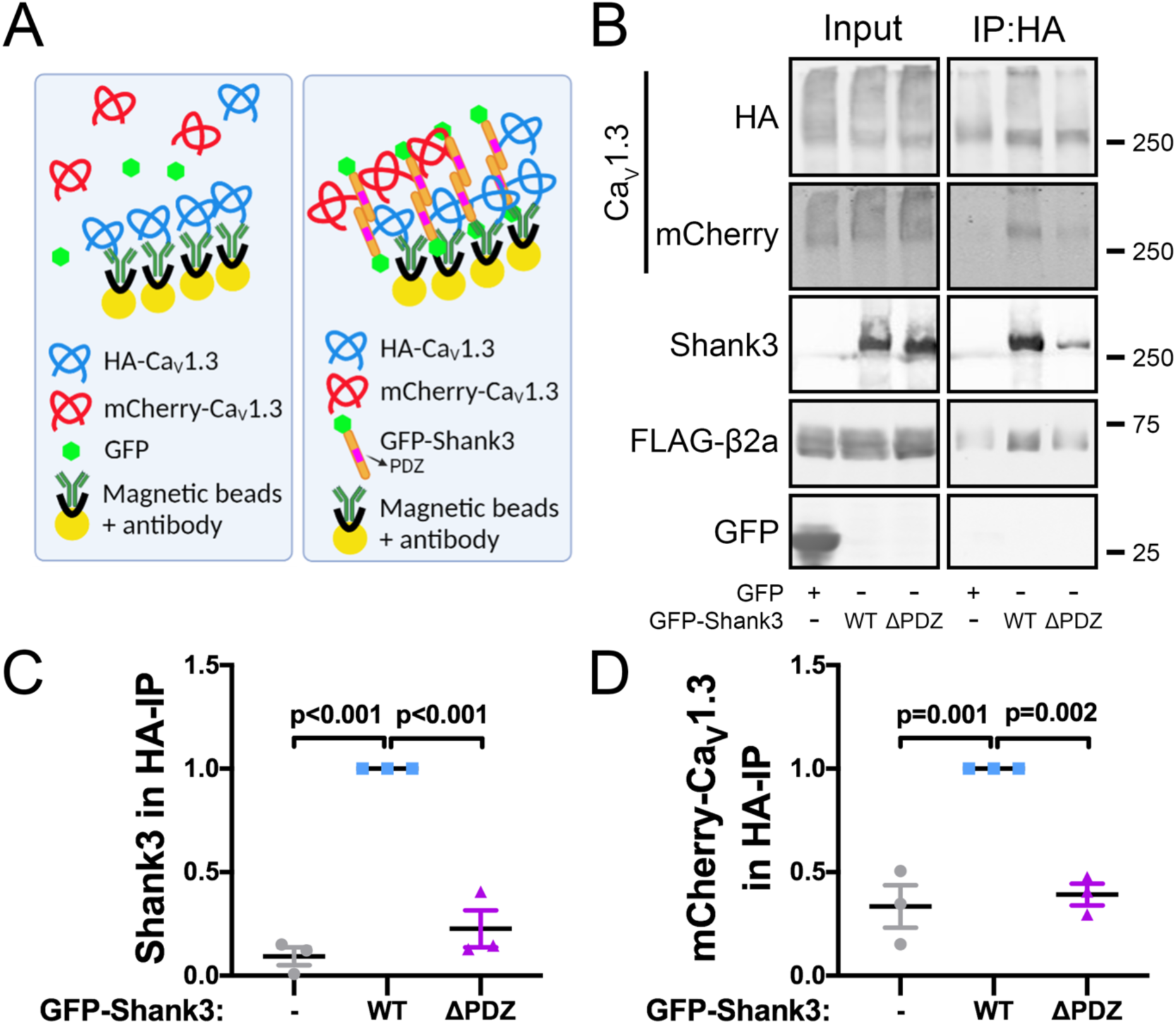
Assembly of multi-Ca_V_1.3 LTCC complexes requires the Shank3 PDZ domain. A) Schematic of experimental design to test the hypothesis that Shank3 mediates the assembly of complexes containing multiple Ca_V_1.3 α1 subunits. In the presence of GFP (left), mCherry-Ca_V_1.3 cannot associate with anti-HA IPs. PDZ domains in GFP-Shank3 dimers associate with both HA- and mCherry-Ca_V_1.3, mediating the isolation of both GFP-Shank3 and mCherry-Ca_V_1.3 by anti-HA IP (right). B) Representative immunoblots for HA- and mCherry-Ca_V_1.3, FLAG-β2a, Shank3 and GFP in the inputs and anti-HA immunoprecipitations (IPs) from soluble fractions of HEK293T cells co-expressing HA- and mCherry-tagged Ca_V_1.3 and FLAG-β2a with either GFP or GFP-Shank3 (WT or ΔPDZ). C) Quantification of GFP/GFP-Shank3 (WT or ΔPDZ) signals in HA-IPs, normalized to HA-Ca_V_1.3 signal, from three independent transfections. D) Quantification of mCherry-Ca_V_1.3 signals in HA-IPs, normalized to HA-Ca_V_1.3 signal, from three independent transfections. Mean ± SEM: One-way ANOVA followed by Tukey’s post hoc test.

**Figure 5.**
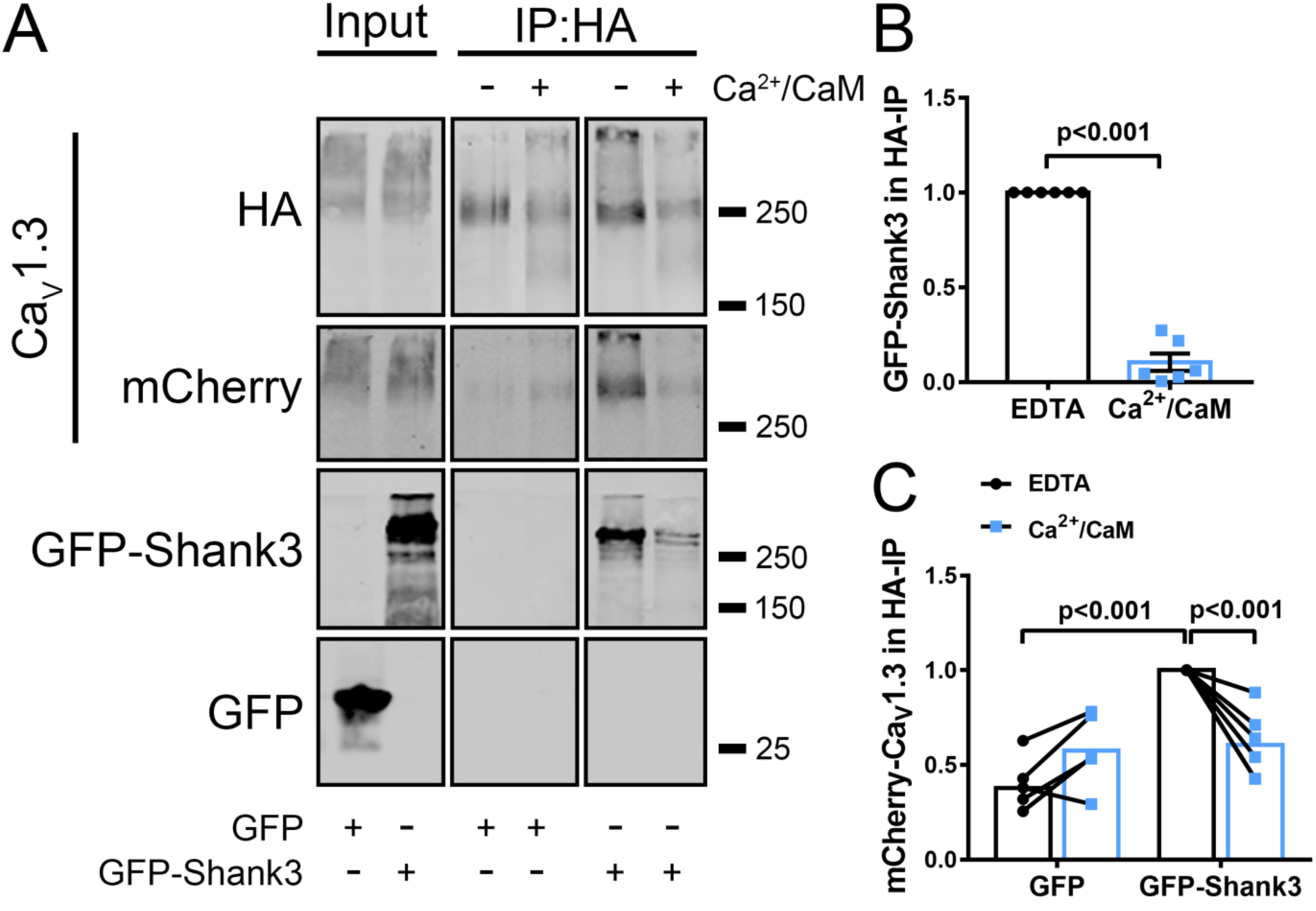
Assembly of multi-Ca_V_1.3 LTCC complexes by Shank3 is suppressed by Ca^2+^/CaM. A) Representative immunoblots for HA- and mCherry-Ca_V_1.3, and GFP in inputs and anti-HA immunoprecipitations (IPs) from soluble fractions of HEK293T cells co-expressing HA- and mCherry-tagged Ca_V_1.3 and FLAG-β2a with either GFP or GFP-Shank3 without (EDTA) or with Ca^2+^/CaM addition. B) Quantification of GFP-Shank3 in HA-IPs from six independent transfections normalized to the EDTA control; analyzed using a one-sample t-test. C) Quantification of mCherry-Ca_V_1.3 in HA-IPs from six independent transfections normalized to the EDTA/GFP-Shank3 control; analyzed using a two-way ANOVA followed by Sidak’s post hoc test.

#### GST pulldown

GST-Shank3 constructs were created, expressed, and purified as previously described (Perfitt et al., 2020). Transfected cell supernatants (see above) were incubated at 4°C with ∼150 nM of the indicated full-length GST fusion proteins (or GST control) and 10 μl prewashed glutathione magnetic beads for 1-2 h. Beads were then separated magnetically and washed three times with GST pulldown buffer (50 mM Tris-HCl pH 7.5; 200 mM NaCl; 1% (v/v) Triton X-100). GST protein complexes were eluted by incubation with 40 μl of 20 mM glutathione (pH 8.0) (Sigma) in GST pulldown buffer at 4°C for 10 min.

#### Western blot analysis

Samples were resolved on 10% (Figures 1, 2, and 3A; Figure S1) or 7.5% (Figures 3C, 4 and 5) SDS-PAGE gels and transferred to nitrocellulose membrane (Protran, Camp Hill, PA). The membrane was blocked in blotting buffer containing 5% nonfat dry milk, 0.1% (v/v) Tween-20, in Tris-buffered saline (20 mM Tris, 136 mM NaCl) at pH 7.4 for 1 h at room temperature. The membrane was incubated at 4°C with primary antibody (see dilutions above) in blotting buffer overnight. After washing with washing buffer (0.1% (v/v) Tween 20 in Tris-buffered saline) two times (10 min/time), membranes were incubated with IR dye-conjugated (all replicates of GST-pulldown experiments and most replicates of Co-IP experiments) or HRP-conjugated secondary antibody (three replicates of the Co-IP experiments in Figure 5) for 1 h at room temperature and washed again before development. Secondary antibodies conjugated to infrared dyes (LI-COR Biosciences) were used for development with an Odyssey system (LI-COR Biosciences). Blots incubated with HRP-conjugated secondary antibodies were incubated with the Western Lightening Plus-ECL, enhanced chemiluminescent substrate (PerkinElmer, Waltham, MA) and visualized using Premium X-ray Film (Phenix Research Products, Candler, NC) exposed to be in the linear response range. Images were quantified using Fiji software (RRID: SCR_003070). Background signals in equivalent areas from the negative control lanes were subtracted from signals in the experimental lanes. Similar results were obtained when the same samples were analyzed in parallel using ECL and Odyssey-based methods in some studies.

#### HEK cell stimulation, immunocytochemistry

BayK 8644 (BayK) was prepared as a 50 mM stock solution in DMSO. For the experiment in Figure 8, HEK293 cells were incubated in HEPES buffer (150 mM NaCl, 5 mM KCl, 2 mM MgCl_2_, 10 mM HEPES pH 7.4, 10 mM Glucose) for 10 min. Cells from different wells were then switched to each of these conditions: HEPES buffer + DMSO (0.02% v/v), HEPES buffer + BayK (10 μM), 2.5 mM Ca^2+^ buffer (HEPES buffer + 2.5 mM CaCl_2_) + DMSO, or 2.5 mM Ca^2+^ buffer + BayK for 10-15 min each. After a further 10-15 min, cells were fixed using ice-cold 4% paraformaldehyde containing 4% sucrose in 0.1 M Phosphate Buffer (pH 7.4) for 10 min. Cells expressing mCherry-Ca_V_1.3 and GFP or GFP-Shank3 (WT or ΔPDZ) (Figure 8) were washed three times with PBS after fixation and then mounted on slides using Prolong Gold Antifade Mountant.

**Figure 6.**
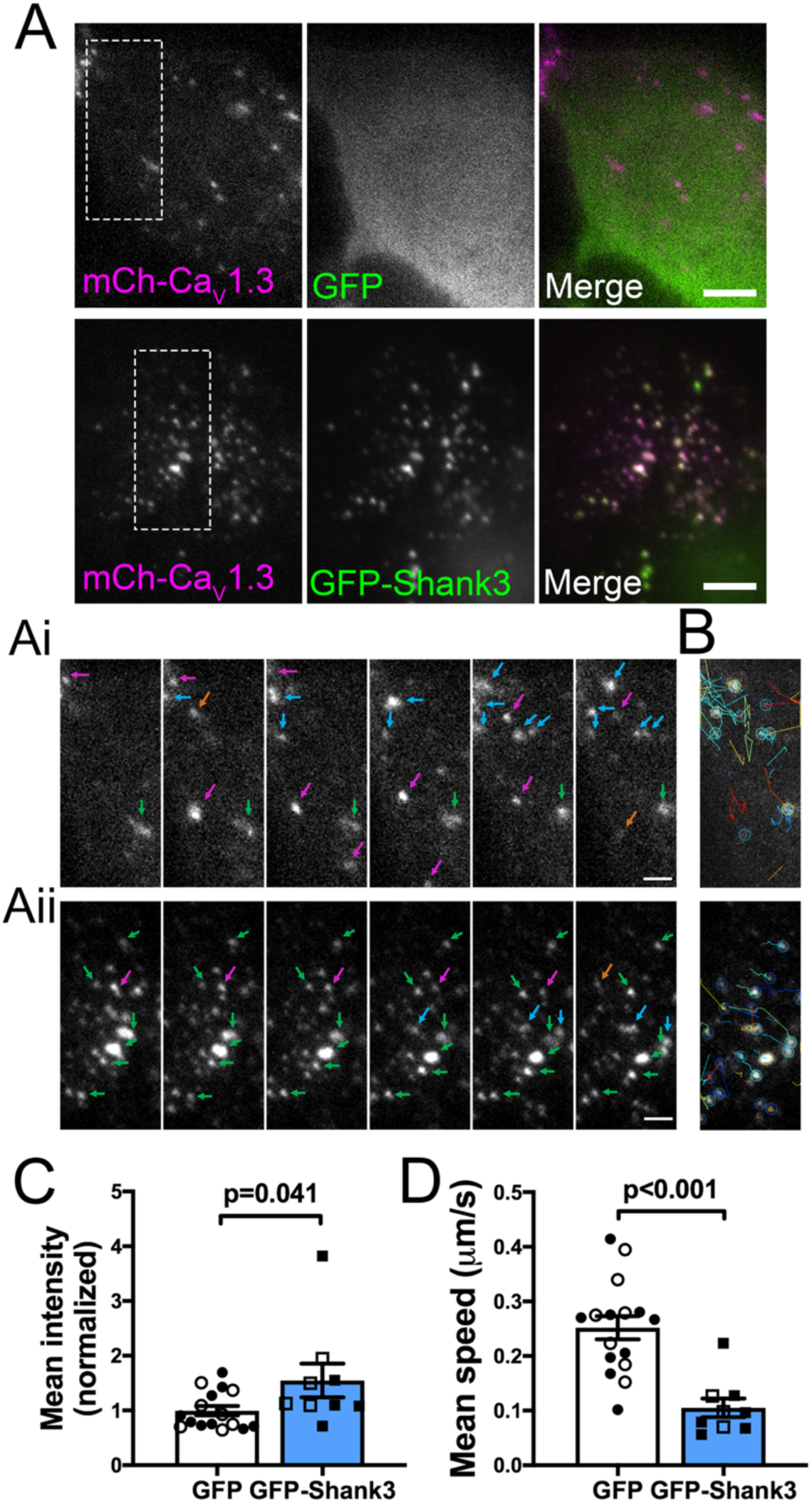
GFP-Shank3 modulates mCherry-Ca_V_1.3 dynamics in HEK293 cell plasma membranes. TIRF microscope imaging of live HEK293 cells co-expressing mCherry-Ca_V_1.3. A) Representative single channel and merged TIRF microscope images of live HEK293 cells expressing mCherry-Ca_V_1.3, FLAG-β3 and either GFP (top) or GFP-Shank3 (bottom). Enlarged time lapse mCherry images within the indicated rectangular regions of interest are shown in Ai and Aii (Supplemental Movies 1 and 2 show the entire time course). Colored arrows indicate the properties of selected mCherry puncta: Green, puncta present throughout; Red, puncta that disappear; Orange, puncta that appear transiently; Blue, puncta that appear but remain to the last time point. Scale bars, 5 μm in A and 2 μm in Ai and Aii. B) Tracking lateral movement of individual Ca_V_1.3 puncta in the plane of the TIRF image using the FIJI TrackMate plug-in, superimposed on images from the last time point in Ai and Aii. C) Quantification of the average intensity of mCherry-Ca_V_1.3 puncta. D) Quantification of the speed of lateral movement of mCherry-Ca_V_1.3 puncta (TrackMate). Data in panels C and D were collected from 16 (GFP) or 9 (GFP-Shank3) cells from 5 independent transfections. Open and solid symbols are from cells transfected with FLAG-β2a or FLAG-β3, respectively. Mean ± SEM: unpaired t-test.

**Figure 7.**
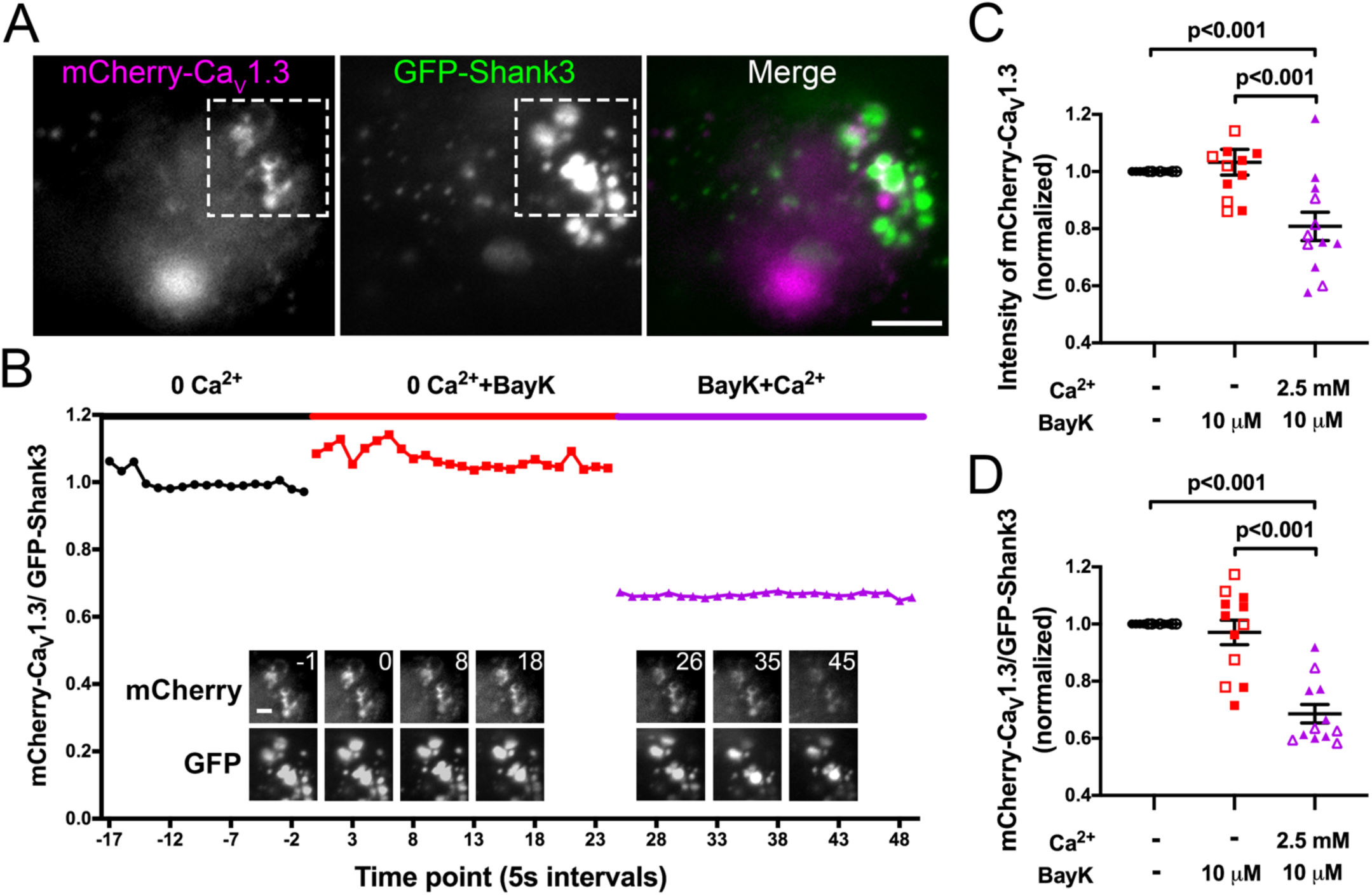
Ca^2+^ influx dissociates GFP-Shank3 from mCherry-Ca_V_1.3 in live HEK293 cells. A) Representative mCherry, GFP and merged TIRF microscope image of a live HEK293 cell co-expressing mCherry-Ca_V_1.3, FLAG-β3 and GFP-Shank3 at the start of the experiment (scale bar, 5 µm). B) The cell was imaged every 5 s for 2-3 minutes each in “no Ca^2+^” buffer, following the addition of BayK 8644 (10 µM), and following the further addition of Ca^2+^ (2.5 mM CaCl_2_). No images were collected for ∼1 min during each buffer addition. The ratio of mCherry-Ca_V_1.3 to GFP-Shank3 signal intensity in the region of interest (highlighted in panel A) was quantified at each time point. Insets show enlarged ROI images of mCherry-Ca_V_1.3 (top row) and GFP-Shank3 (bottom row) images at selected time points (scale bar, 2 μm). Supplemental Movie 3 shows all time points. C) Summary of average mCherry-Ca_V_1.3 signal intensity from all time points under each condition, normalized to the “no Ca^2+^” condition. D) Ratio of mCherry-Ca_V_1.3 to GFP-Shank3 signal intensity from all time points under each condition, normalized to the “no Ca^2+^” condition. Data in panels C and D were collected from 12 cells analyzed from six transfections (open and solid symbols indicate expressing FLAG-β2a or FLAG-β3, respectively). One-way ANOVA followed by Tukey’s post hoc test was used for comparisons.

**Figure 8.**
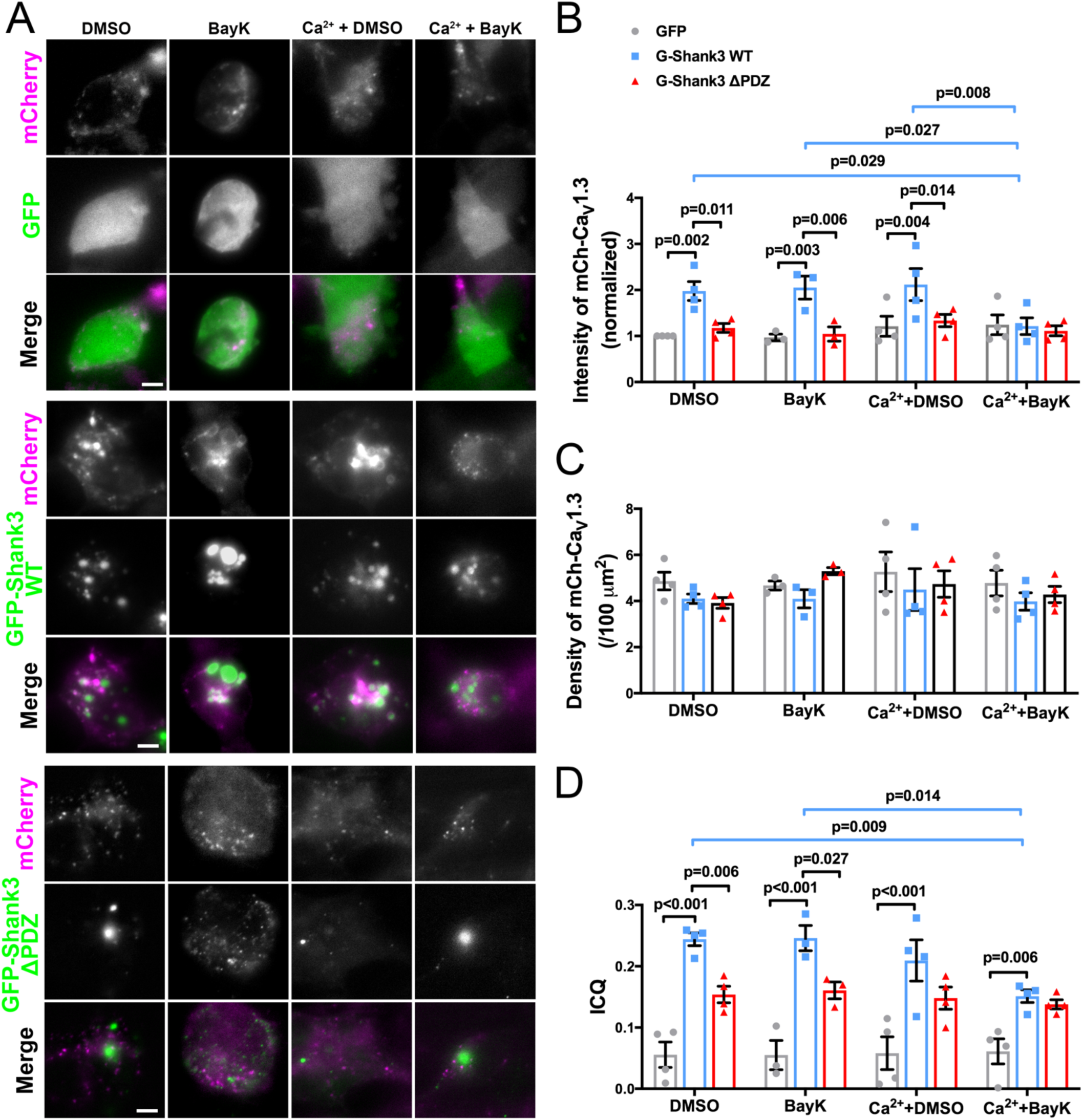
Shank3 and Ca^2+^ influx regulate mCherry-Ca_V_1.3 puncta intensity in HEK293 cell plasma membranes. A) Representative TIRF microscope images of single HEK293 cells co-expressing mCherry-Ca_V_1.3 and FLAG-β3 with either GFP or GFP-Shank3 (WT or ΔPDZ), fixed following incubation for 10-15 min in “no Ca^2+^” or Ca^2+^ buffer with vehicle (DMSO) or BayK 8644 (BayK, 10 μM), as indicated (scale bar, 5 μm). B) Quantification of mCherry-Ca_V_1.3 puncta intensity. C) Quantification of mCherry-Ca_V_1.3 puncta density. D) Intensity correlation analysis of GFP/mCherry colocalization. Panels B-D plot the mean ± SEM, with each data point representing the average of 7-15 cells per condition from 3 or 4 independent transfections. Data were compared using a two-way ANOVA followed by Tukey’s multiple comparisons test.

#### Total Internal Reflection Fluorescence (TIRF) Microscopy

All HEK cell imaging was performed using a Nikon Multi Excitation TIRF microscope with a 60x/1.49 n.a. TIRF objective (Nikon, Tokyo, Japan), Andor Xyla sCMOS camera (Andor, Belfast, UK); 405-, 488-, 561-, and 640 nm solid-state lasers (Nikon LU-N4); HS-625 high-speed emission filter wheel (Finger Lakes Instrumentation, Lima, NY); and standard filter sets. Images were acquired using NIS-Elements (Nikon) with the same exposure time of 30-100 ms for both channels and 3-5% laser power for 488 nm and 10-15% laser power of 561 nm. These parameters were kept the same for all cell imaging in the same replicate.

For the long-term time-lapse imaging of live HEK293 cells, the live tissue chamber (TOKAI HIT, Japan) with atmosphere heater, stage heater, humidity, and CO_2_ control was used. Perfect Focus (PFS) was on during the whole imaging session to hold the correct focal plane. The interval was set as 5 seconds, and the duration of each treatment (phase) was 2-3 minutes (Figure 7) or 5-10 min (Figure 6). In Figures 7, cells were first imaged in 0 Ca^2+^ HEPES buffer (see above). Image collection was paused and the buffer was changed to HEPES buffer + 10 μM BayK for the 2^nd^ imaging phase and then to 2.5 mM Ca^2+^ buffer + 10 μM BayK for the 3^rd^ imaging phase. The time gap between each phase was about one minute. The position of target cells was confirmed after each buffer change before re-starting image collection.

All images were opened and processed in Fiji software. The GFP channel and the Polygon Selection tool were used to select the region of interest (ROIs) corresponding to the outline of the cell. The background was flattened and the mCherry-Ca_V_1.3 ROIs were thresholded based on the fluorescence signal. The threshold was defined using the mean intensity of mCherry plus two-times the standard deviation. Analyze Particles was used to calculate the intensity, area, and numbers of mCherry-Ca_V_1.3 clusters above the threshold. Cluster density was calculated using the cluster number divided by ROI area. For mCherry-Ca_V_1.3 intensity analysis in images of live cells, ROIs that include at least four mCherry clusters colocalized with GFP-Shank3 were used for quantification, and Analyze Particles was applied to all-time series. For colocalization analysis, GFP and mCherry channels were automatically thresholded before calculating the intensity correlation quotient (ICQ), which quantifies co-localization from complete segregation to perfect overlap on a -0.5 to +0.5 scale, as previously described (Li et al., 2004; Perfitt et al., 2020).

#### Tracking of mCherry-Ca_V_-1.3 LTCC clusters on the cell surface

The motility of mCherry-tagged Ca_V_1.3 clusters with or without Shank3 co-expression was compared by automatic tracking using TrackMate in Fiji. LoG detector was used, and the estimated diameter of particles was set to 0.8 μm. Then, HyperStack Displayer was selected as the mode of viewer. Quality was added as a filter to rule out the background selection and spots color was set by mean intensity. Finally, Simple LAP Tracker was used, and the maximum frame gap was set to 2 (one or two missing time points were allowed while tracking). The linking and gap-closing maximum distance was adjusted individually depending on the observation of satisfactory trajectories from frame to frame by visual inspection to avoid false connections. The dynamic parameter (including track ID, displacement, duration, X, Y, Z location, and mean speed of the event being tracked) of all tracks were exported from Fiji for further analysis.

#### Primary hippocampal neuron cultures and immunocytochemistry

Dissociated hippocampal neurons were prepared from E18 Sprague Dawley rat embryos, as previously described (Shanks et al., 2010). The brains from all the available embryos were dissected and pooled to prepare the cultures, so presumably, the cultures contain an ∼50:50 mix of neurons from male and female pups. Neurons were transfected at 14 days in vitro (DIV) using Lipofectamine 2000 following the manufacturer’s directions (Thermo Fisher Scientific). sHA-Ca_V_1.3, α2δ, and FLAG-β subunit (β3 or β2a) were co-transfected with GFP-nonsense shRNA (nssh) or GFP-Shank3-shRNA or GFP-Shank3 (ratio of α1: α2δ: β: GFP was 3:1:1:1). A total of 1 μg of DNA was transfected for each well of a 12-well plate for 2-3 hours before switching back to the conditioned medium. Neurons were used for immunostaining at DIV20-21. All procedures were pre-approved by the Vanderbilt University Institutional Animal Care and Use Committee and followed the National Institutes of Health Guide for the Care and Use of Laboratory Animals.

Neurons were live-stained for surface sHA-Ca_V_1.3 labeling. Briefly, half of the conditioned medium (500 μl) was collected for secondary antibody dilution and then the anti-HA antibody (1:200) was added into the remaining medium for 15-20 min (Stanika 2016). Neurons were quickly but carefully washed using prewarmed HBSS (Gibco) 3 times after primary antibody incubation. Neurons were incubated in the conditioned medium containing secondary antibodies (1:200) for another 15-20 min at 37°C. After three quick washes with prewarmed HBSS, neurons were immediately fixed using ice-cold 4% paraformaldehyde containing 4% sucrose in 0.1 M Phosphate Buffer pH 7.4 for 3 min and -20°C methanol for 10 minutes. Neurons were washed with PBS three times, and then permeabilized with PBS containing 0.2% Triton X-100, and then incubated with blocking solution (1X PBS, 0.1% Triton X-100 (v/v), 2.5% BSA (w/v), 5% Normal Donkey Serum (w/v), 1% glycerol (v/v)) at room temperature for 1 hour. Cells were then incubated with the blocking solution containing rabbit anti-Shank3 antibody overnight at 4°C. The following day, cells were washed three times in PBS containing 0.2% Triton X-100, then incubated with the blocking solution containing secondary antibody for 1 hour at room temperature. After washing with PBS three times, cells were mounted on slides using Prolong Gold Antifade Mountant with DAPI.

#### Neuronal imaging and quantification

All neuronal imaging was performed using a 63x/1.40 Plan-APOCHROMAT oil lens as the primary objective on Zeiss LSM880 with AiryScan (Carl Zeiss Microscopy, Jena, Germany). The binocular lens was used to identify the transfected neurons based on GFP expression driven by the shRNA constructs. For whole-cell imaging, focal plane z stacks (0.3 μm steps; 1.5-2.4 μm range) were acquired. Fiji software (ImageJ, NIH) was used to merge a series of z stack images into one maximum intensity projection image.

The AiryScan module was used to maximize sensitivity and resolution for imaging surface localized sHA-Ca_V_1.3 in neurons. The scanned area was 73.51 x 73.51 μm. Images were opened and quantified in Fiji (ImageJ, NIH). The GFP channel was used to select the regions of interest (ROIs) for measuring the numbers and intensity of sHA-Ca_V_1.3 clusters. Analysis of somatic clusters was the same as in HEK cells. A 15–25 μm segment of 2-3 secondary dendrites were selected for analysis which meet criteria: (1) > 50 μm away from the soma; (2) no other crossing dendrites; (3) similar thickness. After selecting the ROI, the background was subtracted and sHA-Ca_V_1.3 ROIs were thresholded as for HEK cell analyses. Analyze Particles in Fiji was then used to measure the intensity, area, and numbers of the surface localized sHA-Ca_V_1.3 clusters. In addition, a segmented line was used to measure the length of selected dendritic segments. Dendritic cluster density in each dendritic segment was calculated by dividing the total cluster number by the length, and then the average density across all dendritic segments was calculated for each neuron. A total of 6-10 neurons were analyzed per experiment, and 3-5 independent experiments were performed using different batches of neurons.

#### Statistical analysis

Data are shown as mean ± SEM, and n refers to the number of cells or independent experiments, as specified in each figure legend. Statistical analyses were performed in GraphPad Prism 8 software (GraphPad, La Jolla, CA, USA). For comparisons between two groups, Student’s t-test or one-sample t-test was used. For comparisons between three or more samples, one-way ANOVA followed by Tukey’s post hoc test was used. Comparisons between three or more groups with two independent variables were analyzed using two-way ANOVA followed by the post hoc tests recommended by Prism; all significant post hoc testing differences are defined as specific P values (correct to three decimal places) in the figures. All conditions statistically different from controls are indicated by p values labeled above columns in each figure. The complete output from Prism for each of the statistical analyses is provided in a supplementary Excel file (Supplementary Table 1).

## Results

### Shank3-Ca_V_1.3 interaction requires the Shank3 PDZ domain and Ca_V_1.3 PDZ-binding motif

Prior studies indicate that the Shank3 PDZ and SH3 domains interact directly with the C terminal-ITTL motif of Ca_V_1.3_L_ and an adjacent proline-rich region, respectively (Perfitt et al., 2020; Zhang et al., 2005). However, structural studies indicated that the Shank3 SH3 domain is atypical and has only weak (or no) interaction with multiple Ca_V_1.3-based proline-rich peptides (Ishida et al., 2018; Ponna et al., 2017). In addition, an N-terminal extension to the Shank3 PDZ domain is critical for high-affinity interactions with GKAP (Zhou et al., 2016). Therefore, we further investigated the roles of the Shank3 SH3 and PDZ domains in interactions with Ca_V_1.3_L_.

We generated five GST-Shank3 fusion proteins containing different segments of the amino acid sequence between residue 325 (N-terminal to the SH3 domain) to residue 664 (C-terminal to the PDZ domain) (Figure 1A). GST fusion proteins (or a GST negative control) were individually incubated with lysates of HEK293T cells expressing the entire C terminal domain of the Ca_V_1.3_L_ α1 subunit preceded by an HA epitope tag (HA-Ca_V_1.3-CTD), and protein complexes were isolated using magnetic glutathione beads (Figure 1B). We detected similar robust binding of HA-Ca_V_1.3-CTD to the three GST-Shank3 fusion proteins containing the PDZ domain; truncation of the SH3 domain or internal deletion of residues 543-564 (N-terminal PDZ extension) had no substantial impact on the interaction. Moreover, we did not detect any interaction of the HA-Ca_V_1.3-CTD with any fusion protein lacking the PDZ domain (containing only the SH3 domain). We then investigated interactions of a non-overlapping library of GST fusion proteins spanning the entire Shank3 protein with HA-tagged full-length Ca_V_1.3_L_ (Figure 1C). While full-length HA-Ca_V_1.3_L_ interacted with the GST-Shank3-PDZ domain, we did not detect interaction with any other GST-Shank3 fusion protein (Figure 1D). Taken together, these findings indicate that the Shank3 PDZ domain is primarily responsible for binding to Ca_V_1.3_L_, and that the Shank3 SH3 domain has a minimal role in the interaction.

### The presence of Ca_V_β subunits aids Shank3 assembly with Ca_V_1.3 LTCCs

Although Ca_V_1.3_L_ can directly bind to the Shank3 PDZ domain, it is possible that LTCC auxiliary subunits also play a role. Therefore, we investigated the impact of Ca_V_β subunits on the interaction by performing HA co-immunoprecipitation (co-IP) experiments from lysates of HEK293T cells expressing HA-Ca_V_1.3_L_, α2δ, with or without Flag-tagged β subunits (Flag-β3 or Flag-β2a), and either GFP or GFP-Shank3. Although GFP-Shank3 co-immunoprecipitated with HA-Ca_V_1.3_L_ in the absence of β subunits, co-expression of FLAG-β3 or -β2a enhanced the co-precipitation of GFP-Shank3 (Figure 2A, B). Interestingly, while FLAG-β3 significantly increased the co-precipitation of GFP-Shank3 by ∼2-fold, FLAG-β2a had a significantly greater ∼4-fold effect, even though FLAG-β3 and -β2a were expressed at similar levels. Moreover, co-expression of GFP-Shank3 increased by 2-3-fold the amounts of HA-Ca_V_1.3_L_ that were immunoprecipitated relative to the GFP control, independent of whether or which β subunit was co-expressed. These data indicate that Shank3 indeed associates with the full length Ca_V_1.3_L_ and that β subunits may stabilize the interaction.

To further explore the role of β subunits in Ca_V_1.3-Shank3 interaction, we incubated lysates of HEK293T cells expressing HA-Ca_V_1.3 and α2δ with or without β3 or β2a subunit with GST or GST-Shank3-PDZ. As seen in Figure 1A, HA-Ca_V_1.3 associated with GST-Shank3-PDZ on magnetic glutathione beads in the absence of β subunits. However, the co-expression of either FLAG-β2a or FLAG-β3 had no significant impact on the the amount of HA-Ca_V_1.3 that associated with GST-Shank3-PDZ (Figure 2D, E). These data indicate that β subunits do not affect the direct interaction of the Ca_V_1.3 α1 subunit with the Shank3 PDZ domain, suggesting that the ability of β subunits to enhance full-length Shank3 co-immunoprecipitation with full length Ca_V_1.3 (Figure 2A, B) requires other domains in Shank3.

To test the hypothesis that Shank3 may interact with LTCCs β subunits in the absence of Ca_V_1.3, we co-expressed FLAG-β3 or -β2a in HEK293T cells with either full-length GFP-Shank3, GFP-Shank3-ΔPDZ (with an internal deletion of the PDZ domain) or a GFP control. Immunoprecipitation using an anti-GFP antibody revealed that significantly more FLAG-β3 than FLAG-β2a associated with full length GFP-Shank3, but that the amounts of co-precipitated β subunit were unaffected by deletion of the PDZ domain (Figure 3A, B). However, reciprocal immunoprecipitations using a FLAG antibody indicated that similar amounts of full length GFP-Shank3 associated with FLAG-β3 or FLAG-β2a. The amount of GFP-Shank3 associated with both FLAG-β3 and FLAG-β2a appeared to be reduced by deletion of the PDZ domain (Figure 3C, D), but the reduction was not statistically significant in post hoc tests (Supplementary Table 1). In an effort to determine which domains in Shank3 are sufficient for β subunit binding, we investigated the interaction of full-length FLAG-β3 or FLAG-β2a with our family of GST-Shank3 fusion proteins (Figure 1C). However, we failed to detect interactions of either FLAG-β3 or FLAG-β2a with any of the GST-Shank3 fusion proteins (Supplemental Figure 1). Taken together, these data indicate that LTCC β subunits can associate with Shank3 independent of the Ca_V_1.3 α1 subunit, and that this interaction does not strictly require the Shank3 PDZ domain, although there may be some modest quantitative effects. The interaction of Shank3 with β subunits may contribute to β subunit-dependent enhancement of Shank3 association with full-length Ca_V_1.3 observed in Figures 2A and 2B.

### The Shank3 PDZ domain mediates assembly of complexes containing multiple Ca_V_1.3_L_ LTCCs

The amount of HA-Ca_V_1.3_L_ immunoprecipitated using an HA antibody was consistently increased by GFP-Shank3 co-expression, independent of the β subunit (Figure 2C). Since the HA antibody immunoprecipitated only a fraction of the total HA-Ca_V_1.3_L_ from these lysates, we hypothesized that this might be due to the clustering of multiple HA-Ca_V_1.3_L_ subunits by Shank3 multimers (Naisbitt et al., 1999). To directly test this hypothesis (Figure 4A), we co-expressed mCherry-tagged Ca_V_1.3 (mCherry-Ca_V_1.3_L_) and HA-Ca_V_1.3_L_, along with α2δ and FLAG-β2a subunits and either GFP, GFP-Shank3, or GFP-Shank3-ΔPDZ. GFP-Shank3 specifically and efficiently co-precipitated with HA-Ca_V_1.3_L_ relative to the GFP control, and deletion of the PDZ domain significantly reduced the co-immunoprecipitation by ∼80% (Figure 4B, C). Presumably, the residual co-immunoprecipitation of GFP-Shank3-ΔPDZ with HA-Ca_V_1.3_L_ is mediated by the β2a subunit. Notably, mCherry-Ca_V_1.3 was readily detected in HA-immune complexes isolated from cells co-expressing GFP-Shank3, but only low levels of mCherry-Ca_V_1.3 were detected in complexes isolated from cells co-expressing GFP or GFP-Shank3-ΔPDZ (Figure 4B, D). These data provide direct biochemical support for the hypothesis that Shank3 can cluster multiple Ca_V_1.3_L_ in a complex and that the PDZ domain is crucial for this clustering.

### Shank3-dependent in vitro clustering of Ca_V_1.3_L_ in disrupted by Ca^2+^/calmodulin

Next, we tested whether the assembly of mCherry-Ca_V_1.3_L_ with HA-Ca_V_1.3_L_ was affected by Ca^2+^/CaM. Lysates of cells co-expressing mCherry-Ca_V_1.3_L_, HA-Ca_V_1.3_L_, α2δ and FLAG-β2a subunits with either GFP or GFP-Shank3 were HA-immunoprecipitated under basal conditions (with EDTA) or following the addition of Ca^2+^/CaM (Figure 5A). In GFP control cell lysates, the addition of Ca^2+^/CaM slightly increased the amount of mCherry-Ca_V_1.3 detected in the HA-immune complexes in 5 out of 6 experiments, but the average ∼1.5-fold increase was not statistically significant. As seen in Figure 4, the co-expression of GFP-Shank3 significantly increased the levels of mCherry-Ca_V_1.3_L_ detected in HA-immune complexes under basal (EDTA) conditions, but the addition of Ca^2+^/CaM significantly reduced the levels of co-precipitated mCherry-Ca_V_1.3_L_ (Figure 5B). Moreover, Ca^2+^/CaM addition also significantly reduced the levels of GFP-Shank3 that co-precipitated with HA-Ca_V_1.3_L_ (Figure 5C). These data demonstrate that the Shank3-dependent assembly of complexes containing multiple Ca_V_1.3 α1 subunits can be disrupted by the addition of Ca^2+^/CaM to cell lysates, potentially simulating conditions in close proximity to the activated LTCCs in the plasma membrane.

### Shank3 stabilizes Ca_V_1.3 LTCCs in the plasma membrane under basal conditions *in situ*

As an initial test of the hypothesis that Shank3 clusters Ca_V_1.3_L_ in the plasma membrane, we used TIRF microscopy to detect fluorescent proteins residing within ∼100 nm of the cover slip in live HEK293 cells co-expressing mCherry-Ca_V_1.3_L_, α2δ and β2a/β3 subunits with either GFP or GFP-Shank3. We detected mCherry puncta in cells co-expressing GFP-Shank3, or the GFP control (Figure 6A), presumably predominantly reflecting LTCCs that had been trafficked to the plasma membrane. Moreover, GFP-Shank3 strongly colocalized with many of the mCherry-Ca_V_1.3_L_ puncta (Figure 6A). Notably, mCherry puncta were significantly more intense in cells expressing GFP-Shank3 than in GFP control cells (Figure 6B), consistent with the hypothesis that GFP-Shank3 increases the number of mCherry-tagged α1 subunits within each puncta. Repeated imaging of these cells over 3-5 minutes indicated that mCherrry-Ca_V_1.3_L_ clusters generally appeared transiently in the TIRF images of GFP control cells (Figure 6Ai), whereas in the presence of GFP-Shank3 most mCherry-Ca_V_1.3_L_ puncta in the TIRF images remained for the duration of the imaging session (Figure 6Aii). Moreover, mCherry-Ca_V_1.3_L_ puncta were quite motile within the plane of the plasma membrane in GFP control cells, moving at average speeds of ∼0.25 µm/s, whereas in cells expressing GFP-Shank3 they moved significantly slower (∼0.1 µm/s) (Figure 6D). Figures 6C and 6D summarize data from multiple experiments co-expressing either FLAG-β3 (solid symbols) or FLAG-β2a (open symbols), indicating that the impact of GFP-Shank3 on mCherry-Ca_V_1.3_L_ is essentially independent of the identity of the β subunit. Taken together, these data are consistent with a model in which Shank3 stabilizes Ca_V_1.3_L_ α1 subunit clusters in HEK293 cell plasma membranes.

### The Shank3 PDZ domain mediates basal Ca_V_1.3 clustering in intact cells

We next tested for an effect of LTCC-mediated Ca^2+^ influx on mCherry-Ca_V_1.3 clustering in transfected HEK293 cells co-expressing GFP-Shank3. LTCCs were activated pharmacologically using Bay K8644 (BayK) (10 µM) in the absence or presence of added extracellular Ca^2+^ while monitoring surface localized mCherry-Ca_V_1.3_L_ and GFP-Shank3 in single HEK293 cells by live-cell TIRF imaging. After collecting baseline data in a 0 mM Ca^2+^ buffer, cells were switched to 0 mM Ca^2+^ buffer with BayK for several minutes, and then to 2.5 mM Ca^2+^ buffer with BayK. Figure 7A shows a single representative cell at t=0, and marks a region of interest containing co-localized mCherry-Ca_V_1.3_L_/GFP-Shank3 clusters. The ratio of mCherry/GFP fluorescence in this region of interest was measured at 5 s intervals and plotted in Figure 8B under each of the buffer conditions, with about a one-minute gap as the buffer solutions were switched; the insets show images of the region of interest at selected time points. The mCherry/GFP ratio was relatively stable throughout the incubation with 0 Ca^2+^, in the absence or presence of BayK. However, addition of the Ca^2+^/BayK buffer decreased the mCherry/GFP ratio within one minute of buffer changing, mainly due to a substantial reduction in the mCherry-Ca_V_1.3_L_ intensity (Figure 7C). Figure 7D shows the mCherry/GFP ratio from 12 cells in each of the three buffer conditions, normalized to the 0 Ca^2+^ buffer, indicating that Ca^2+^ influx significantly reduces the intensity of mCherry-Ca_V_1.3_L_ clusters colocalized with GFP-Shank3.

In order to provide further insight into the role of Shank3 in Ca_V_1.3 LTCC clustering *in situ*, HEK293 cells transfected with mCherry-Ca_V_1.3 and GFP, GFP-Shank3, or GFP-Shank3-ΔPDZ were incubated for 10-15 min in a HEPES buffer containing 0 or 2.5 mM Ca^2+^, in the absence or presence of BayK (Figure 8A), and then fixed for imaging using a TIRF microscope. As seen in live HEK293 cells (Figures 6 and 7), mCherry-Ca_V_1.3_L_ puncta were readily detected near the cell surface under all conditions. We first quantified the puncta intensity (Figure 8B) and density (Figure 8C) under each incubation condition. In cells co-expressing GFP (gray circles/bars), both parameters were unaffected by incubation of the cells with or without extracellular Ca^2+^ and/or BayK. The intensity of mCherry-Ca_V_1.3_L_ puncta was significantly increased ∼2-fold by the co-expression of GFP-Shank3 (blue squares/bars) when cells were incubated in the absence of extracellular Ca^2+^ (+/- BayK) or with Ca^2+^ in the absence of BayK. However, incubation of cells expressing GFP-Shank3 with both Ca^2+^ and BayK significantly reduced the mCherry-Ca_V_1.3_L_ puncta intensity to levels observed in GFP control cells. Notably, GFP-Shank3 co-expression had no effect on the mCherry-Ca_V_1.3_L_ puncta density, and the puncta density in cells expressing GFP-Shank3 was unaffected by the Ca^2+^/BayK incubations. Importantly, the co-expression of GFP-Shank3-ΔPDZ (red triangles/bars) had no significant effect on the intensity or density of mCherry-Ca_V_1.3_L_ puncta compared to the GFP control under any condition tested.

In parallel, we quantified the co-localization of GFP signals with mCherry-Ca_V_1.3_L_ using the ICQ method (Figure 8D). The ICQ score in cells expressing soluble GFP and mCherry-Ca_V_1.3_L_ was very low (∼0.05) under all conditions, as expected for mostly random overlap. In contrast, GFP-Shank3 significantly colocalized with mCherry-Ca_V_1.3_L_ puncta (ICQ ∼0.25) when cells were pre-incubated in the absence of extracellular Ca^2+^ (+/- BayK) or with Ca^2+^ in the absence of BayK. However, the simultaneous addition of Ca^2+^ and BayK significantly decreased the ICQ to ∼0.15. Moreover, GFP-Shank3-ΔPDZ was only weakly co-localized with mCherry-Ca_V_1.3_L_ (ICQ ∼ 0.15), independent of the specific cell incubation condition. Taken together, these data indicate that the Shank3 PDZ domain is essential for efficient colocalization with Ca_V_1.3, and also for efficient Ca_V_1.3 clustering under basal conditions, and that LTCC-mediated Ca^2+^ influx disrupts the effect of Shank3.

### Endogenous Shank3 clusters Ca_V_1.3_L_ in cultured hippocampal neurons

Previous over-expression studies in cultured neurons indicate that Shank3 interaction with the Ca_V_1.3 C-terminal domain facilitates Ca_V_1.3 LTCC surface expression in dendrites (Stanika et al., 2016; Zhang et al., 2006). However, the role of endogenous Shank3 has not been investigated. Therefore, we expressed Ca_V_1.3_L_ with an extra-cellular HA tag (sHA-Ca_V_1.3), α2δ and either FLAG-β3 or -β2a, with or without a well-characterized highly effective and specific shRNA to knock down endogenous Shank3 expression (Perfitt et al., 2020; Verpelli et al., 2011). First, we confirmed the efficacy of Shank3 knockdown in DIV21 neurons that were co-transfected to express the LTCC subunits at DIV14. In non-transfected neurons (NT) or neurons expressing control nonsense shRNA (nssh), punctate staining for endogenous Shank3 was readily detected in the soma and dendrites (Supplemental Figure 2A), consistent with previous studies (Perfitt et al., 2020; Zhang et al., 2005). Moreover, the intensity of somatic Shank3 puncta was essentially identical in non-transfected neurons and neurons expressing the control RNA (nssh/NT ratios: 1.19±0.14 and 1.14±0.12 in neurons co-expressing β3 and β2a subunits, respectively) (Supplemental Figure 2B). However, expression of the Shank3-shRNA (SK3-sh) significantly reduced the intensity of endogenous Shank3 fluorescence by ∼80% (SK3-sh/NT ratios: 0.28±0.04 and 0.17±0.03 in neurons co-expressing β3 and β2a subunits, respectively) (Supplemental Figure 2B). The high density of non-transfected dendrites in these cultures/images precluded quantification of dendritic Shank3 levels in transfected neurons. These data confirm reliable knock down of Shank3 protein expression by the shRNA under the current experimental conditions, albeit with somewhat reduced efficacy than we observed previously in younger neurons (Perfitt et al., 2020).

We then examined the impact of Shank3 knockdown on sHA-Ca_V_1.3 cell surface expression. Consistent with previous studies (Moreno et al., 2016; Stanika et al., 2016; Zhang et al., 2016), we detected dense cell surface clusters of sHA-Ca_V_1.3_L_ in neurons expressing control shRNA, that were partially colocalized with endogenous Shank3 on the soma and dendrites (Figure 9A). The Shank3 shRNA clearly suppressed endogenous Shank3 staining in the soma and dendrites, but some residual Shank3 still colocalized with sHA-Ca_V_1.3_L_ clusters. Notably, Shank3 knockdown significantly decreased the average intensity of sHA-Ca_V_1.3_L_ clusters in neuronal dendrites, when expressed with either FLAG-β3 (Figure 9C and Supplemental Figure 3B) or FLAG-β2a (Supplemental Figure 4C). However, there was only a trend for a decrease of somatic sHA-Ca_V_1.3_L_ cluster intensity (Figure 9B, Supplemental Figure 3A, and Supplemental Figure 4A). In contrast, Shank3 knockdown significantly reduced the density (number) of both somatic and dendritic sHA-Ca_V_1.3_L_ clusters when expressed with either FLAG-β3 (Figure 9B, C) or FLAG-β2a (Supplemental Figure 4). To explore if Shank3 specifically affects Ca_V_1.3_L_ LTCC clustering, we examined Ca_V_1.2 LTCC cell surface expression with or without Shank3 knockdown (Figure 10). Surface sHA-Ca_V_1.2 clusters were not strongly colocalized with endogenous Shank3 in control cells (Figure 10A), as expected because the Ca_V_1.2 α1 subunit does not directly interact with Shank3 (Zhang et al., 2005). Neither the intensity nor the density of dendritic sHA-Ca_V_1.2 clusters were affected by Shank3 knockdown and the intensity of somatic Ca_V_1.2 clusters also remained unchanged, although the density of somatic Ca_V_1.2 clusters was modestly, but significantly, reduced by Shank3 knockdown (Figure 10B, C and Supplemental Figure 5A, B). In combination, these data indicate that endogenous Shank3 plays an important role in the dendritic clustering and overall surface expression of Ca_V_1.3_L_ LTCCs under basal conditions.

**Figure 9.**
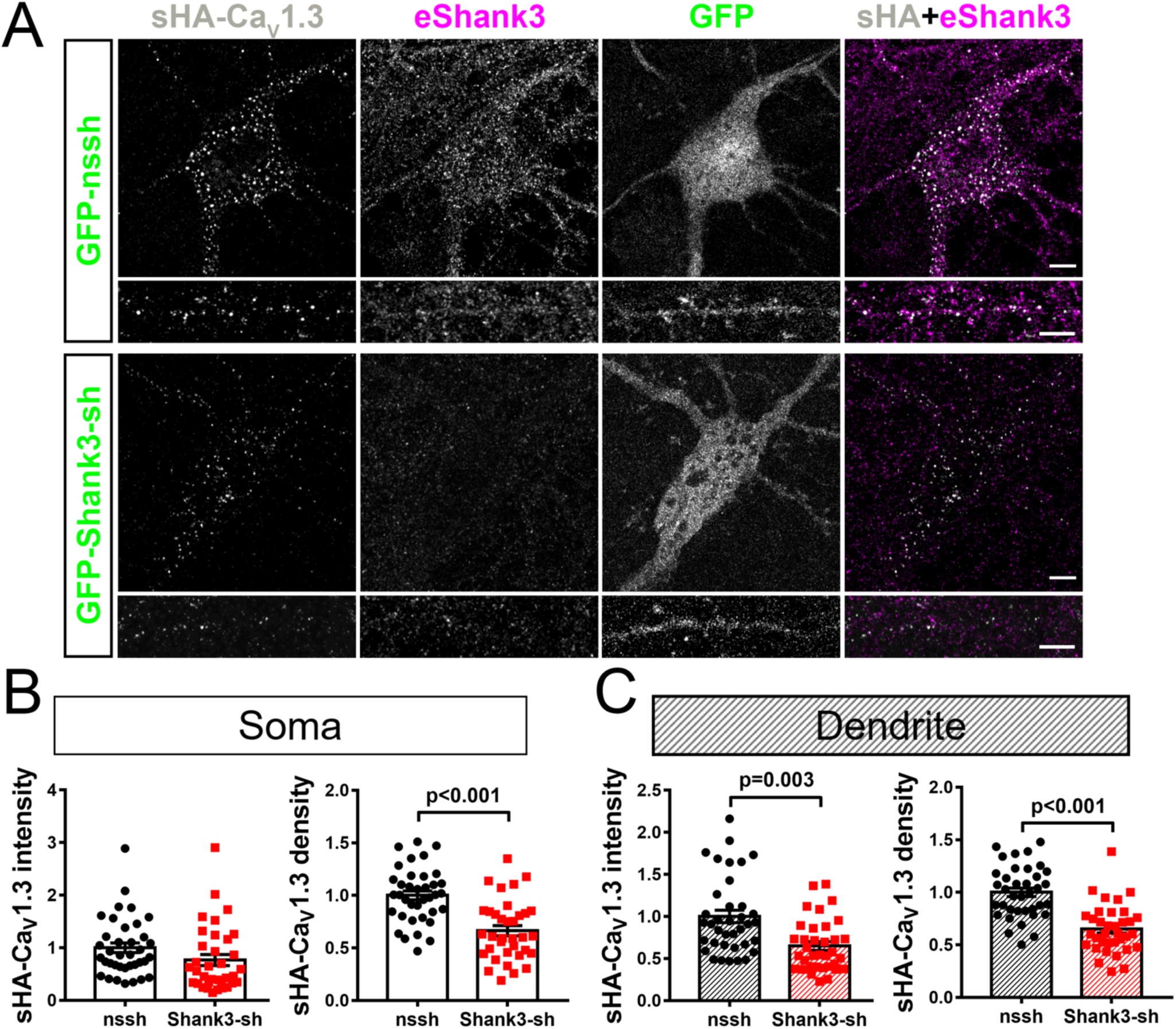
Effects of Shank3 knock-down on surface-expressed Ca_V_1.3 puncta in neurons. Primary rat hippocampal neurons (14 DIV) expressing sHA-Ca_V_1.3 and FLAG-β3 with either GFP-nonsense shRNA (GFP-nssh) or GFP-Shank3 shRNA (GFP-Shank3-sh) were live-immunostained for the HA tag at DIV21, fixed, permeabilized and then immunostained for endogenous Shank3 and DAPI. Neurons were imaged using Airyscan super-resolution confocal microscopy. A) Representative images of soma and dendrites. Scale bar, 5 μm. B) and C) Quantification of sHA-Ca_V_1.3 cluster intensity and cluster density, respectively, of n = 37 (GFP-nssh) or 35 (GFP-Shank3-sh) neurons from three independent cultures/transfections; comparisons made using an unpaired t-test.

**Figure 10.**
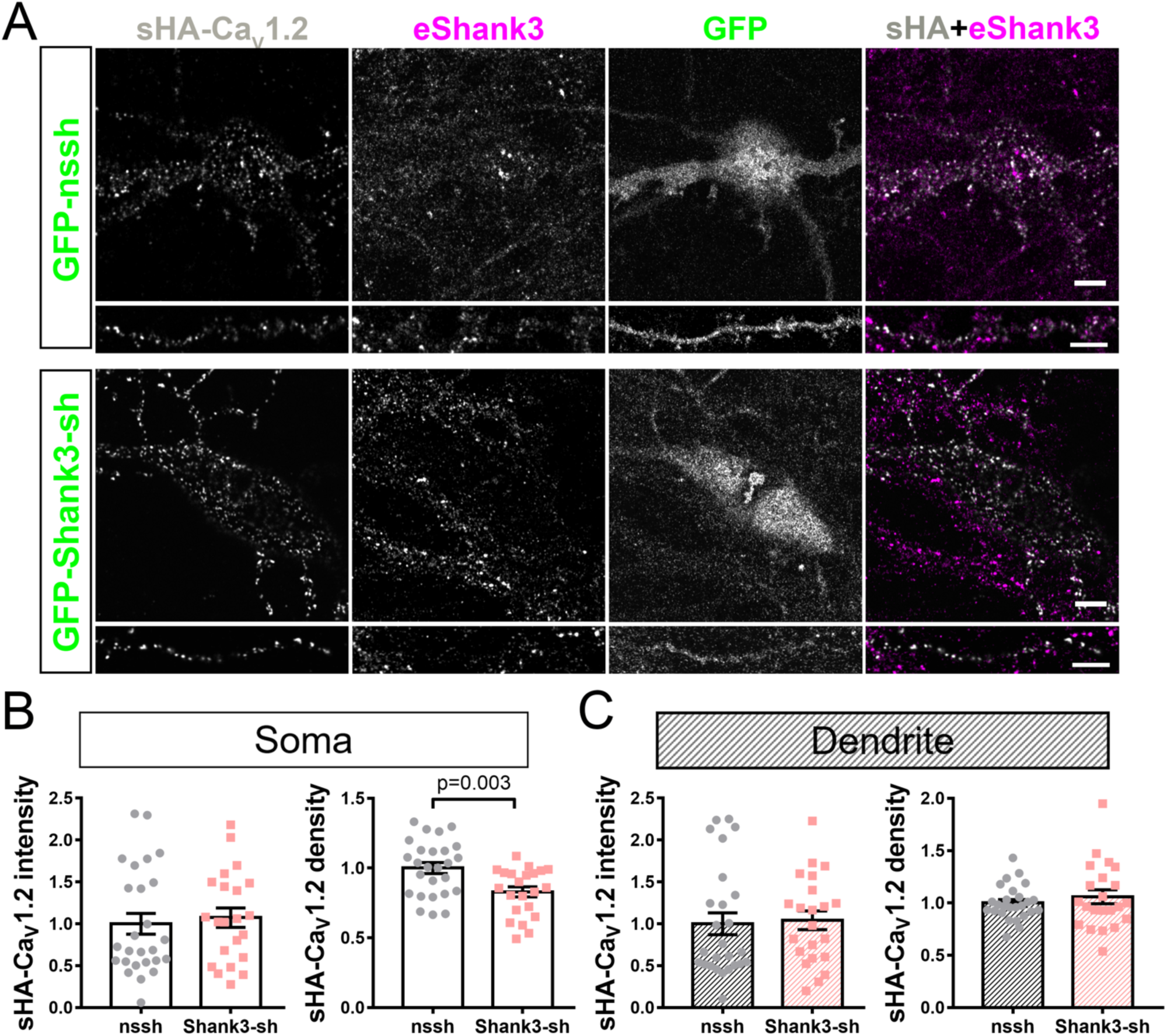
Shank3 knock-down has no effect on Ca_V_1.2 surface puncta intensity in neurons. Primary rat hippocampal neuron (14 DIV) expressing sHA-Ca_V_1.2 and FLAG-β3 with either GFP-nonsense shRNA (GFP-nssh) or GFP-Shank3 shRNA (GFP-Shank3-sh) were live-immunostained for the HA tag at DIV21, fixed, permeabilized and then immunostained for endogenous Shank3 and DAPI. Neurons were imaged using Airyscan super-resolution confocal microscopy. A) Representative images of soma and dendrites. Scale bar, 5 μm. B) and C) Quantification of sHA-Ca_V_1.2 cluster intensity and cluster density from n = 26 (GFP-nssh) or 22 (GFP-Shank3-sh) neurons from three independent cultures/transfections; comparisons made using an unpaired t-test.

## Discussion

Here we provide new insights into the role of Shank3 in controlling Ca_V_1.3 LTCC clustering. Complementary co-immunoprecipitation and fluorescence microscopy studies using heterologous cells demonstrate that a direct interaction between the C-terminal domain of the Ca_V_1.3 α1 subunit and the PDZ domain of Shank3 can mediate the clustering of multiple Ca_V_1.3 LTCCs. Our data also indicate that LTCC β3 or β2a auxiliary subunits facilitate Shank3 clustering of Ca_V_1.3 LTCCs, perhaps by directly (or indirectly) interacting with Shank3 independent of the Ca_V_1.3 α1 subunit. Significantly our data indicate Shank3-Ca_V_1.3 association and Ca_V_1.3 clustering can be disrupted by increasing Ca^2+^ and CaM in cell lysates, indicating that Ca_V_1.3 clustering can be dynamically modulated by LTCC activation. Finally, we confirmed prior studies showing that endogenous Shank3 partially colocalizes with plasma membrane Ca_V_1.3 clusters in cultured hippocampal neurons, and we showed that Shank3 knockdown disrupted dendritic Ca_V_1.3 clustering. Taken together, our data substantially advance our understanding of the role of Shank3 in Ca_V_1.3 LTCC clustering.

### The Shank3 PDZ domain mediates Ca_V_1.3_L_ LTCC clustering under basal conditions

Shank proteins are multi-domain scaffolding proteins localized to excitatory synapses, where they coordinate the assembly of several multiprotein complexes (Naisbitt et al., 1999; Sheng & Hoogenraad, 2007; Sheng & Kim, 2000). It is well established that Shank PDZ domains can interact with multiple synaptic proteins, and deletion of the Shank3 PDZ domain in mice results in synaptic dysfunction and autism-related behavioral phenotypes (Peça et al., 2011), demonstrating the importance of the Shank3 PDZ domain interactions.

Here, our in vitro studies using GST fusion proteins showed that the Shank3 PDZ domain is necessary and sufficient for binding to the C-terminal domain of Ca_V_1.3_L_ or to the full length Ca_V_1.3_L_ α1 subunit. In contrast to some prior reports, our data provided no indication that the Shank3 SH3 domain plays a significant role in this interaction (Zhang et al., 2005). The reasons for this discrepancy are unclear, but it is possible that a low affinity interaction of Ca_V_1.3_L_ with the SH3 domain (e.g., Ishida et al., 2018) could not be detected under our experimental conditions. These in vitro studies also indicated that β auxiliary subunits had a hitherto unappreciated role in facilitating Shank3 interactions with Ca_V_1.3_L_ LTCCs, apparently by also interacting with Shank3. However, preliminary in vitro studies (Supplementary Figure 1) failed to detect a direct interaction of either β2a or β3 with the PDZ domain or any other tested fragment of Shank3. Further studies are required to define the domains in Shank3 and the β subunit that mediate this interaction.

We then adapted our co-immunoprecipitation assay to detect interactions between co-expressed Ca_V_1.3_L_ LTCCs with different epitope tags, demonstrating that Shank3 can mediate the assembly of complexes containing multiple Ca_V_1.3_L_ LTCCs, and that the Shank3 PDZ domain is essential for assembly of these complexes. We extended these studies to investigate the impact of Shank3 on Ca_V_1.3_L_ clustering in the plasma membrane of heterologous (HEK293) cells. TIRF microscopy provided no evidence that co-expression of Shank3 modulated cell surface expression levels (Ca_V_1.3_L_ puncta density) in HEK293 cells under basal cell incubation conditions. Rather, we found that Shank3 increased the average intensity (or brightness) of cell surface Ca_V_1.3_L_ puncta in both live-cell imaging studies (Figure 6) and in fixed cells (Figure 7). We interpret the increased signal intensity/brightness as an increase of the average number of mCherry-Ca_V_1.3_L_ α1 subunits within each puncta, or Ca_V_1.3_L_ LTCC clustering, and this increase was also dependent on the Shank3 PDZ domain. Prior cell biology studies have indicated that Ca_V_1.3_S_ and Ca_V_1.3_L_ variants can “self-cluster” in the plasma membrane, with each cluster containing an average of ∼8 α1 subunits (Moreno et al., 2016). Thus, the Shank3-dependent clustering observed in our co-immunoprecipitation and cell imaging studies may representing higher-order assembly of intrinsic Ca_V_1.3_L_ clusters.

Consistent with these biochemical and heterologous cell studies, we found that knocking down Shank3 expression in cultured neurons had a significant impact on expression of Ca_V_1.3_L_ (Figure 9) but not Ca_V_1.2 (Figure 10) in the plasma membrane. Shank3 knockdown significantly decreased the overall density of cell surface Ca_V_1.3_L_ puncta in both the soma and dendrites, indicating that Shank3 enhances cell surface expression of Ca_V_1.3_L_ LTCCs in neurons. Similar decreases in density were observed in neurons that co-expressed Ca_V_1.3_L_ with either the β2a or β3 subunits. These findings are consistent with prior reports indicating that Shank3 enhances Ca_V_1.3_L_ trafficking to the neuronal plasma membrane (Zhang et al., 2006). However, Shank3 knockdown also significantly decreased the intensity of surface Ca_V_1.3_L_ puncta in neuronal dendrites, but not in the soma, once again irrespective of the identity of the co-expressed β subunit. This observation indicates an additional dendritic role for Shank3 in increasing the number of Ca_V_1.3_L_ α1 subunits within each puncta. We interpret this observation as being consistent with the hypothesis that endogenous Shank3 promotes the clustering of Ca_V_1.3_L_ LTCCs in neuronal dendrites under basal culture conditions.

### Shank3 binding and Ca_V_1.3_L_ clustering is disrupted in the presence of Ca^2+^

We also hypothesized that Ca_V_1.3_L_ clustering might be sensitive to increased Ca^2+^. Neuronal depolarization has been shown to enhance the physical and/or functional coupling of Ca_V_1.2 and Ca_V_1.3_S_ LTCCs (Dixon et al., 2015; Moreno et al., 2016), and some data indicate that Ca^2+^/CaM can induce homodimerization of Ca_V_1.2 LTCCs (Fallon et al., 2009). However, we found that Ca^2+^/CaM addition to transfected HEK293 cell lysates dissociated Shank3 from Ca_V_1.3_L_ complexes and disrupted Shank-3 dependent co-immunoprecipitation of HA- and mCherry-tagged Ca_V_1.3_L_ (Figure 5). Moreover, we found that Shank3-colocalizes with Ca_V_1.3_L_ in HEK293 cell plasma membranes and Ca_V_1.3_L_ LTCCs clustering could be disrupted by the addition of both extracellular Ca^2+^ and the LTCC agonist, BayK8644, but not by either Ca^2+^ or BayK8644 alone (Figures 7 and 8). These data suggest Ca^2+^ influx via the LTCC itself causes these effects and that neither BayK8644-induced conformational changes (Marom et al., 2010) nor Ca^2+^ influx via endogenous HEK293 cell channels is sufficient to disrupt Shank3-binding and LTCC clustering. Interestingly, the N- and C-terminal lobes of CaM interact with motifs in the N- and C-terminal tails of Ca_V_1.3 LTCCs, respectively, in the presence of Ca^2+^ (Banerjee et al., 2018; Ben Johny et al., 2013). We speculate that these interactions result in conformational changes in the LTCC N- and C-terminal domains that interfere with binding of the Shank3 PDZ domain to the C-terminal ITTL motif of Ca_V_1.3_L_.

### The potential roles of Ca_V_1.3 channel clustering

Activation of neuronal LTCCs has been suggested to create local Ca^2+^ nanodomains near the plasma membrane that have privileged roles in initiating downstream signaling cascades, such as excitation-transcription coupling. It seems likely that the clustering of multiple LTCCs within a single complex facilitates the formation of Ca^2+^ nanodomains that are larger or attain higher Ca^2+^ concentrations, enhancing downstream signaling. In support of this notion, several different experimental approaches have indicated that Shank3 has a key role in facilitating Ca_V_1.3 LTCC-induced excitation-transcription coupling. We suggest that this facilitation of E-T coupling is due to the Shank3-dependent Ca_V_1.3_L_ clustering reported here. Although it may seem somewhat paradoxical that Shank3-dependent Ca_V_1.3_L_ clustering is disrupted by Ca^2+^influx, several other mechanisms undoubtedly contribute to the control of LTCC clustering and the dynamics of Ca^2+^ nanodomains. For example, clustering of Ca_V_1.3_S_ channels (which cannot bind Shank3) enhances Ca^2+^ influx by allowing for Ca^2+^/CaM-dependent functional coupling within the cluster (Moreno et al., 2016). However, even though Ca_V_1.3_L_ LTCCs form clusters with similar physical dimensions, they do not seem to be regulated by this Ca^2+^/CaM-dependent functional coupling mechanism. Further studies will be required to develop a deeper understanding of the molecular mechanisms controlling the regulation of Ca_V_1.3 splice variant clustering and the physiological significance of clustering. Since genetic variants of Shank3 and LTCCs in humans are being increasingly linked to autism spectrum disorders, schizophrenia and other neuropsychiatric disorders (Gauthier et al., 2010; Guilmatre et al., 2014; Martínez-Rivera et al., 2017; Monteiro & Feng, 2017; Pinggera et al., 2015), such studies also may provide insight into the pathophysiology of these disorders.

## Acknowledgements

This work was supported by Vanderbilt University and an endowed Louise B. McGavock Chair to R.J.C., and by an AHA fellowship to TJP. The content is solely the responsibility of the authors and does not necessarily represent the official views of the National Institutes of Health. Confocal imaging and analysis were performed in part through the use of the Vanderbilt Cell Imaging Shared Resource (supported by National Institutes of Health Grants CA68485, DK20593, DK58404, DK59637, and EY08126). We thank Drs. Craig Garner, Gerald Zamponi, Luk Van Parijs, and Diane Lipscombe for generously providing various original plasmids, as detailed in Key Resources Table.

## Conflict of interest disclosure

The authors declare that they have no competing interests.

## Author contributions

Q.Y. and R.J.C designed research; Q.Y. and T.L.P. performed biochemistry experiments; Q.Y. performed imaging experiments; L.H. prepared rat hippocampal neuronal cultures; Q.Y. and J.Q. analyzed data; Q.Y. and R.J.C. wrote the manuscript; J.Q. helped to modify the manuscript. All authors participated in the discussion, revision, and approval of the final manuscript.

## Key Resources Table

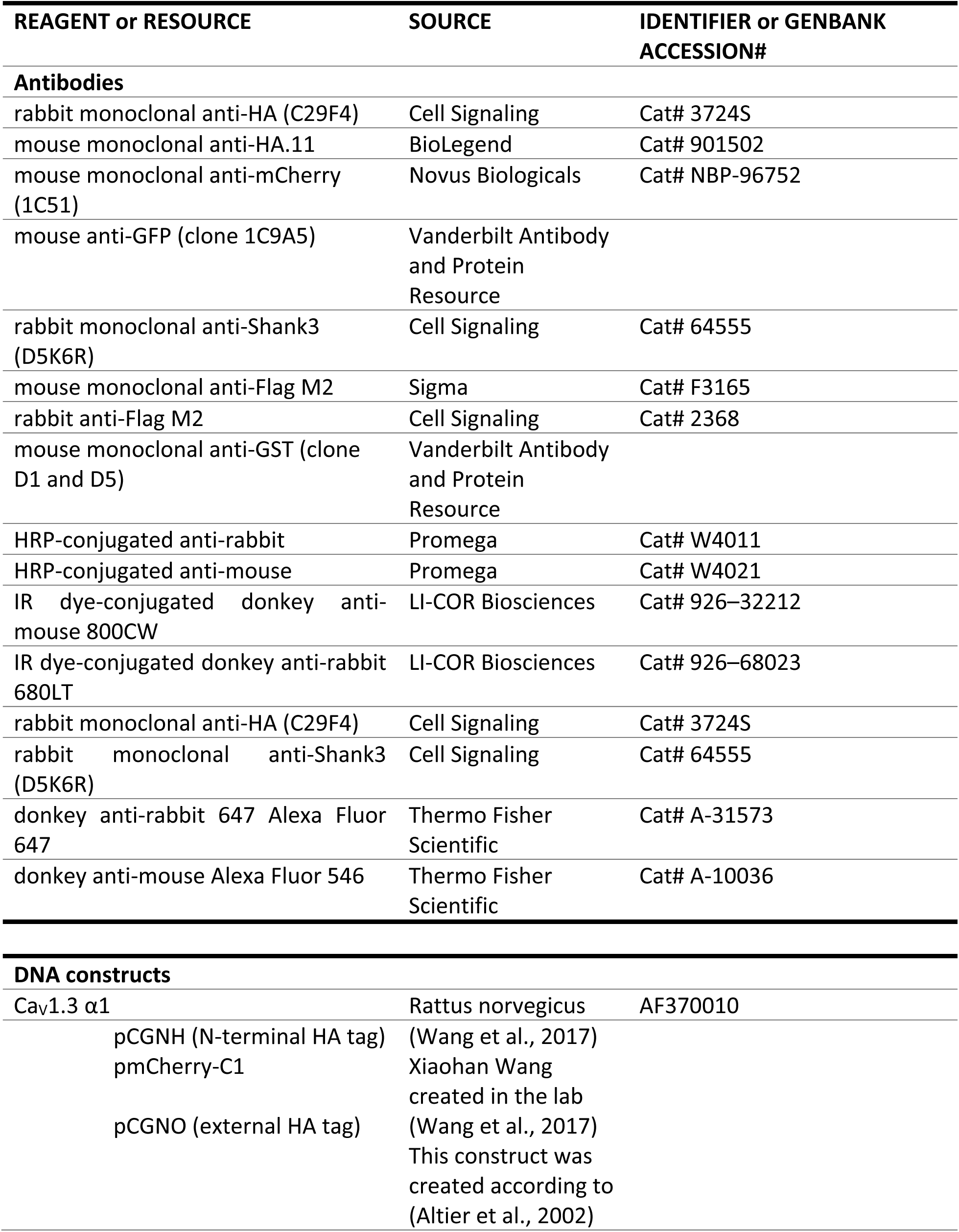
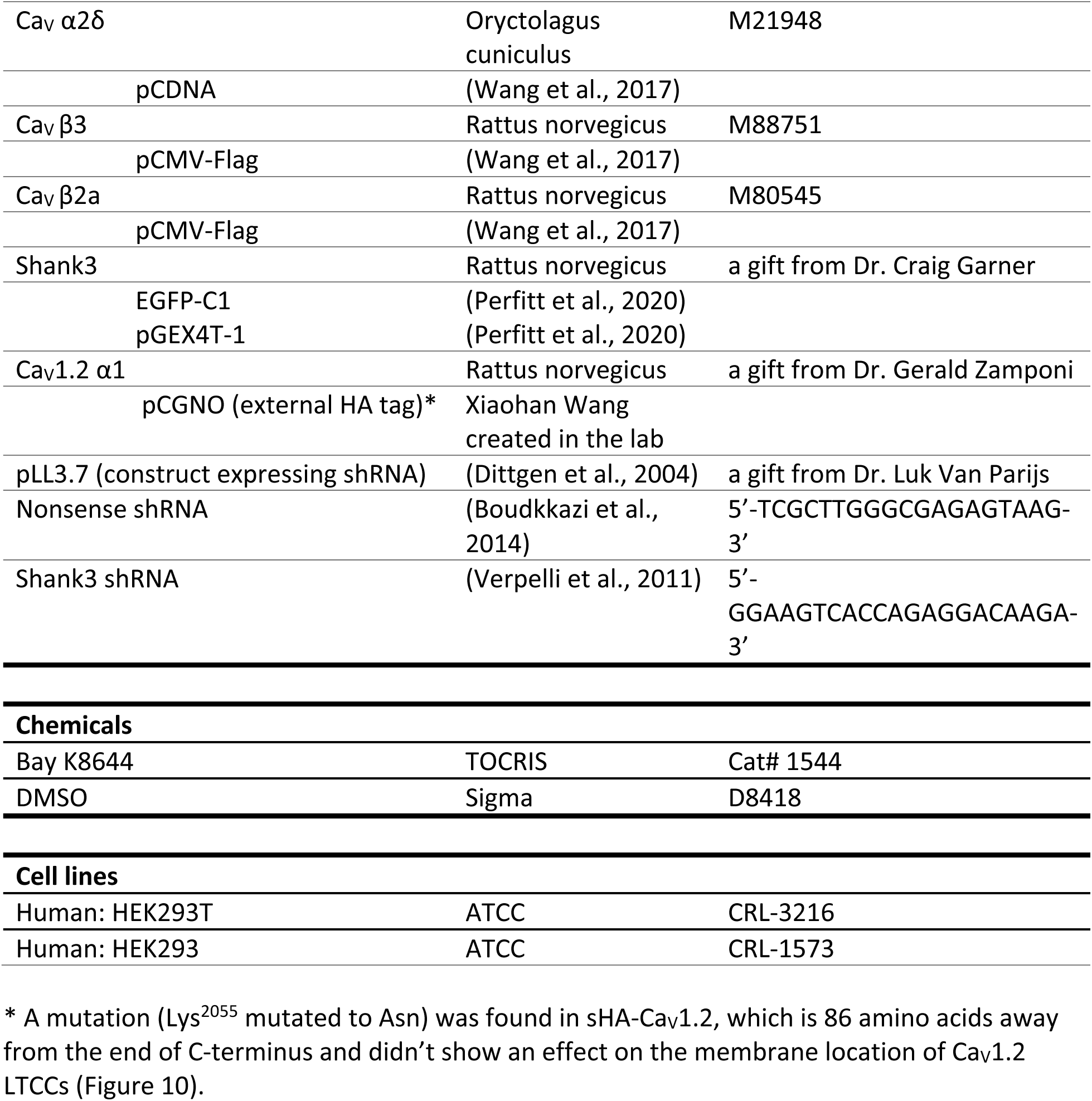

## Supplemental Figures and Legends

**Supplemental Figure 1 (related to Figure 3).**
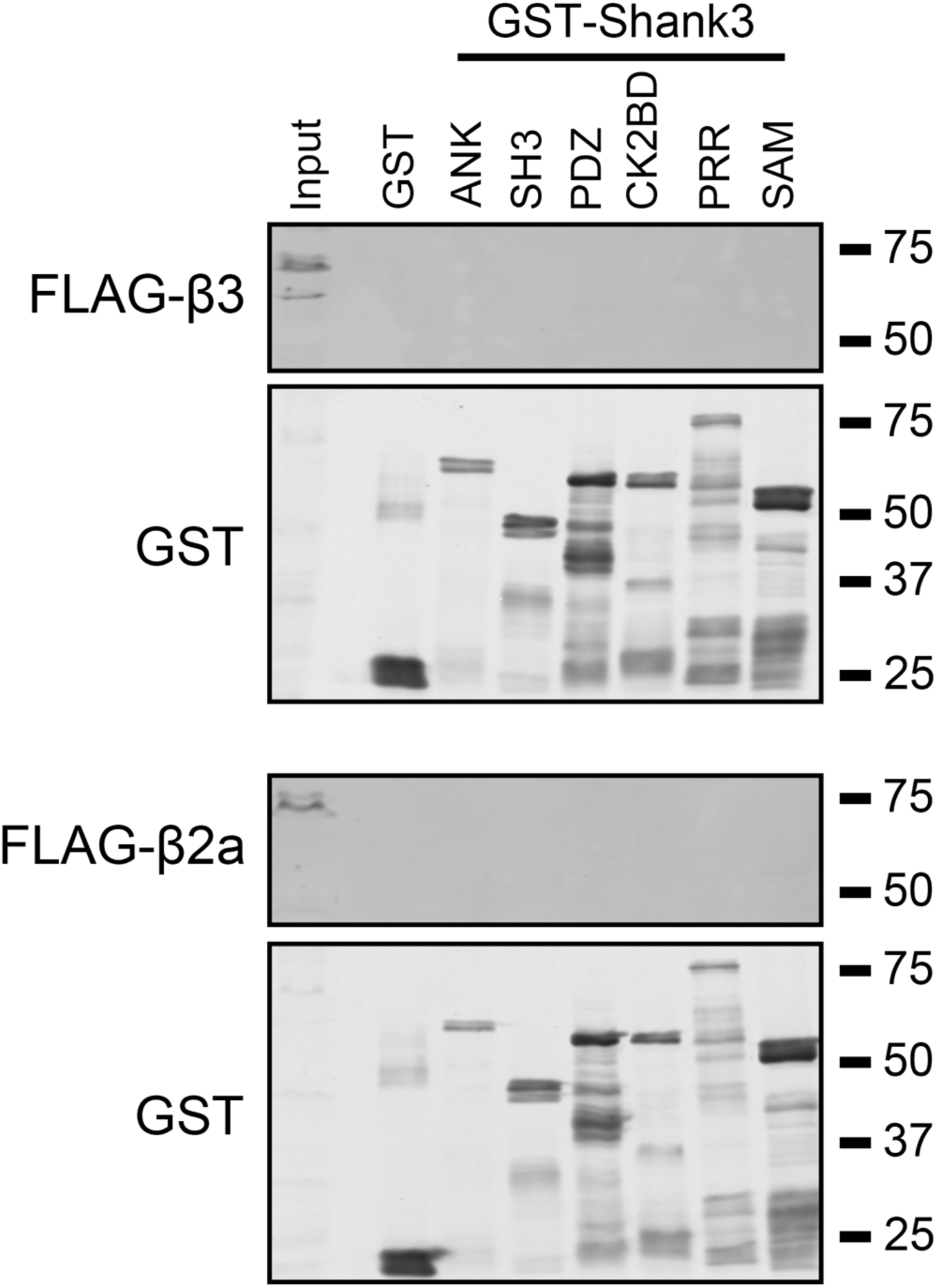
No detectable interaction of FLAG-β3 or -β2a with any GST-Shank3 fusion proteins. Soluble fractions of HEK293T cells expressing either FLAG-β3 (top) or -β2a (bottom) (Input) were incubated with GST or the indicated GST-Shank3 domain constructs (see Fig. 1C). Complexes were isolated using glutathione magnetic beads and then aliquots of the input and complexes were immunoblotted for the FLAG epitope or GST. Representative of three replicates with different transfected cell lysates.

**Supplemental Figure 2 (related to Figure 7).**
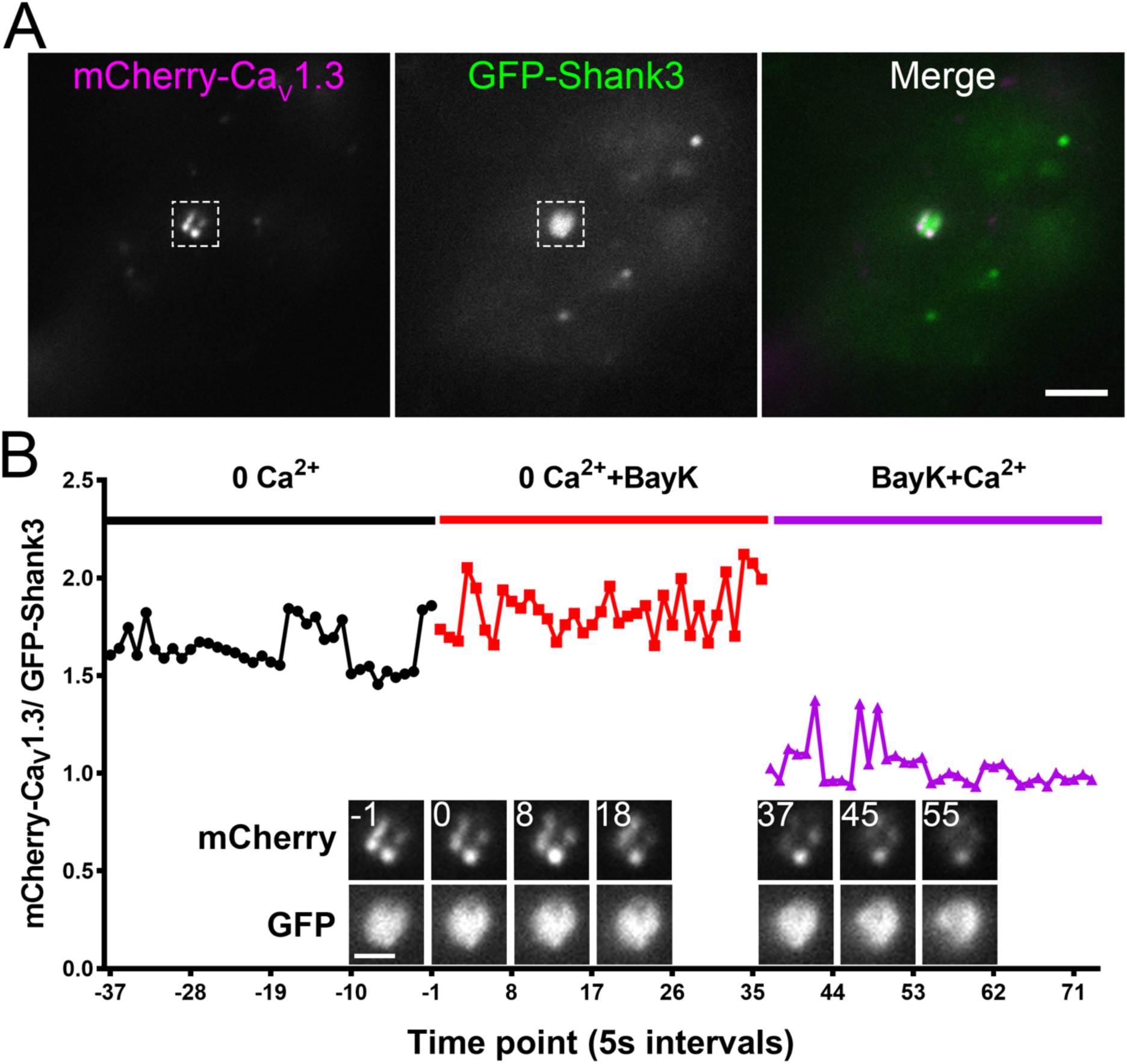
Ca^2+^ influx dissociates GFP-Shank3 from mCherry-Ca_V_1.3 in live HEK293 cells expressed FLAG-β2a. Data were collected in parallel with those shown in Figure 7, except that FLAG-β2a was co-expressed instead of FLAG-β3. The same cell incubation and imaging conditions were used. A) Representative mCherry, GFP and merged image of a HEK293 cell at the start of the experiment (scale bar, 5 µm). B) Quantification of the ratio of mCherry-Ca_V_1.3 to GFP-Shank3 signal intensity by time in the region of interest highlighted in panel A, with insets showing enlarged mCherry-Ca_V_1.3 (top row) and GFP-Shank3 (bottom row) images at selected time points (scale bar, 2 μm). Supplemental Movie 4 shows all time points. Summary quantitative data from all cells expressing FLAG-β2a are included in Figure 7C and 7D.

**Supplemental Figure 3 (related to Figure 9).**
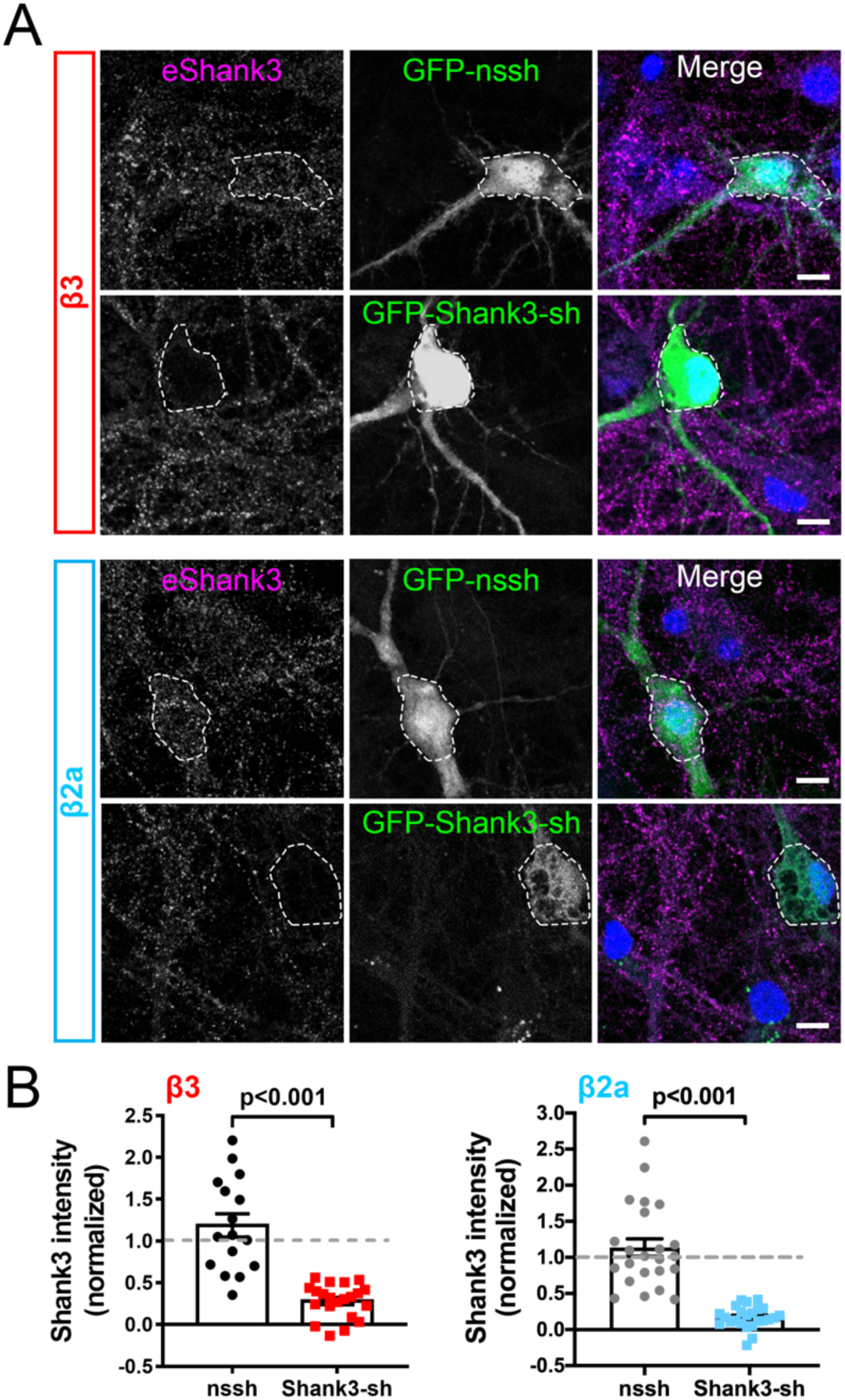
Validation of Shank3 shRNAs in cultured hippocampal neurons. Primary rat hippocampal neuron cultures (14 DIV) were transfected to express sHA-Ca_V_1.3, FLAG-β3 (top panels) or FLAG-β2a (bottom panels), and either GFP-nonsense shRNA (GFP-nssh) or GFP-Shank3 shRNA (GFP-Shank3-sh). Neurons were fixed at DIV21, permeabilized and then immunostained for endogenous Shank3. A series of GFP, DAPI and eShank3 images were collected at different optical planes using a confocal microscope. A) Representative maximum intensity projection of the Z-stack merged using FIJI. White dashed lines were used to outline GFP expression in soma. Scale bar, 10 μm. B) Quantification of total Shank3 staining intensity in the soma of cells expressing FLAG-β3 (left) or FLAG-β2a (right), normalized to Shank3 staining in nearby non-transfected cell somas on the same coverslips. Data collected from 3 independent transfections for each condition. β3: n = 16 and 21 neurons for GFP-nssh, and GFP-Shank3-sh, respectively. β2a: n = 23 and 24 neurons for GFP-nssh and. GFP-Shank3-sh, respectively. Comparisons made using an unpaired t-test.

**Supplemental Figure 4 (related to Figure 9).**
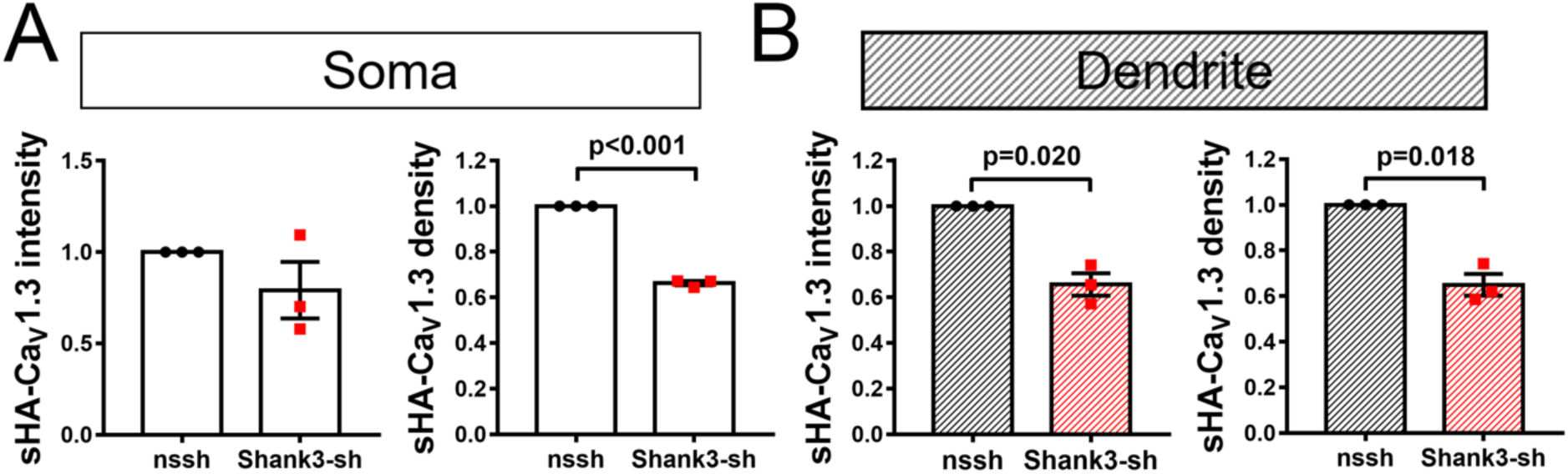
Quantification of surface localized sHA-Ca_V_1.3 cluster density and intensity on soma or dendrites in neurons co-expressing FLAG-β3. Data from Figure 9 were replotted by averaging the cluster intensity or density within each independent replicate and normalizing to values in cells expressing GFP-nssh (n=3 cultures/transfections, 9-16 cells per transfection). Comparisons made using a one-sample t-test.

**Supplemental Figure 5 (related to Figure 9).**
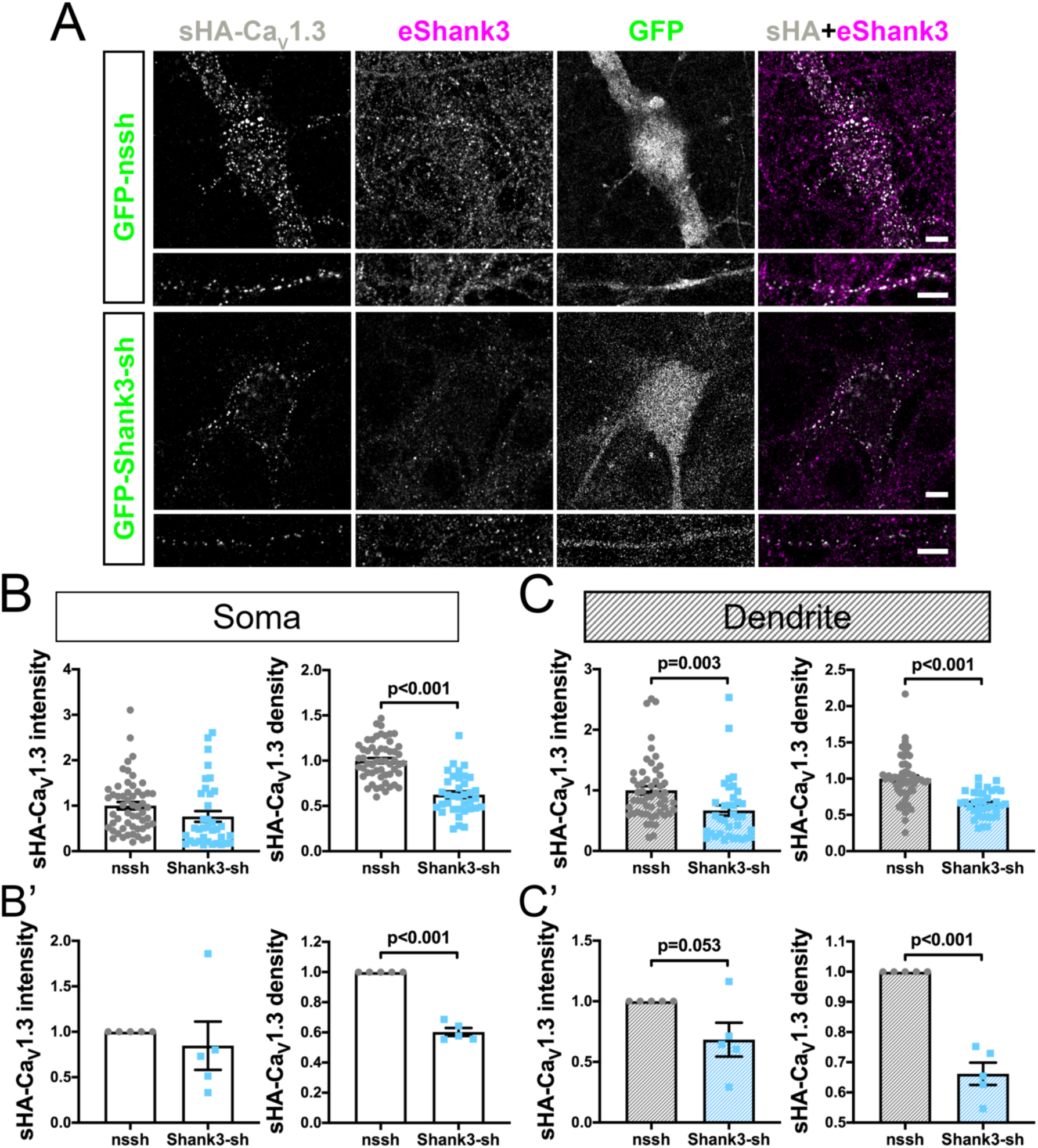
Effects of Shank3 knock-down on surface-expressed Ca_V_1.3 puncta in neurons expressing FLAG-β2a. These experiments were conducted in parallel with those shown in Fig. 9, except that FLAG-β2a was co-expressed instead of FLAG-β3. A) Representative images of soma and dendrites. Scale bar, 5 μm. B) and C) Quantification of sHA-Ca_V_1.3 cluster intensity and cluster density from n = 55 (GFP-nssh) or 36 (GFP-Shank3-sh) neurons from five independent cultures/transfections; comparisons made using an unpaired t-test. B’) and C’) Re-plot of the same data after averaging the cluster intensity or density within each independent replicate and normalizing to values in cells expressing the control shRNA (n=5). Comparisons made using a one-sample t-test.

**Supplemental Figure 6 (related to Figure 10).**
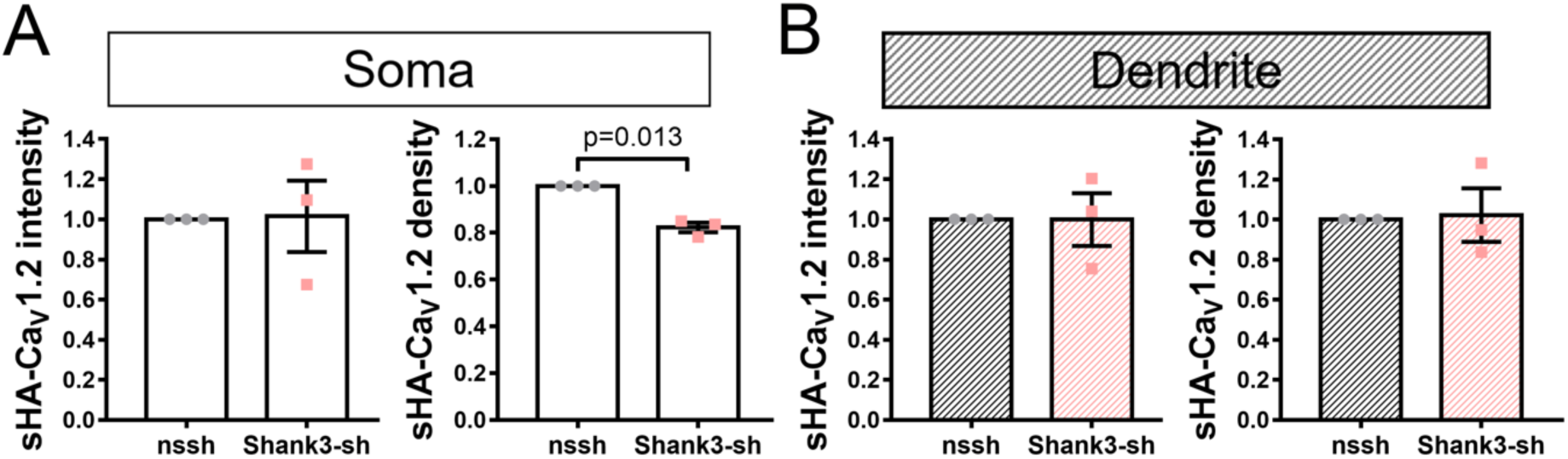
Quantification of cluster density and intensity of surface localized sHA-Ca_V_1.2 channels on soma or in dendrites in neuron co-expressing FLAG-β3. Data from Figure 10 were replotted by averaging the cluster intensity or density within each independent replicate and normalizing to values in neurons expressing the control shRNA (n=3 cultures/transfections, 7-10 cells per transfection). Comparisons made using a one-sample t-test.

Supplemental Movie 1 (related to Figure 6). Live cell imaging of a representative HEK293 cell expressing mCherry-Ca_V_1.3, FLAG-β3 and GFP under basal conditions.

Supplemental Movie 2 (related to Figure 6). Live cell imaging of a representative HEK293 cell expressing mCherry-Ca_V_1.3, FLAG-β3 and GFP-Shank3 under basal conditions.

Supplemental Movie 3 (related to Figure 7). Live cell imaging of a representative HEK293 cell expressing mCherry-Ca_V_1.3, FLAG-β3 and GFP-Shank3. Cell was imaged in “no Ca^2+^” buffer, following the addition of BayK 8644 (10 µM), and following the further addition of Ca^2+^ (2.5 mM CaCl_2_).

Supplemental Movie 4 (related to Supplemental Figure 2). Live cell imaging of a representative HEK293 cell expressing mCherry-Ca_V_1.3, FLAG-β2a and GFP-Shank3. Cell was imaged in “no Ca^2+^” buffer, following the addition of BayK 8644 (10 µM), and following the further addition of Ca^2+^ (2.5 mM CaCl_2_).

**Supplementary Table 1.** The Prism output for the statistical analyses of data shown in the figures.

**Figure 2B.**
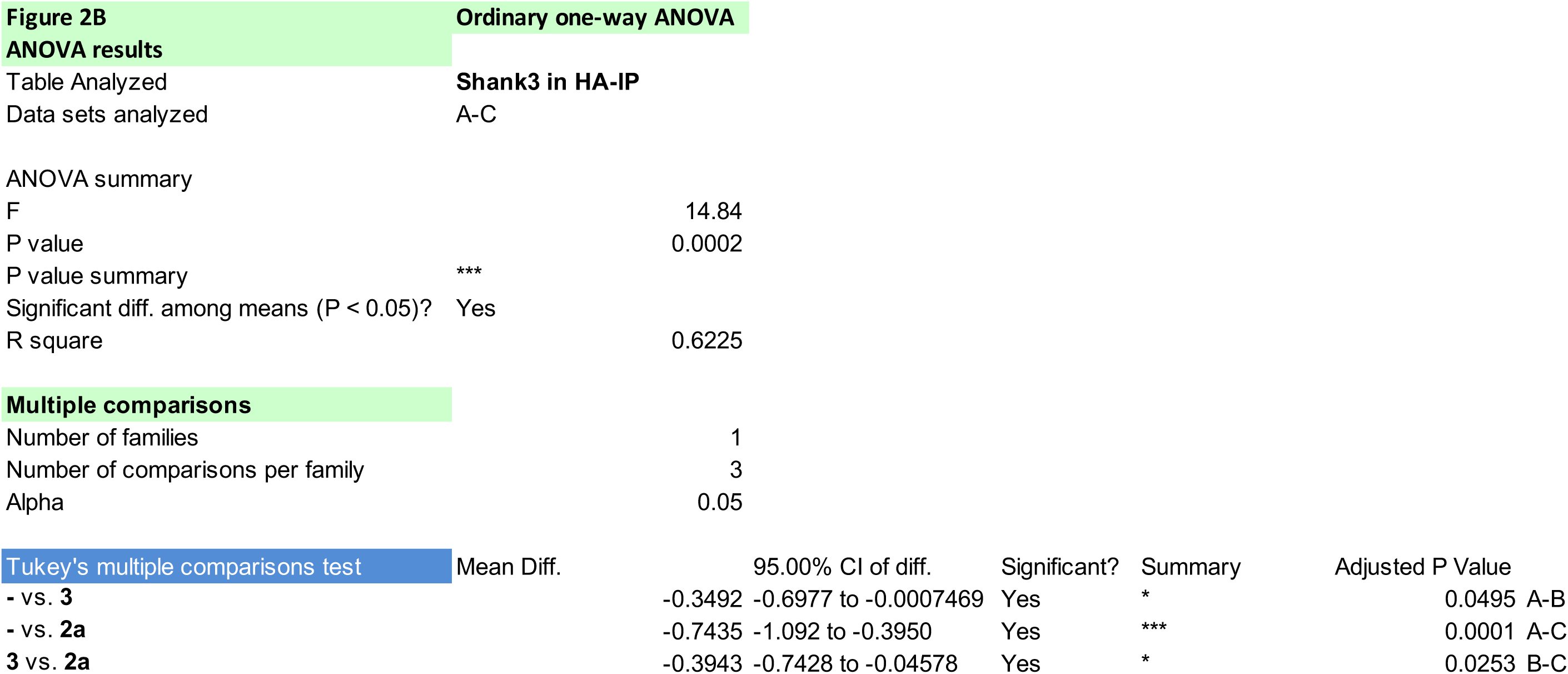

**Figure 2C.**
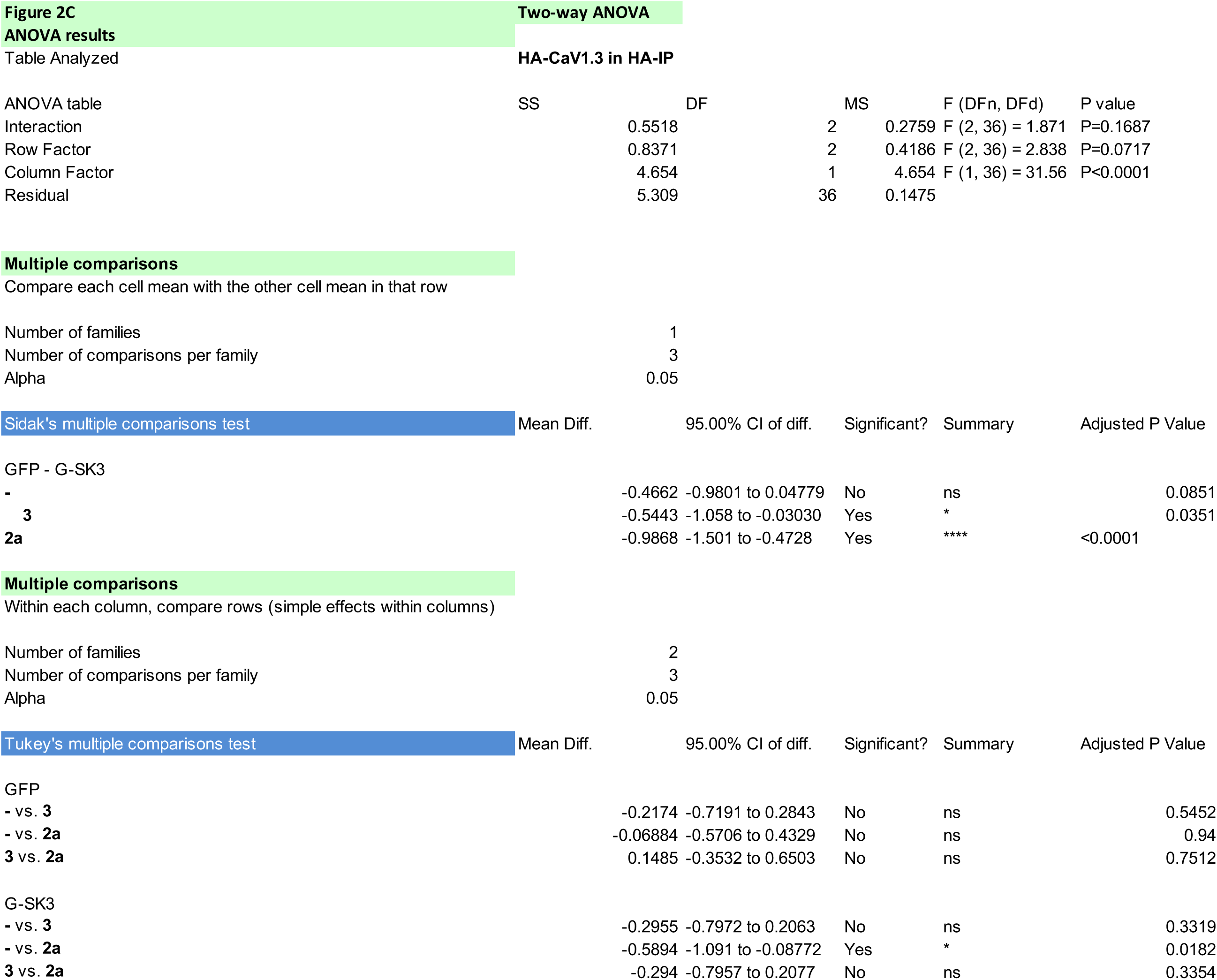

**Figure 2E.**
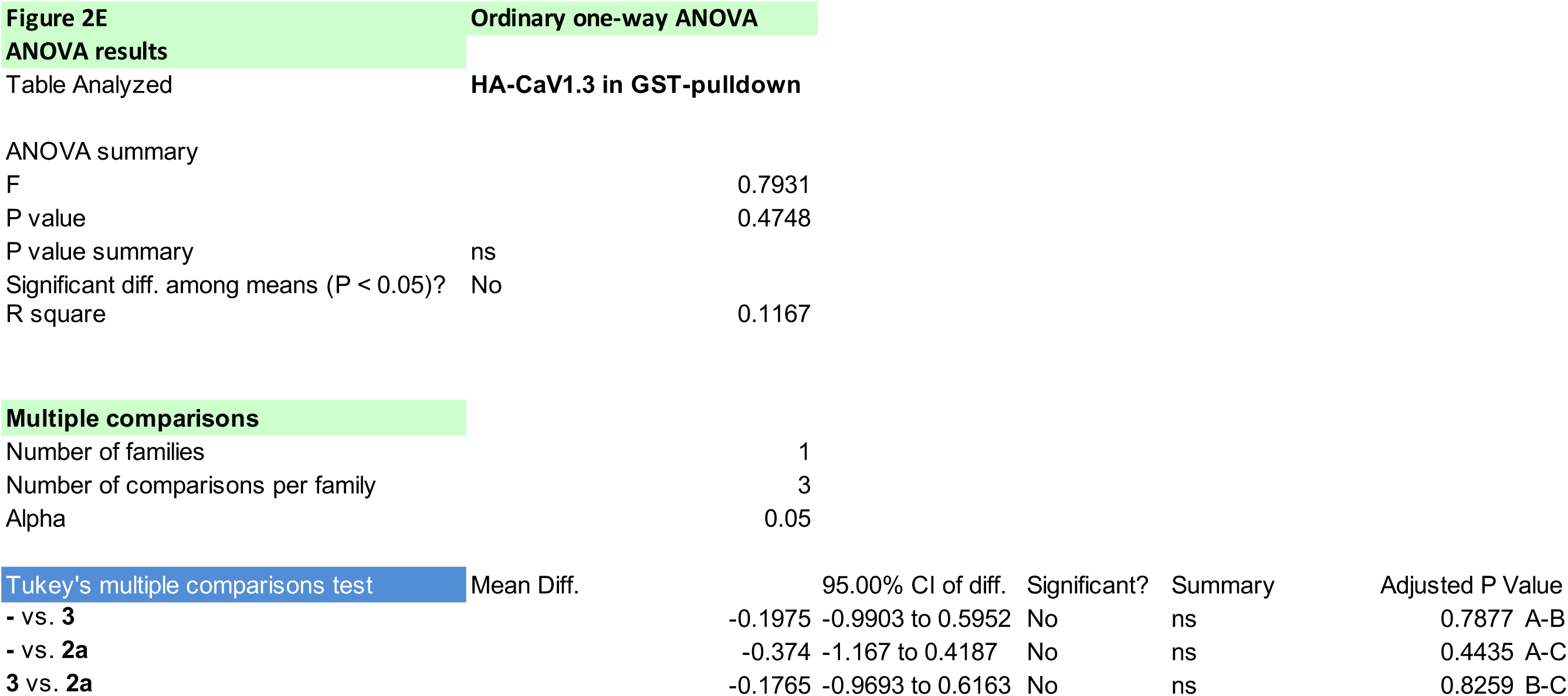

**Figure 3B.**
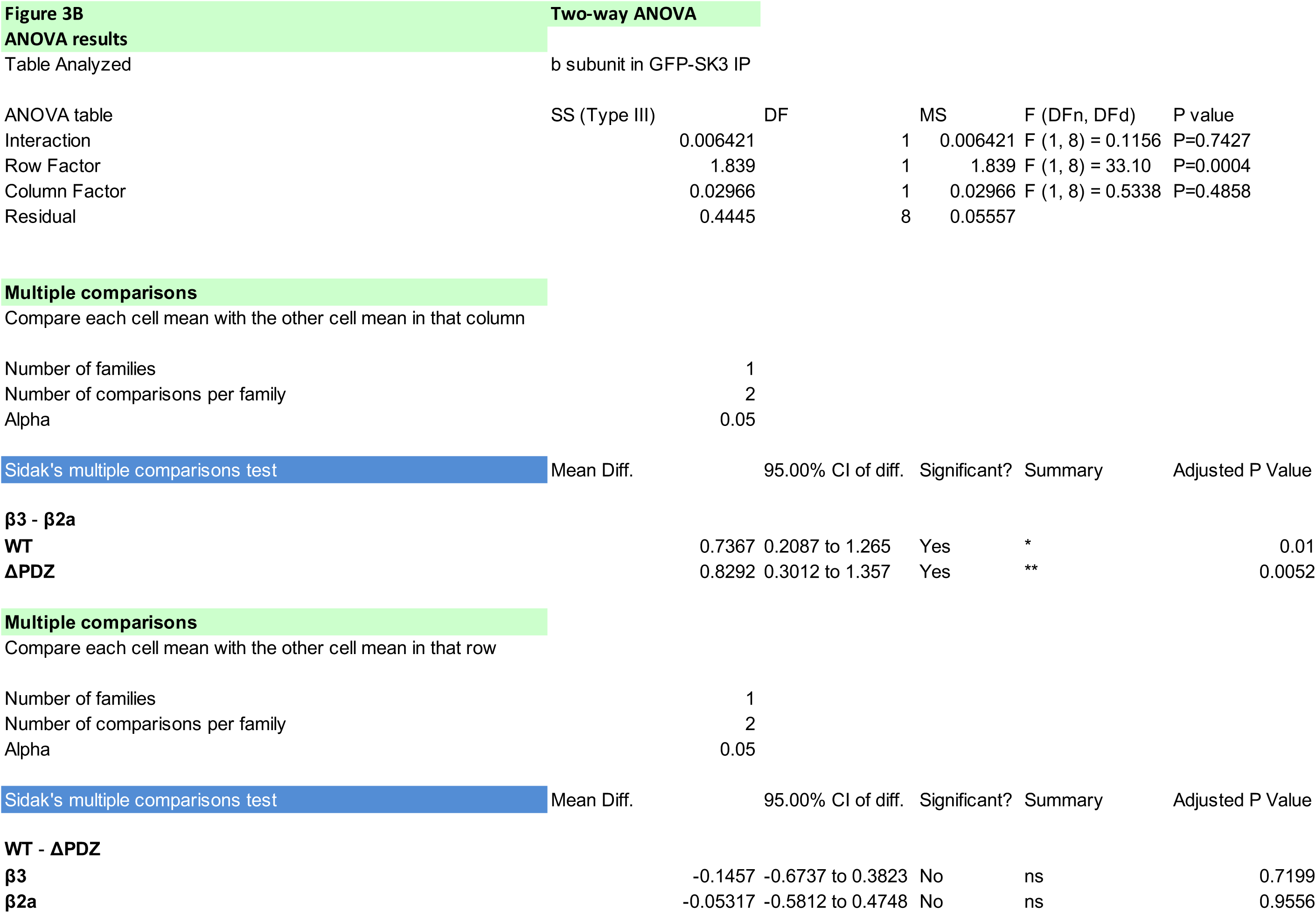

**Figure 3D.**
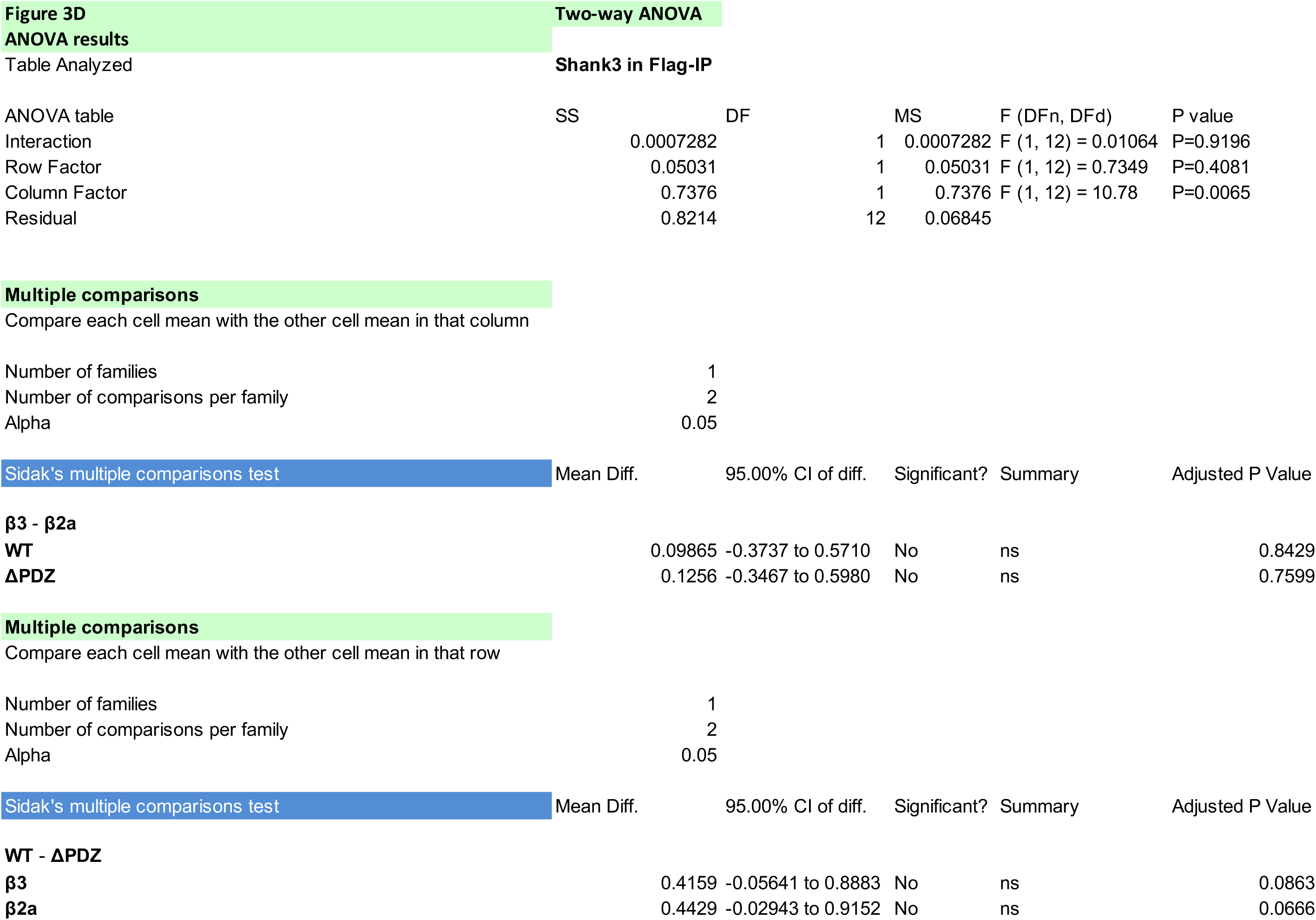

**Figure 4B.**
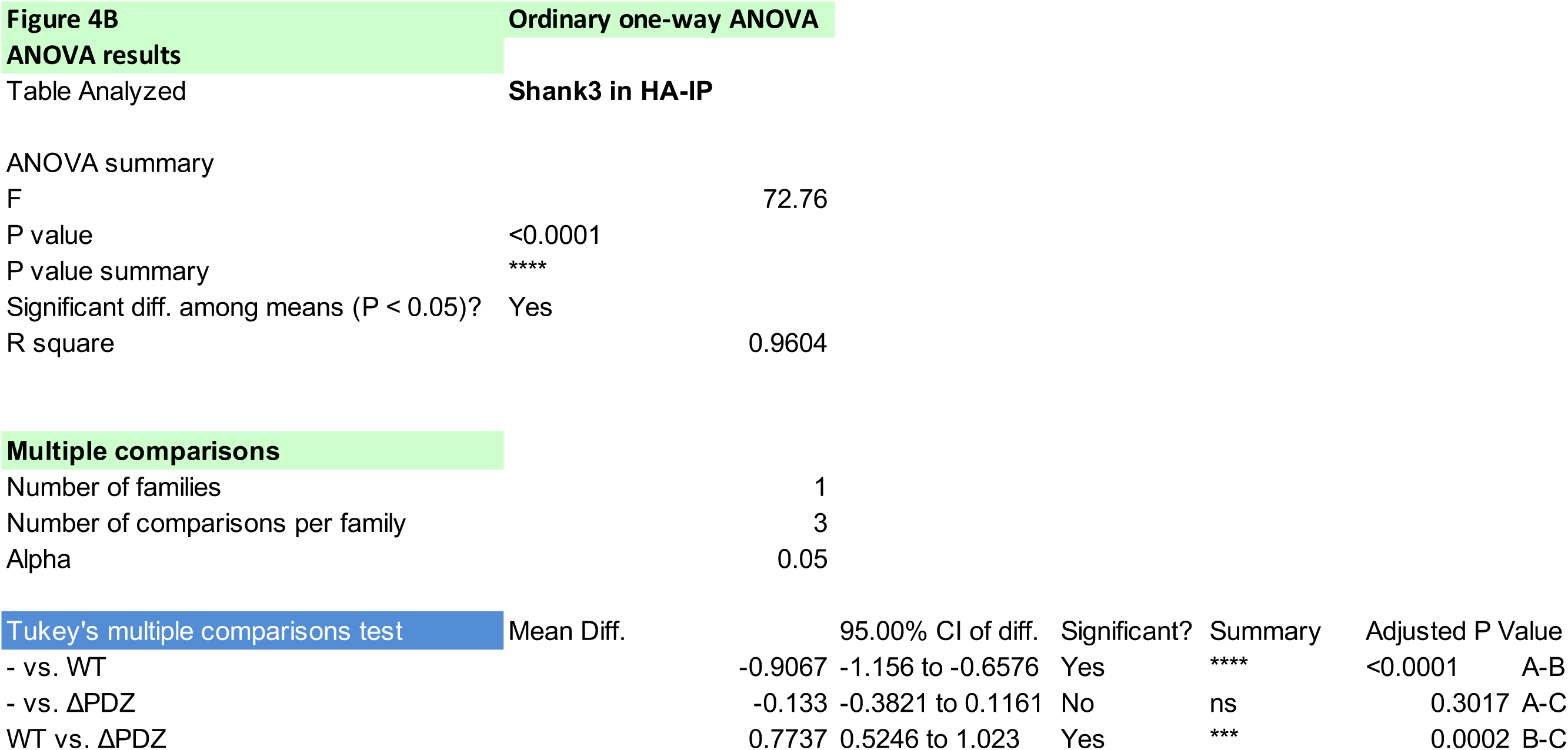

**Figure 4C.**
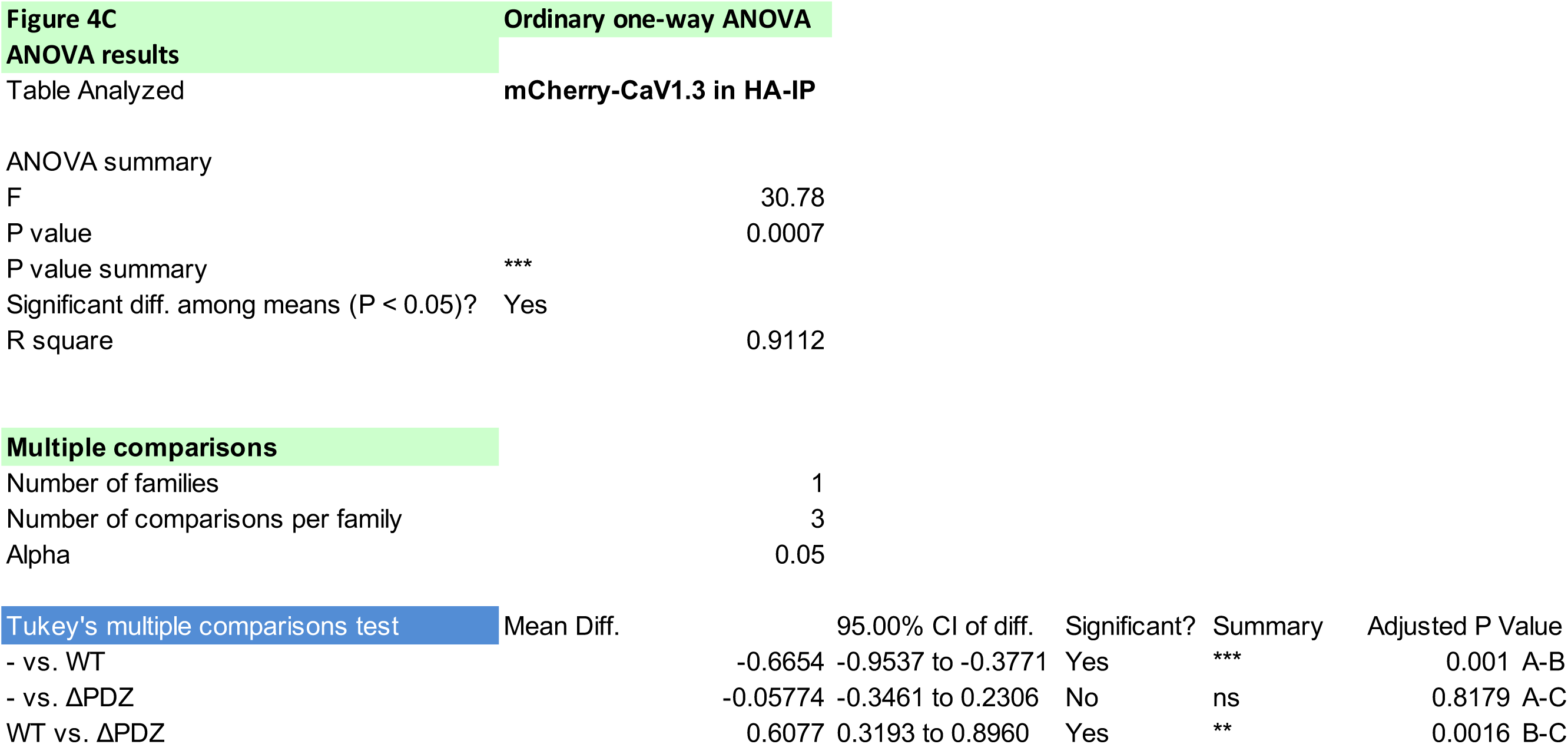

**Figure 5B.**
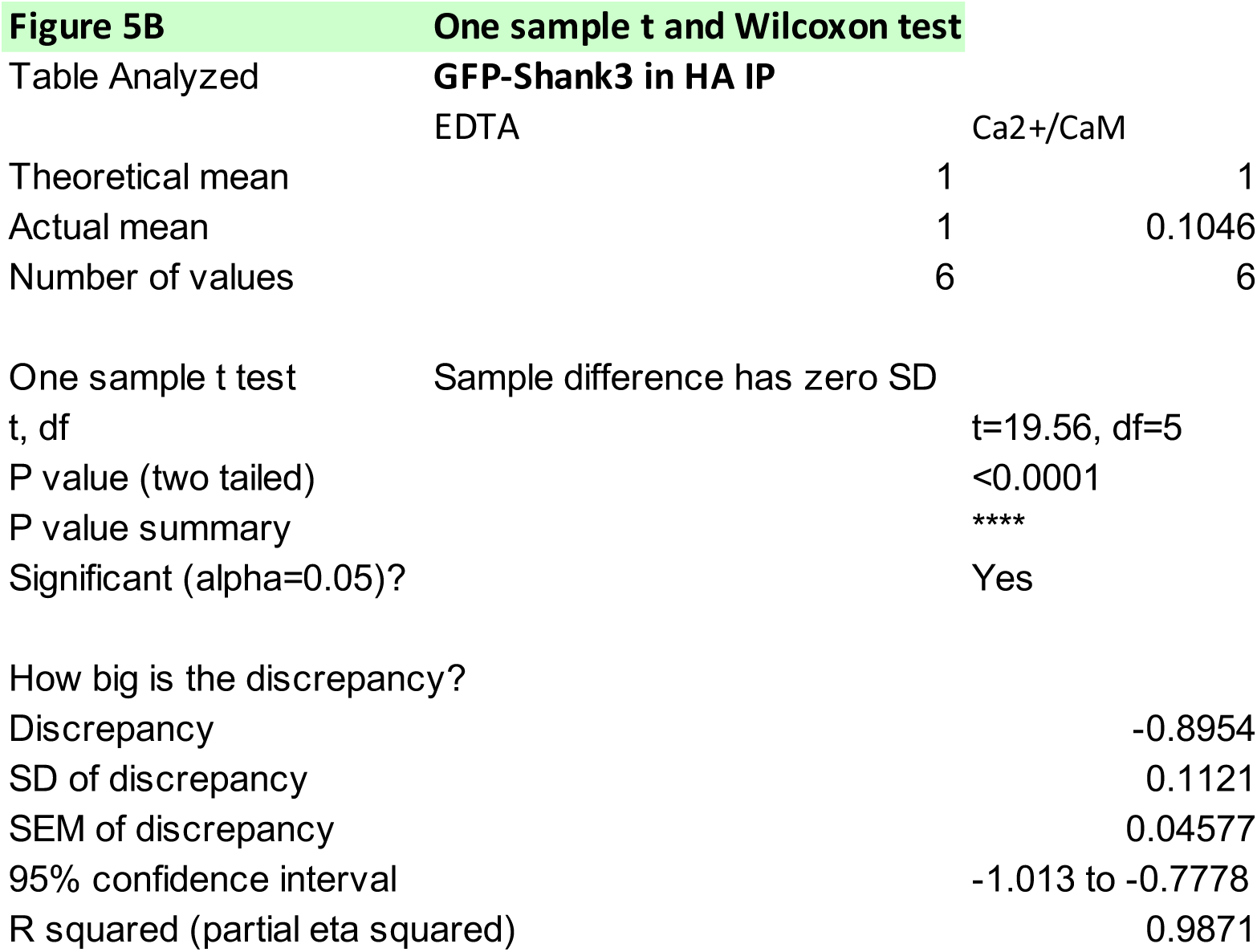

**Figure 5C.**
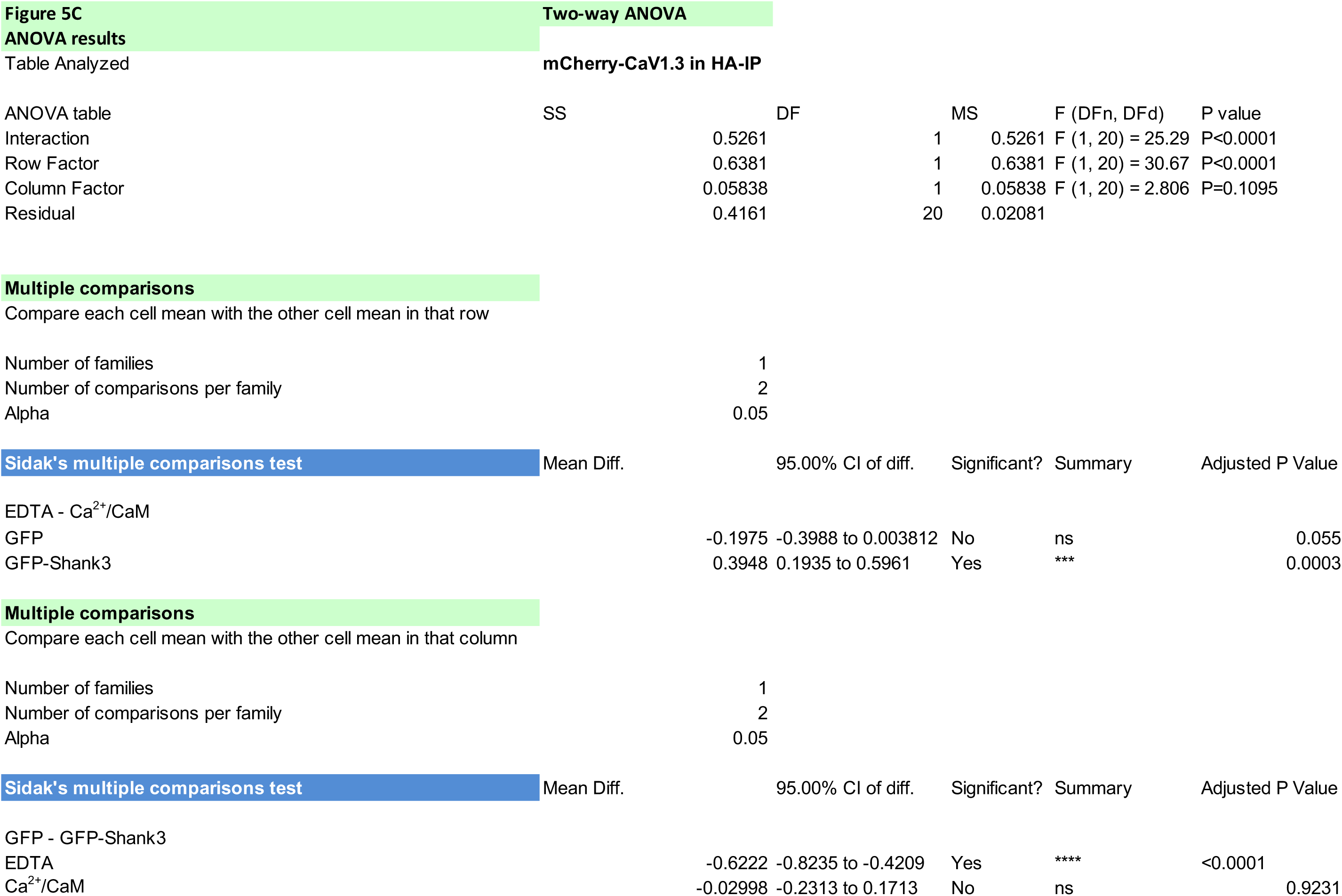

**Figure 6C.**
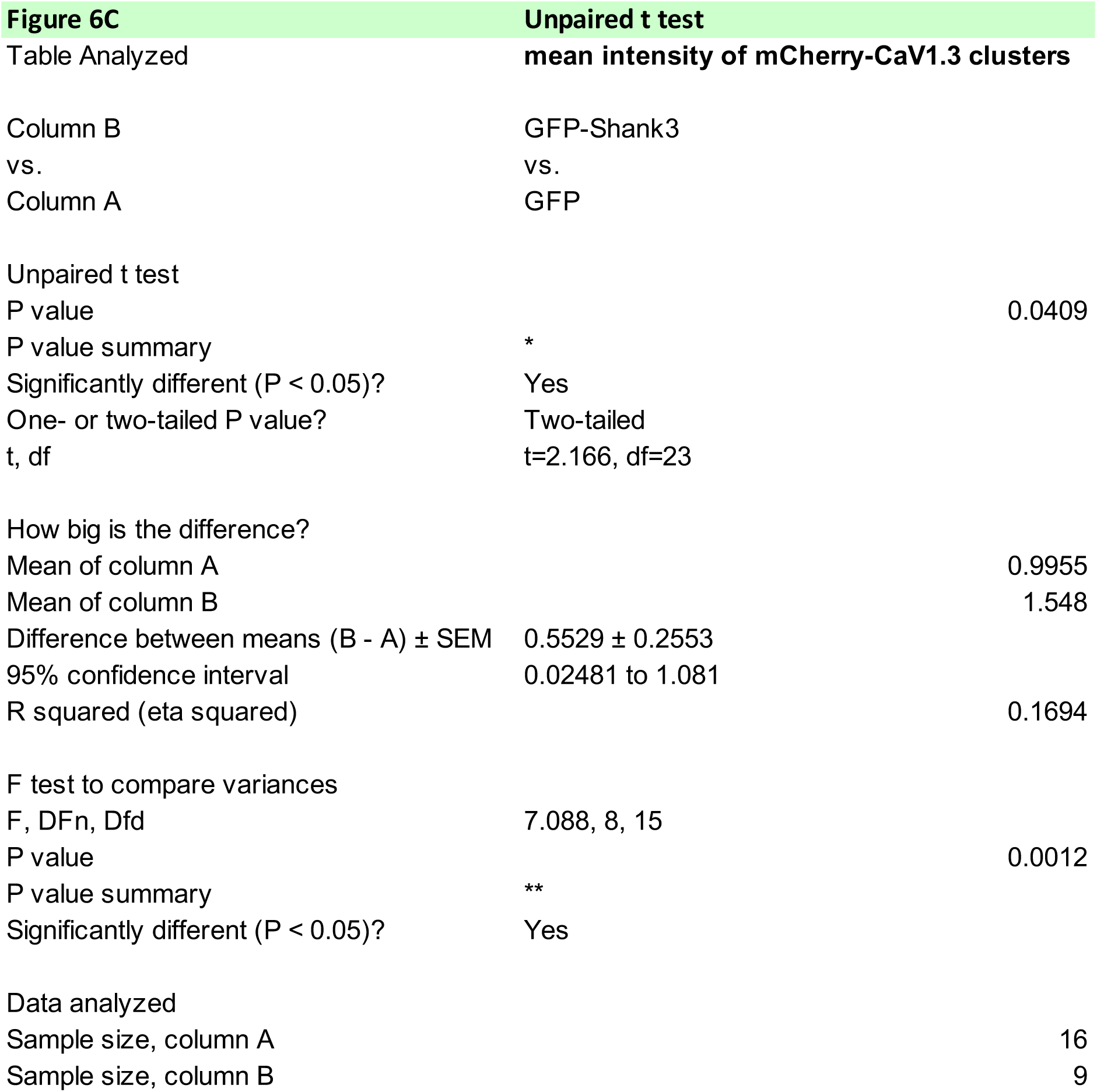

**Figure 6D.**
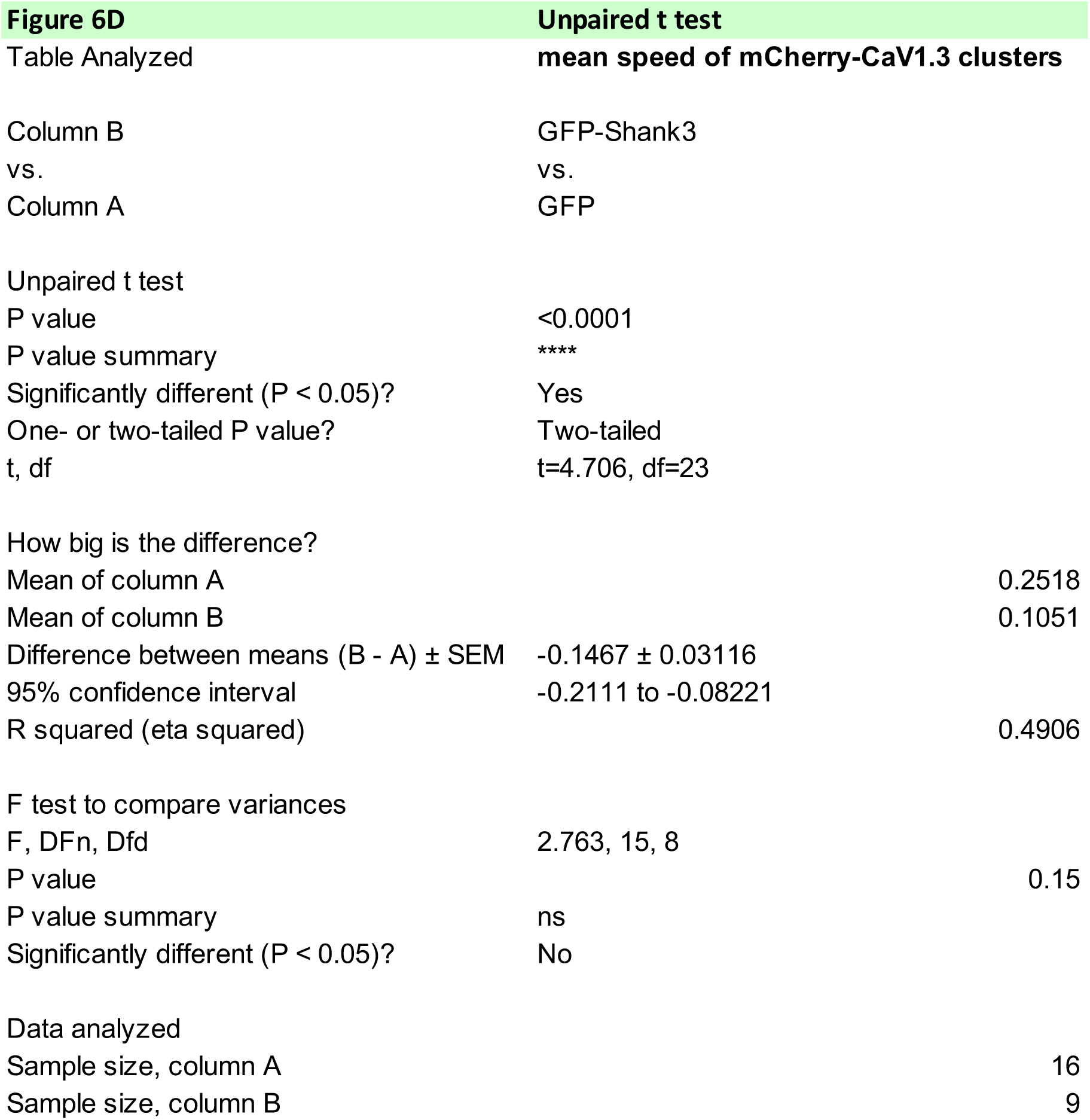

**Figure 7C.**
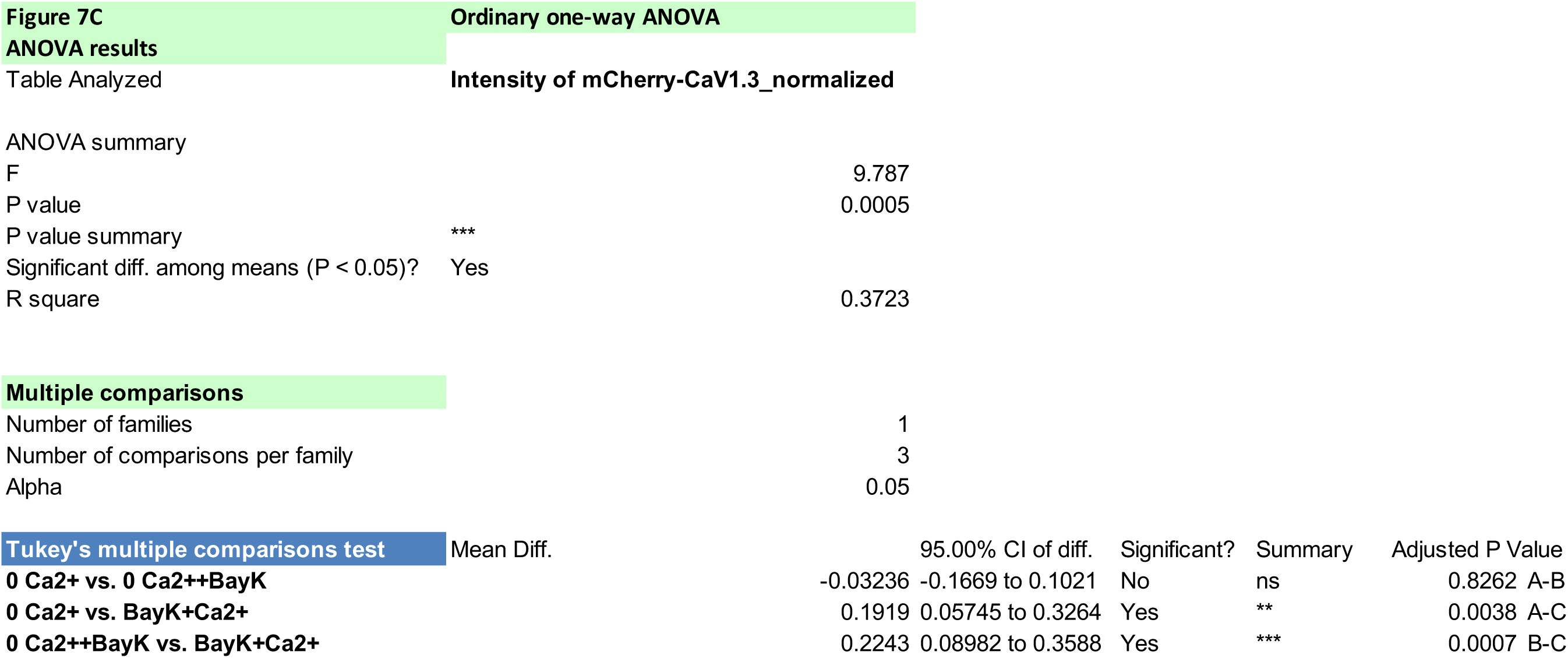

**Figure 7D.**
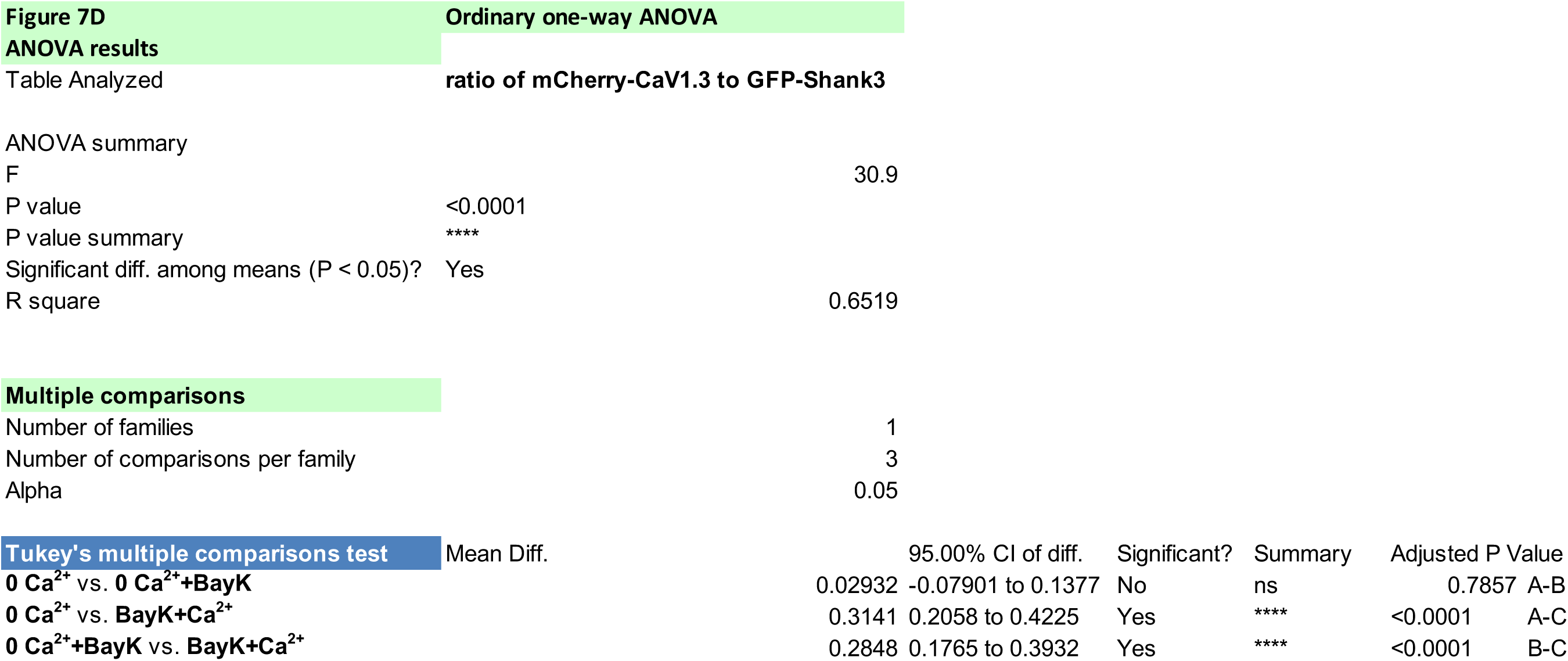

**Figure 8B.**
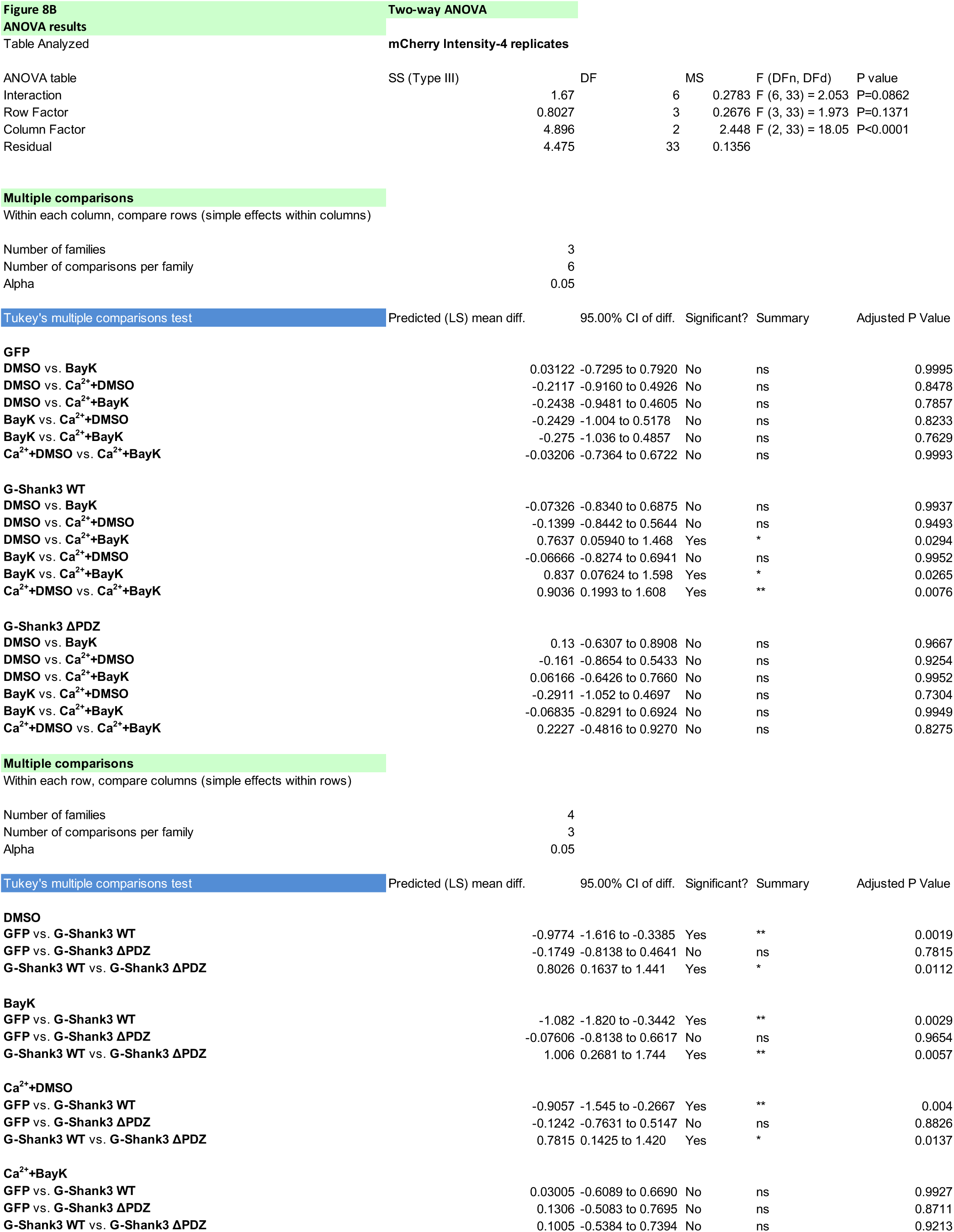

**Figure 8C.**
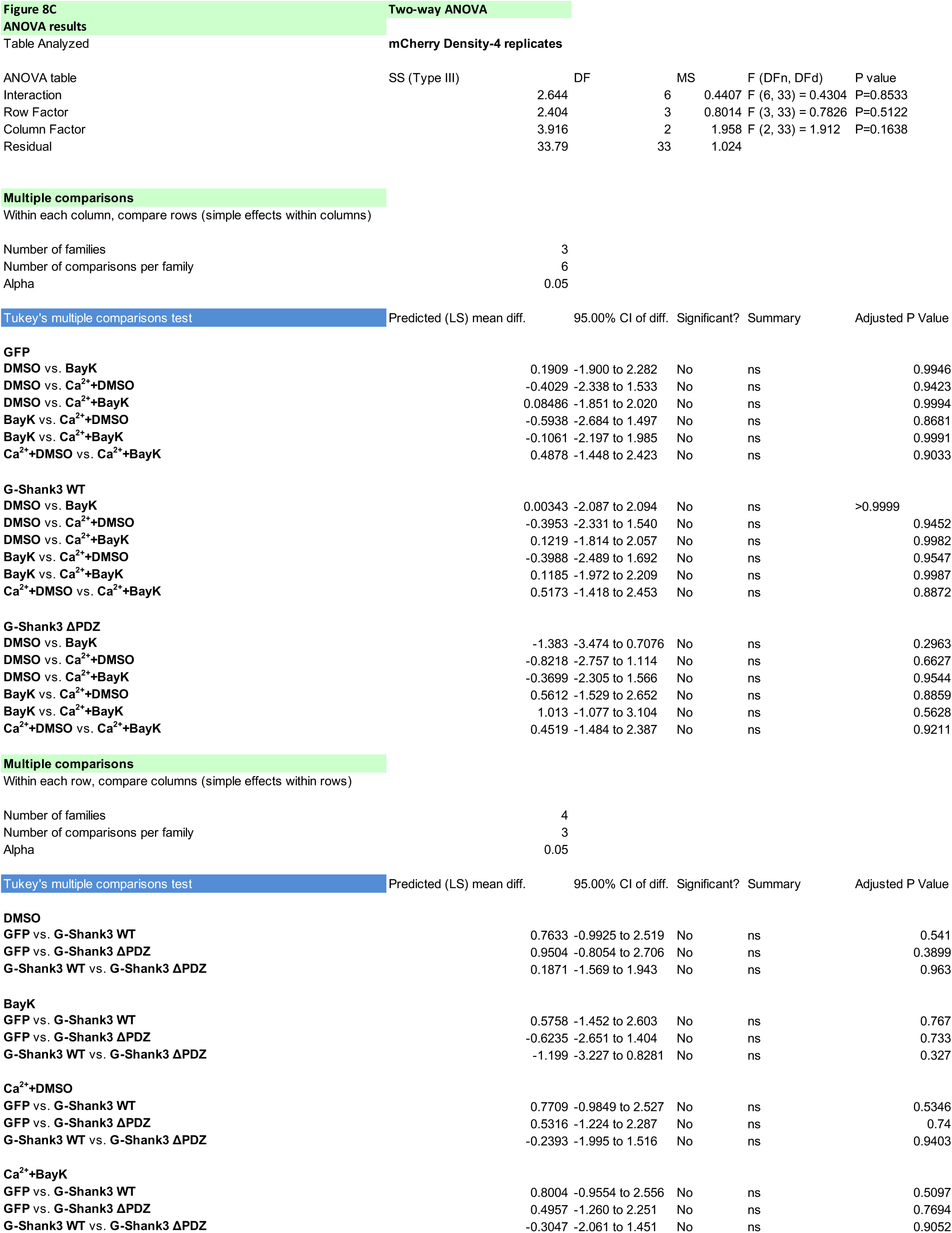

**Figure 8D.**
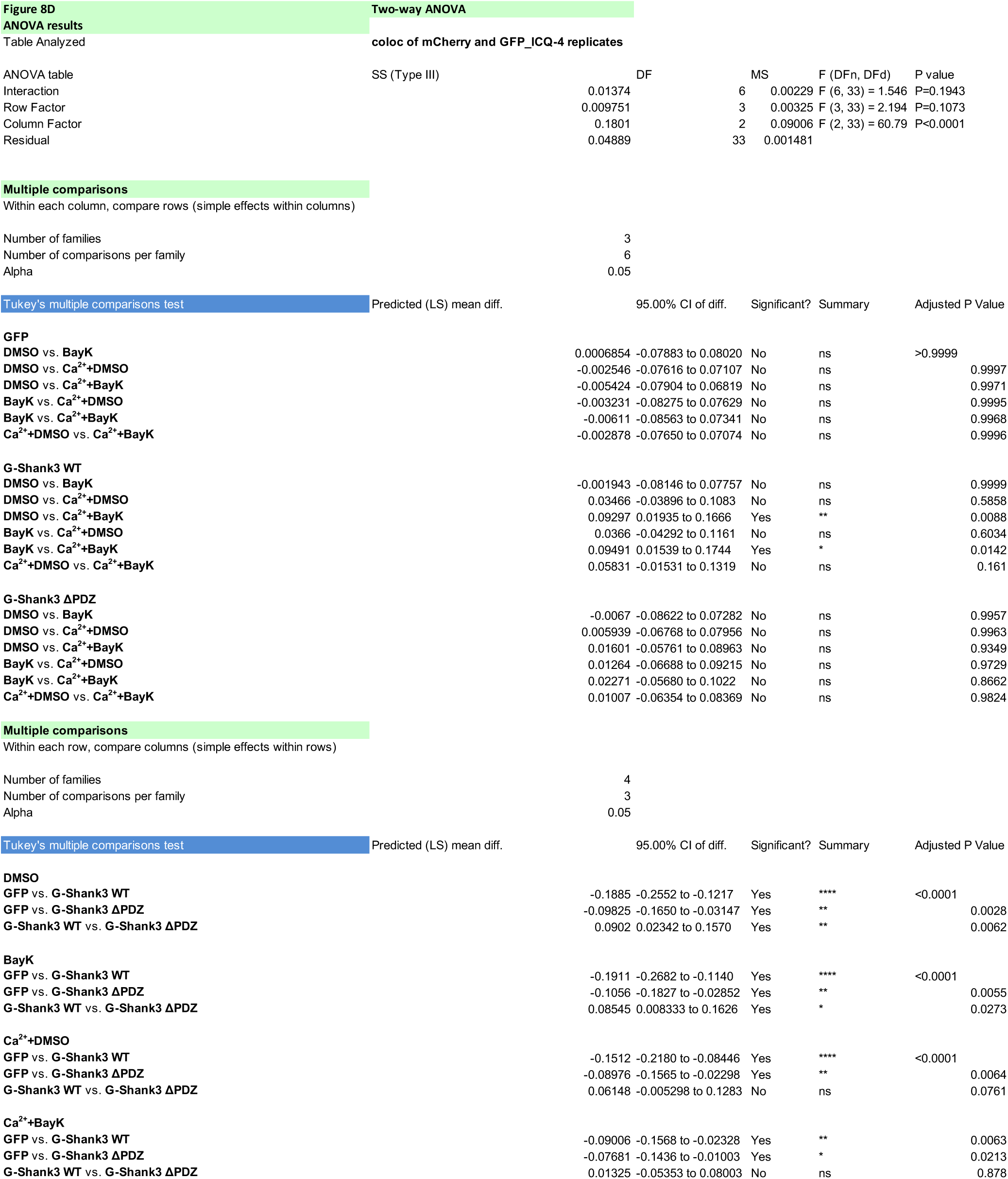

**Figure 9B.**
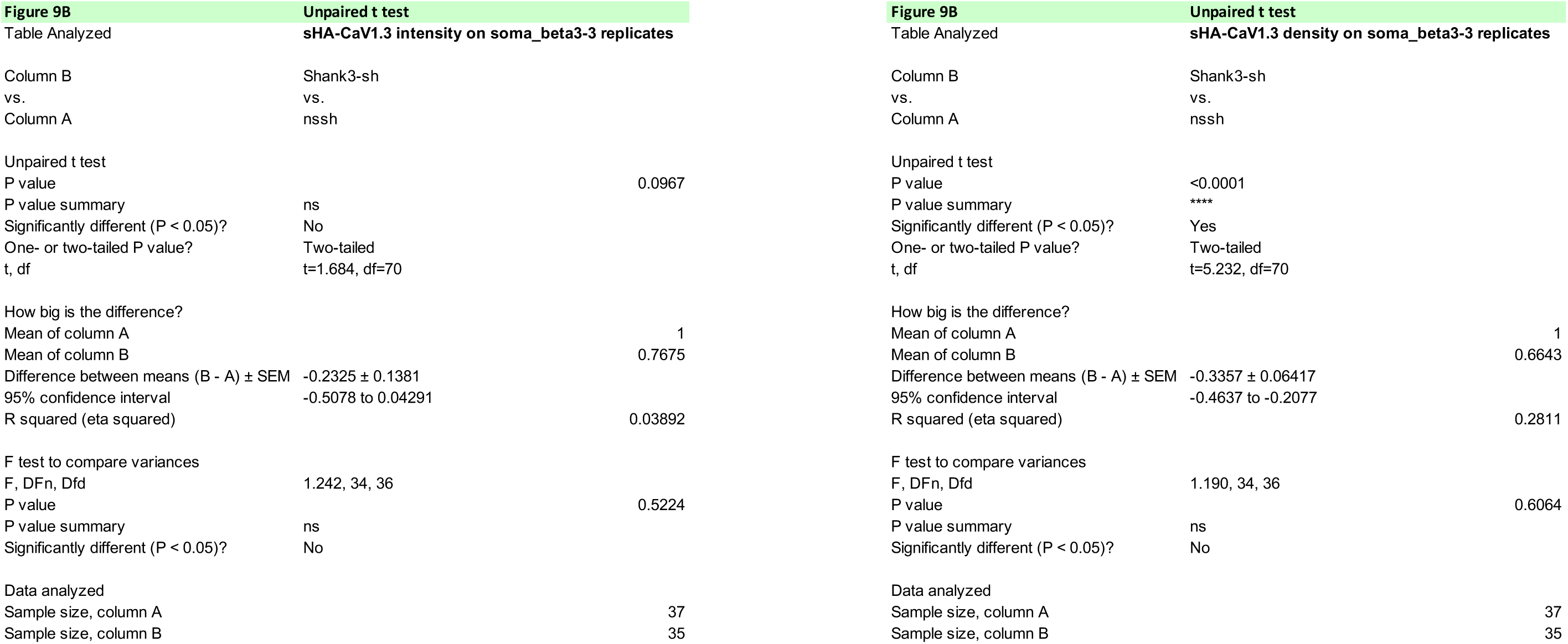

**Figure 9C.**
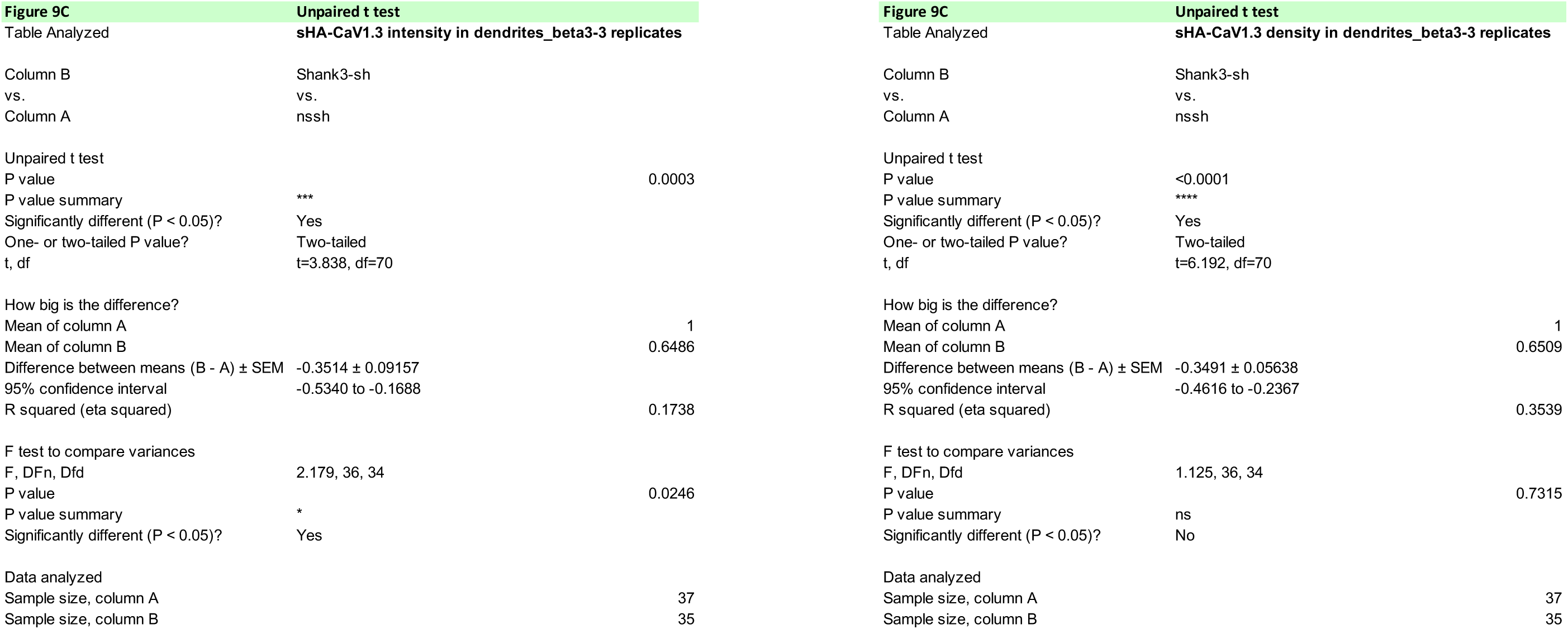

**Figure 10B.**
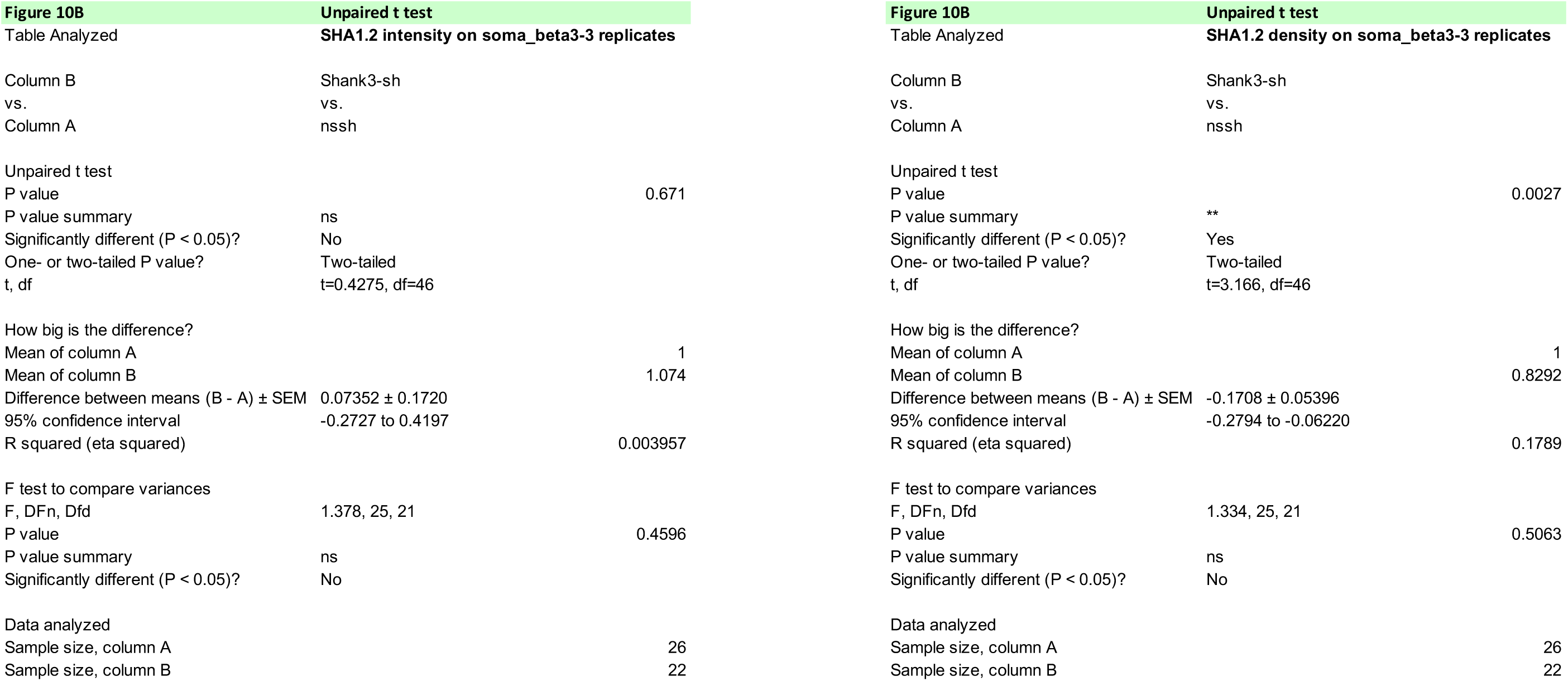

**Figure 10C.**
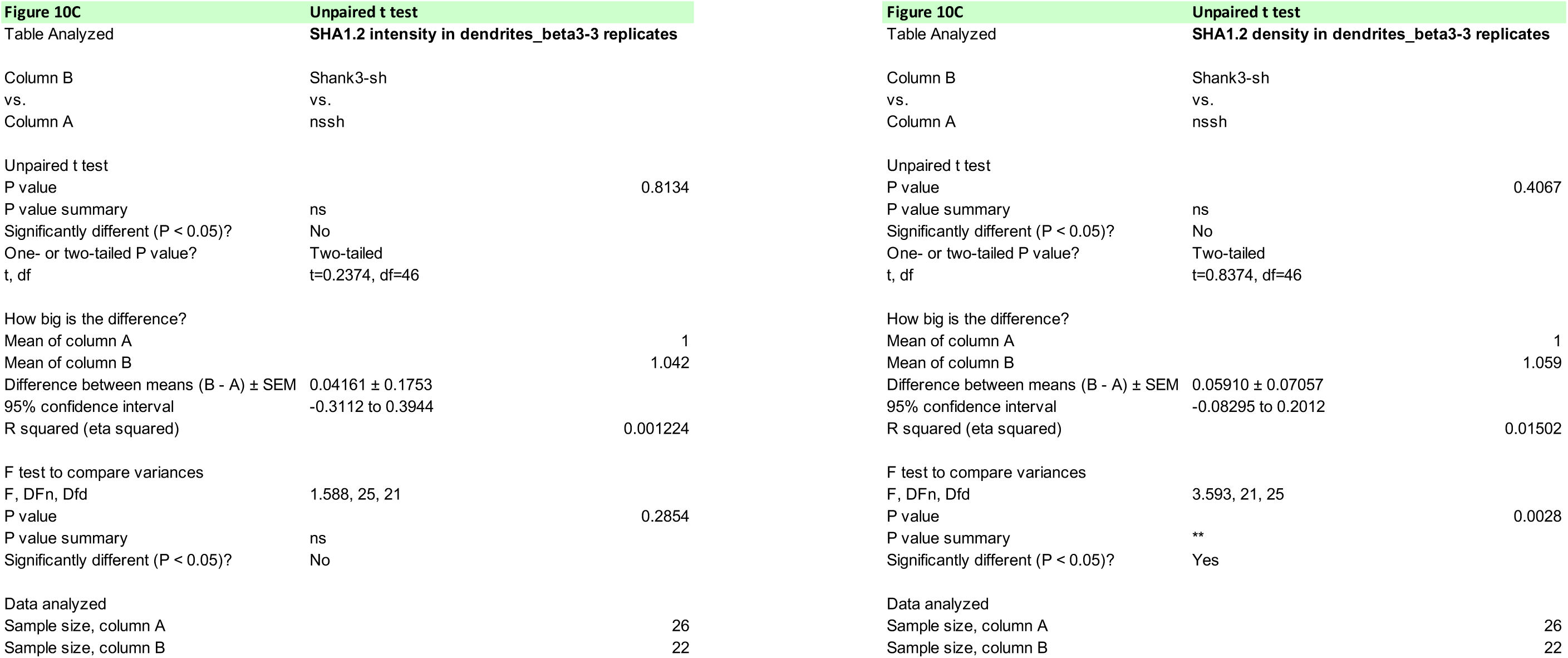

**S Figure 3B.**
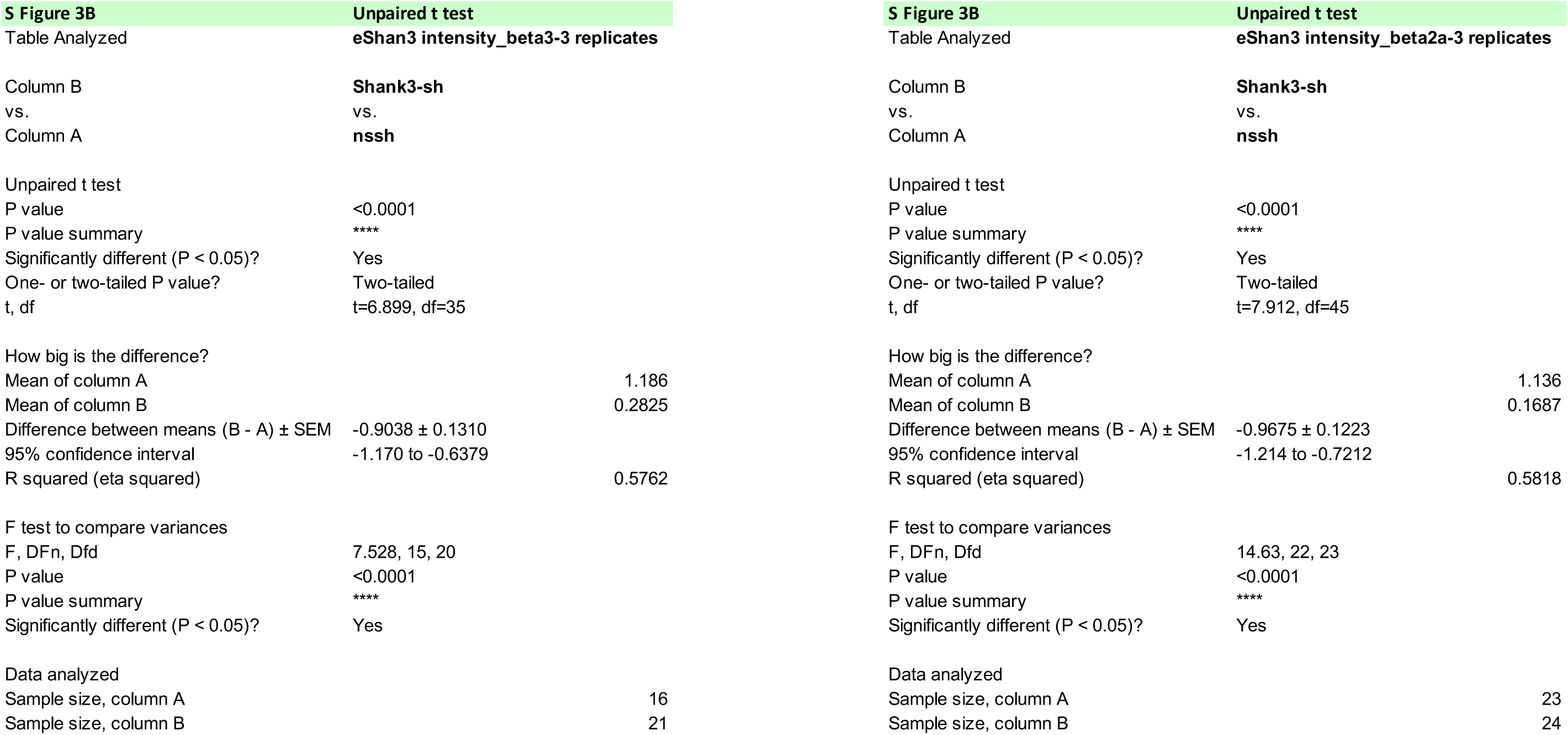

**S Figure 4A.**
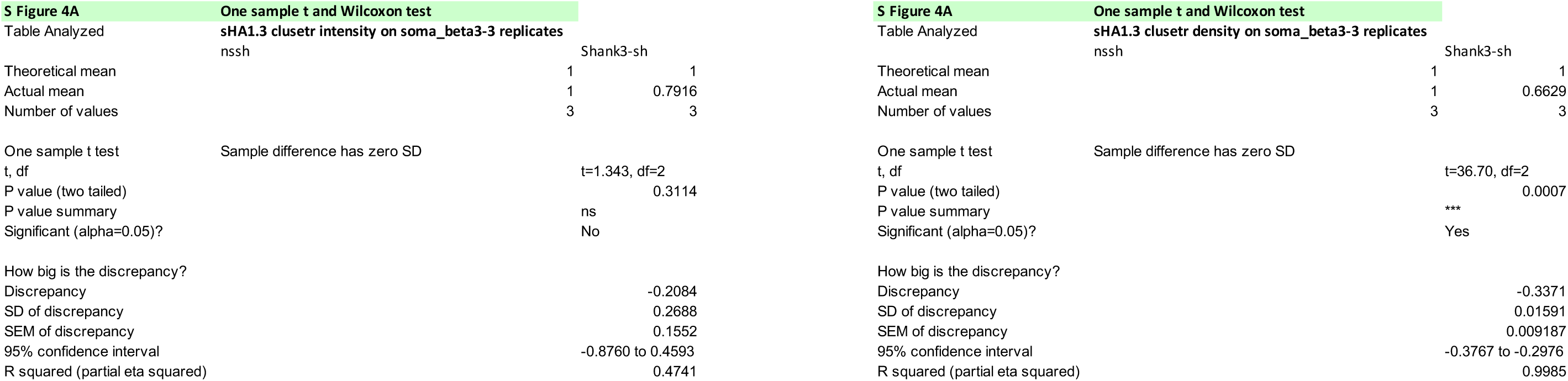

**S Figure 4B.**
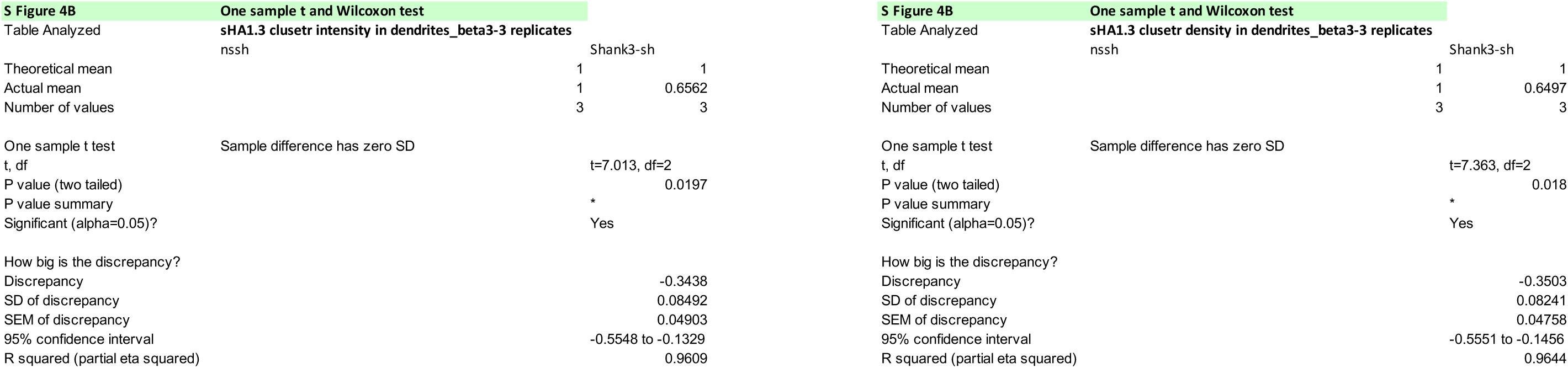

**S Figure 5B.**
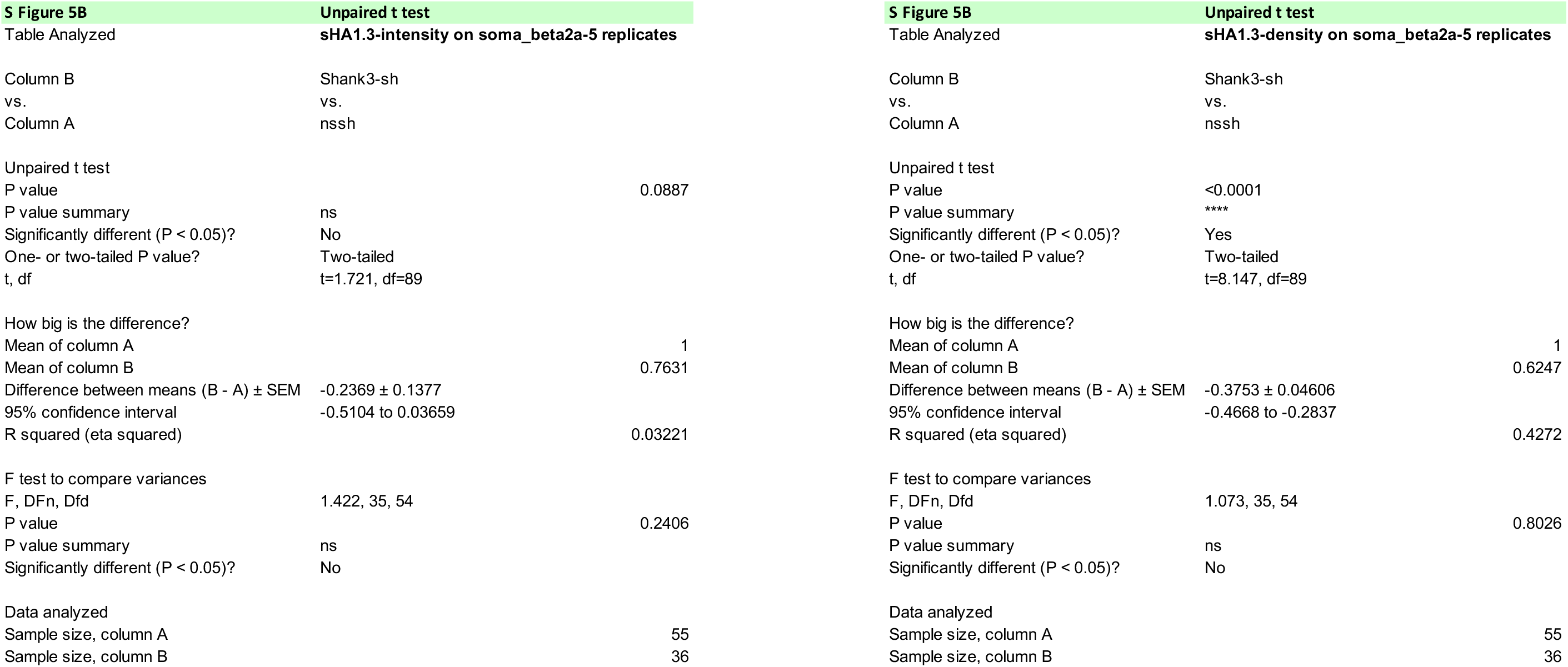

**S Figure 5B”.**
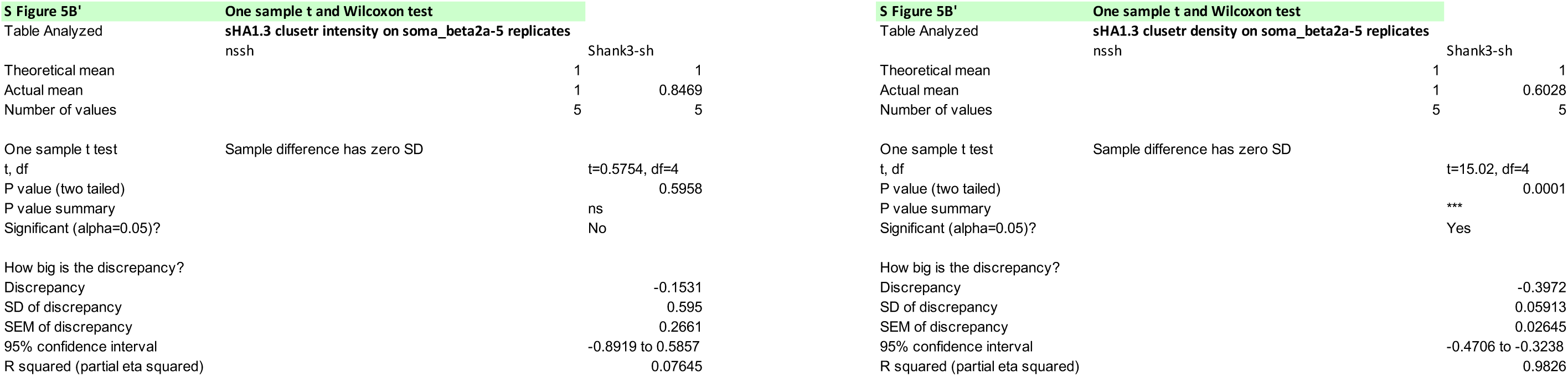

**S Figure 5C.**
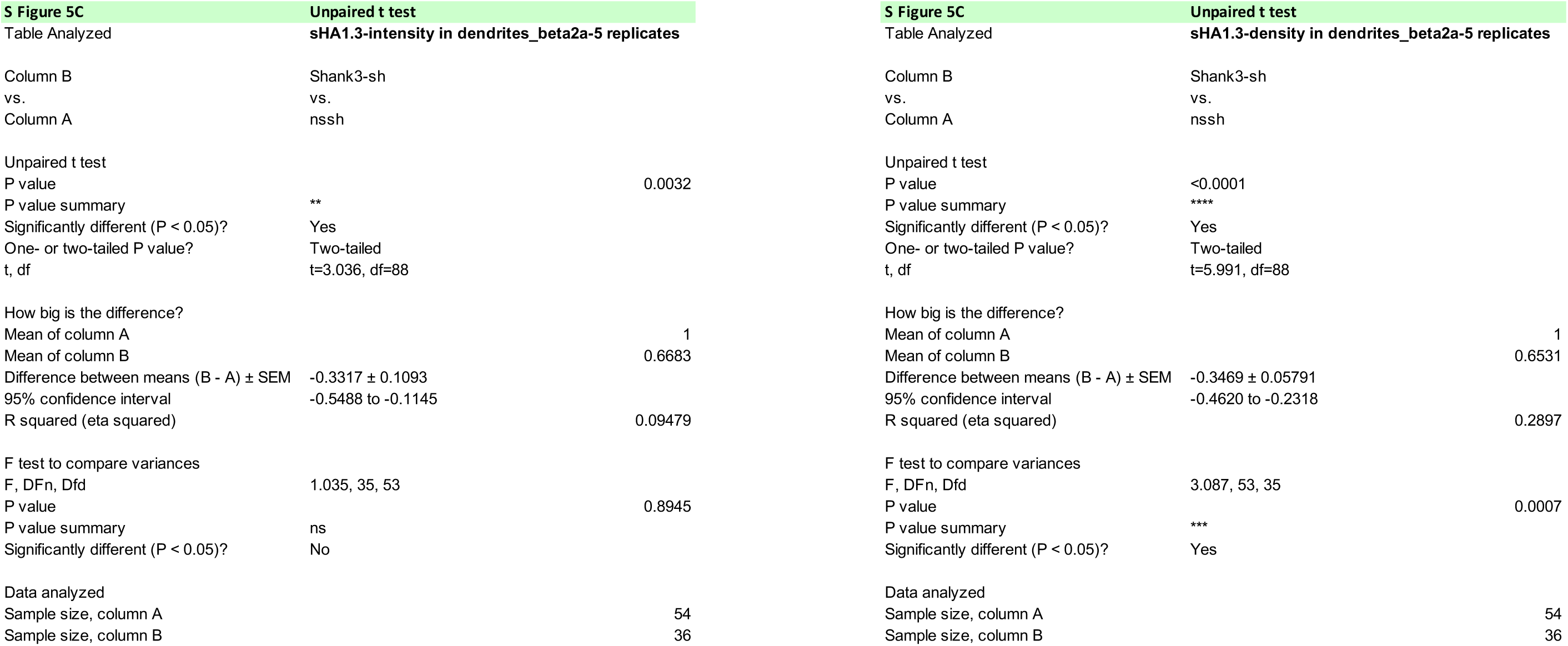

**S Figure 5C”.**
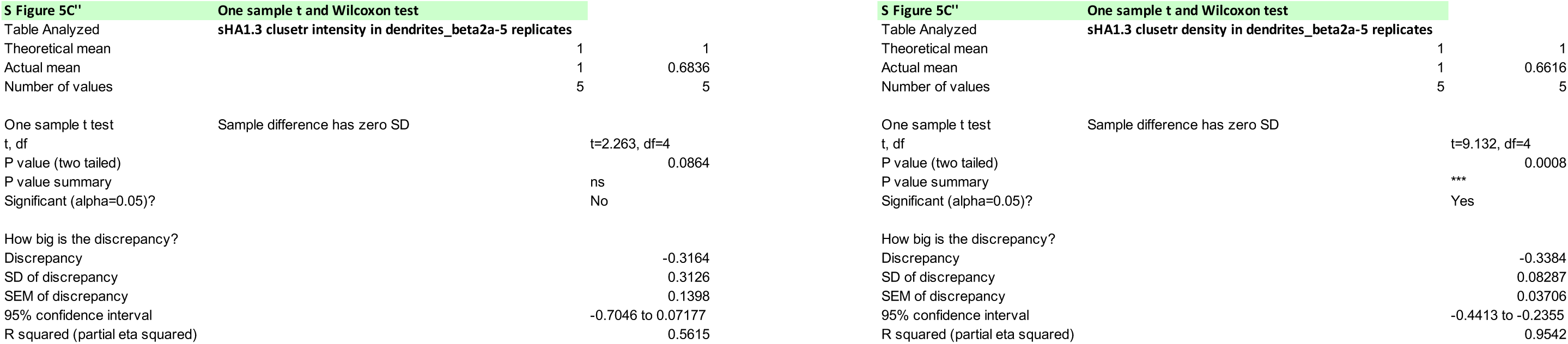

**S Figure 6A.**
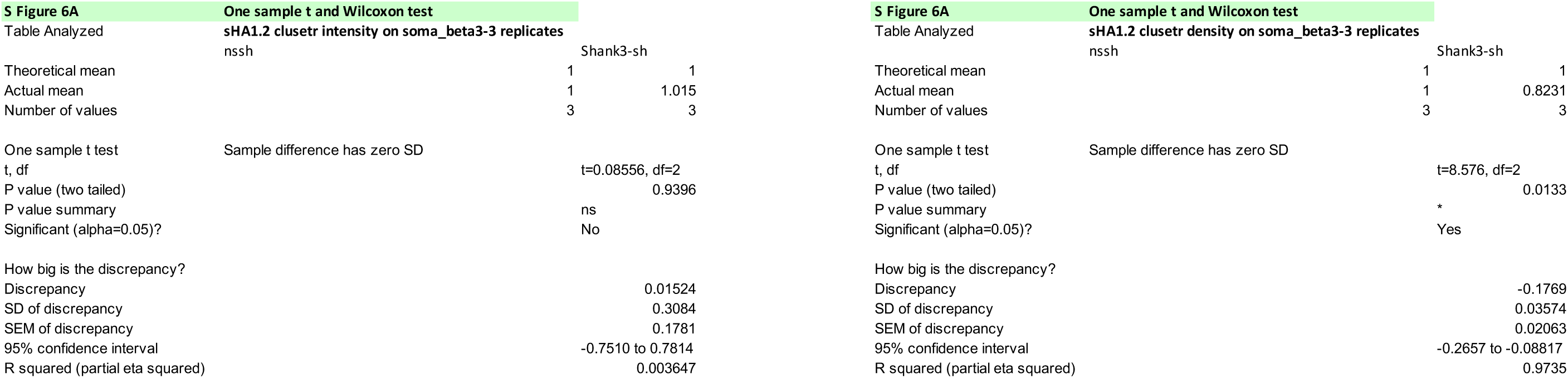

**S Figure 6B.**
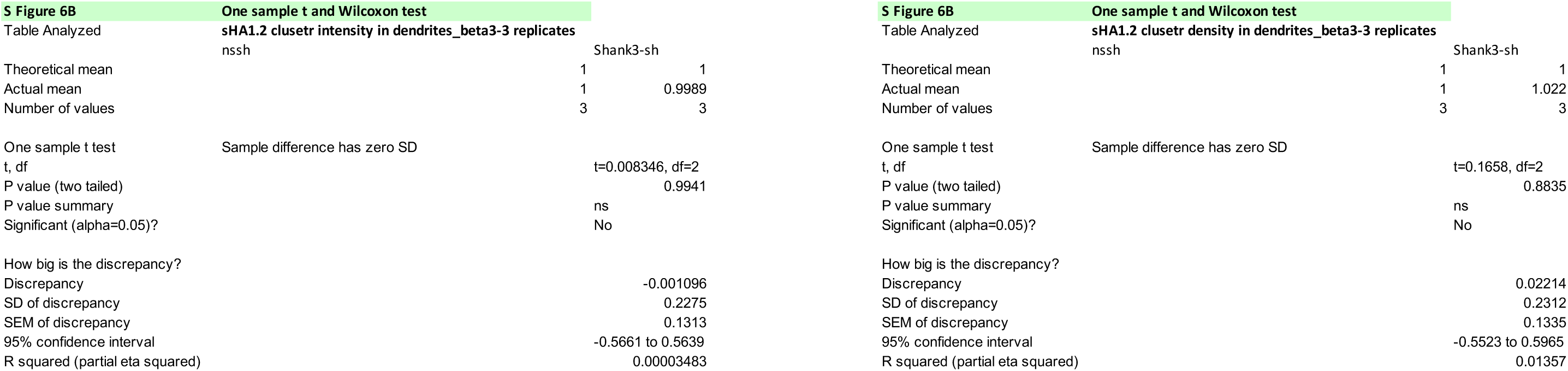

## Notes

### Competing Interest Statement

The authors have declared no competing interest.

